# Epigenetic inheritance mediates phenotypic diversity in natural populations

**DOI:** 10.1101/2021.03.15.435374

**Authors:** Zaigham Shahzad, Jonathan D. Moore, Jaemyung Choi, Gaëlle Cassin-Ross, Hatem Rouached, Daniel Zilberman

**Affiliations:** Department of Cell & Developmental Biology, John Innes Centre, Norwich, NR4 7UH, UK; Department of Life Sciences, Syed Babar Ali School of Science and Engineering, Lahore, Pakistan; Institute of Science and Technology, 3400 Klosterneuburg, Austria; Plant Resilience Institute, Michigan State University, East Lansing, MI 48824, USA; Department of Plant, Soil, and Microbial Sciences, Michigan State University, USA

## Abstract

Genetic variation is regarded as a prerequisite for evolution. In principle, epigenetic information inherited independently of DNA sequence can also enable evolution, but whether this occurs in natural populations is unknown. Here we show that natural epigenetic DNA methylation polymorphism in *Arabidopsis thaliana* gene bodies regulates gene expression and influences the variation of complex traits: fitness under heat and drought, flowering time, and accumulation of diverse minerals. The phenotypic effects of DNA methylation and local DNA sequence polymorphism are comparable, but they operate through largely distinct gene sets. Our epigenetic association analyses directly identify the relevant gene more frequently than genetic associations, likely due to reduced linkage disequilibrium. We identify numerous associations between methylation epialleles and diverse environmental conditions in native habitats, suggesting that intragenic methylation facilitates adaptation to fluctuating environments. Overall, our results demonstrate that epigenetic methylation variation fundamentally shapes phenotypic diversity in natural populations.

## Introduction

The neo-Darwinian or Modern synthesis is at the heart of evolutionary biology^1^. This framework posits that DNA sequence changes generate genetic variation, and mechanisms such as natural selection and genetic drift shape this variation to adapt populations to specific environments^2,3^. Epigenetic information, which can be encoded independently of the DNA sequence, is essential for cell fate determination, development, and environmental responses in eukaryotes^4–7^. In theory, if stably heritable and independent of DNA sequence polymorphism, epigenetic variation could influence inherited traits and contribute to adaptation^8–10^. Epiallelic variation in several angiosperm genes, including *Linaria vulgaris Cyc*, tomato *CNR* and *VTE3*, maize *Spm*, rice *D1*, oil palm *MANTLED*, and *Arabidopsis thaliana FWA, PAI2* and *IAA7*, influences phenotypic outcomes^11,12^. However, these epialleles are either too unstable for the epigenetic states to influence a response to selection^8–10^ (such as *Cyc*^13^, *D1*^14^ and *MANTLED*^15^), are confounded by *trans* genetic variation (such as *PAI2*^16^ and *IAA7*^12^), are artificially induced (such as *FWA*^17^ and *MANTLED*^15^), or evidence is lacking that heritable epiallelic variation at the gene occurs in nature (such as *CNR*^18^*, VTE3*^19^*, Spm*^20^ and *D1*^14^), in part because the relevant studies were conducted in domesticated crops. Due to these limitations, there is currently little evidence that epigenetic inheritance mediates phenotypic diversity or influences evolutionary outcomes within natural populations^11,21^.

DNA methylation can be epigenetically inherited over many generations^11,22^ and occurs in transposable elements (TEs) and bodies of transcribed genes^23–26^. Plant TEs are generally methylated in all sequence contexts – CG, CHG, and CHH (H being A, T, or C)^7,23,24,27^. TE methylation induces silencing^27^, confers genome stability^26,28^, can influence the expression of neighboring genes^12,29–33^, and its variation has been associated with all known epialleles^11,21^. Gene body methylation (gbM) occurs only in the CG context^23,24,34^, although genes can also feature TE-like methylation in all contexts (teM)^35,36^. TeM is associated with silencing^25,35,36^, but the function of gbM has been extensively debated. GbM is nearly ubiquitous in flowering plants^37,38^ and is common in animals^23,24,39^. In both groups, gbM preferentially resides in nucleosome-wrapped DNA within the exons of conserved, constitutively transcribed genes^25,40–43^. Conservation and phenomenological coherence suggest important functions^39^. Indeed, gbM is associated with (small) gene expression differences within and between plant species^36,44–48^, represses aberrant intragenic transcripts^49^, and appears to be under natural selection^46,47,50^. Moreover, loss of methyltransferase function causes developmental abnormalities in honeybees^51^, animals in which methylation is principally restricted to gene bodies^52^. However, loss of gbM has not been causatively linked to changes in gene expression in plants or animals^25,53,54^, leading to the proposals that gbM is a non-functional and somewhat deleterious by-product of TE methylation (in plants)^25,54,55^ or has functions unrelated to gene expression (in animals)^53^. Thus, the functional and evolutionary significance of gbM has been mysterious and controversial.

Recent studies have reported extensive natural variation in TE methylation, gbM and teM in *Arabidopsis*^35,36^. Methylation levels of natural accessions are associated with climate^35^, suggesting that methylation variation could be of adaptive significance. Furthermore, genetically induced methylation polymorphism can account for the inheritance of complex *Arabidopsis* traits^56–58^, and methylation changes have been linked to adaptation under artificial selection^59,60^. Variation in TE methylation and teM has been repeatedly linked to genetic variation^21^, but natural gbM variation is primarily epigenetic, at least in *Arabidopsis*^36,61^, and hence gbM appears particularly suited to mediate natural phenotypic variation. Nonetheless, whether variation of gbM or any other type of methylation underlies phenotypic diversity or drives the evolution of complex traits in natural populations is unknown^11^.

## Results

### GbM and teM are independent phenomena

To elucidate the biological significance of intragenic DNA methylation, we categorized genes of 948 *Arabidopsis* natural accessions into three epigenetic states: unmethylated (UM), gbM and teM using published data^35^. Consistent with published results^46^, we find that gbM conservation is bimodal, with genes most commonly methylated either in over 90% of accessions, or in ≤10% (Figure 1A). The high frequency of gbM conservation supports the hypothesis that gbM is maintained by natural selection^39,46,50^. In contrast, the vast majority of genes exhibit teM in ≤10% of accessions (Figure 1B), suggesting that teM is disfavored in most genes, likely due to its deleterious effects on expression^35^. A small fraction (7.3%) of teM genes is methylated in >50% of accessions (Figure 1B), suggesting that teM effects, potentially including transcriptional silencing^32,62^, might be advantageous in some genes and/or environments.

**Figure 1.**
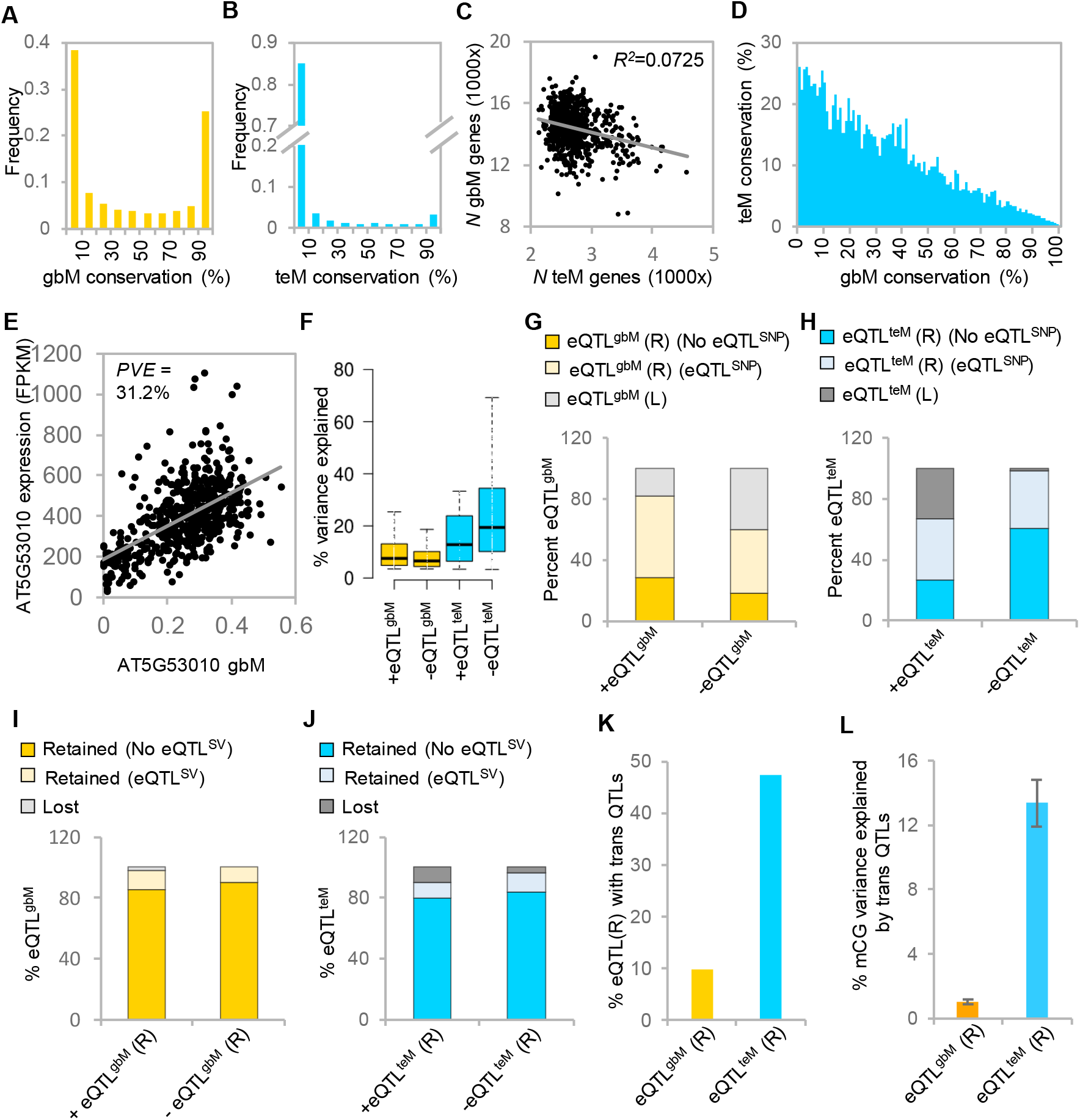
GbM is associated with gene expression independently of genetic variation in natural populations. (**A** and **B**) Frequency distribution of gbM (**A**) and teM (**B**) conservation across 33,056 *Arabidopsis* genes in 948 accessions. (**C**) Correlation between the number (*N*) of gbM and teM genes across accessions. (**D**) Average teM conservation within gbM conservation percentiles. (**E**) GbM and expression of AT5G53010 across accessions. Percent expression variance explained (*PVE*) by gbM is indicated. (**F**) Effect sizes of gbM and teM on expression of positive and negative eQTL^gbM/teM^ genes. (**G** and **H**) Frequency of retained or lost eQTL^gbM^ (**G**) and eQTL^teM^ (**H**) genes after accounting for expression GWA SNPs. (**I** and **J**) Frequency of retained or lost eQTL^gbM^ (**I**) and eQTL^teM^ (**J**) genes after accounting for structural variation (SV) effects on expression. (**K**) Percentage of eQTL^gbM/teM^ retained (R) genes that have *trans* genetic QTLs associated with the variation of gbM or teM. (**L**) Average effects (± standard error) of *trans* polymorphism on mCG variation in all retained eQTL^gbM/teM^ genes.

Numbers of teM and gbM genes are very weakly (negatively) correlated across accessions (Figure 1C) and are similarly weakly (positively) correlated under more restrictive published definitions^55^ of gbM and teM (Figure S1A). Additionally, intragenic teM prevalence across accessions tends to increase with decreasing gbM conservation under either gbM/teM definition (Figures 1D, and S1B- S1F), indicating that gbM and teM tend to occupy different genes. These results do not support the hypotheses that gbM originates as a by-product of teM^54^ or that gbM promotes the transition to teM^55^. Instead, intragenic gbM and teM appear largely independent, which is consistent with many linages having only TE methylation (fungi and some land plants) or only gbM (many invertebrates)^23–25^.

### Intragenic DNA methylation is associated with transcription in natural populations

To investigate the effects of *cis* gbM and teM variation on gene expression, we analyzed associations between CG methylation (mCG) and mRNA levels of individual genes among 625 accessions for which methylation and expression data are available^35^ using a linear model. We identified 614 +eQTL^gbM^ genes (eQTL stands for expression quantitative trait locus) that show a positive association between gbM and gene expression and 151 -eQTL^gbM^ genes that exhibit a negative association (Figures 1E and S2, and Tables S1 and S2) at a conservative significance threshold (Bonferroni α=0.05). Using less stringent thresholds, we identified 1,543 +eQTL^gbM^ and 715 -eQTL^gbM^ genes at 0.05 false discovery rate (FDR), and 1,887 +eQTL^gbM^ and 966 - eQTL^gbM^ genes at 0.1 FDR (Tables S1 and S2). Notably, +eQTL^gbM^ genes are 4-fold enriched under the stringent Bonferroni threshold, and only 2-fold enriched under FDR thresholds, suggesting that positive associations between gbM and expression are more likely to be real. Furthermore, +eQTL^gbM^ genes are more likely to have had gbM prior to the speciation of *A. thaliana* (Figure S3)^46^, suggesting that they are under selection to retain gbM.

Average effects of gbM on expression variance are around 10%, and gbM generally accounts for ≤10% of expression variance (85.3% of eQTL^gbM^ Bonferroni genes), but in exceptional cases gbM explains around 70% of expression variance (Figures 1F and S4A). Non-associated gbM genes (NA^gbM^) and eQTL^gbM^ genes have similar CG dinucleotide composition and length (Figures S5A and S5B) – key distinguishing characteristics of gbM genes^55^ – indicating that eQTL^gbM^ genes are fairly typical. However, +eQTL^gbM^ genes show lower expression than NA^gbM^ genes (Figure S5C), suggesting that gbM may have a stronger effect when transcription is weaker. Overall, our results suggest that gbM tends to have a modest positive effect on gene expression, which is consistent with published correlations between gbM and expression^36,44,45^.

In contrast to gbM, teM associations with expression are overwhelmingly negative. At Bonferroni α=0.05, we identified 202 genes in which teM is negatively associated with gene expression (-eQTL^teM^) and only 15 positively associated genes (+eQTL^teM^) (Table S1). As with gbM, the less stringent FDR thresholds produced more associations, but with a reduced ratio of -eQTL^teM^ to +eQTL^teM^ (Tables S1 and S2). TeM effects on gene expression are stronger than for gbM, as teM explains >20% of expression variance in 46% of eQTL^teM^ Bonferroni genes (Figures 1F and S4B). The primarily negative associations between teM and expression are consistent with existing knowledge^35,36^ and indicate that teM and gbM have fundamentally different effects on gene expression.

### GbM is associated with transcription in the absence of local SNP polymorphism

Because genetic and epigenetic variation can be linked in populations, we investigated whether epigenetic variants influence expression independently of *cis*-acting DNA sequence changes. To identify *cis* single nucleotide polymorphisms (SNPs) associated with expression of the 982 eQTL^gbM^/^teM^ Bonferroni genes we performed genome-wide association (GWA) analyses. For genes where *cis* SNP eQTLs (eQTL^SNP^) colocalized with eQTL^gbM^ or eQTL^teM^, we defined nested populations fixed for GWA SNPs and analyzed associations between mCG and expression within each population (Figure S6). Using this approach, we found that 82.4% of +eQTL^gbM^ were retained, either because an eQTL^SNP^ is not detected (28.7%), or because a +eQTL^gbM^ was detected in at least one nested population (53.7%) (Figure 1G and Table S3). Similarly, nearly all (98.5%) -eQTL^teM^ were retained (Figure 1H and Table S3). However, 39.7% of -eQTL^gbM^ and 33.3% of +eQTL^teM^ were lost after accounting for SNP variation (Figures 1G and 1H, and Table S3). These results support the conclusion that most +eQTL^gbM^ and -eQTL^teM^ associations are real, whereas many -eQTL^gbM^ and +eQTL^teM^ associations are probably spurious and detected due to linkage between epigenetic and local SNP variation.

Allelic heterogeneity is common in *Arabidopsis*^63–67^. To account for confounding of epigenetic eQTLs by residual *cis* SNPs (other than GWA SNPs), we defined haplogroups so that accessions in a haplogroup are invariant for SNPs within retained epigenetic eQTL genes and 4 kb upstream and downstream. This analysis created numerous small groups (Table S4), and consequently many associations are expected to be lost due to lack of statistical power (Figure S7A). Nonetheless, we detected significant associations between mCG and gene expression within at least one haplogroup for a clear majority of gbM eQTLs (70.8% +eQTL^gbM^, 64.4% - eQTL^gbM^), for a similar fraction of -eQTL^teM^ (72.2%) and for a smaller but substantial proportion of +eQTL^teM^ (37.5%; Figures S7B and S7C, and Table S5). Therefore, our results show that intragenic methylation is quantitatively associated with gene expression independently of local SNPs.

### Local structural genetic variation does not confound eQTL^gbM^

To evaluate whether eQTL^gbM/teM^ are confounded by structural genetic variation (SV) such as insertions and deletions, we analyzed associations between methylation and expression in populations invariant for published transposable element insertion polymorphisms (TIPs)^62^ within retained epigenetic eQTL genes and 4 kb upstream and downstream. We found that 98.2% of +eQTL^gbM^ are not confounded by these SVs, either because an eQTL^SV^ is not detected (85.4%) within the analyzed region, or because a +eQTL^gbM^ was detected in the population without SV (12.8%; Figure 1I). Similarly, all -eQTL^gbM^ are retained after accounting for SV (Figure 1I). In comparison to eQTL^gbM^, a higher fraction of eQTL^teM^ (10% +eQTL^teM^ and 4% -eQTL^teM^) are lost after accounting for SV (Figure IJ). More frequent confounding of eQTL^teM^ by SV is expected, because teM variation is known to be associated with TE SVs^32,33,36^, which may promote teM and repress expression (or *vice versa*).

Undetected cryptic SV exists in *Arabidopsis*^62^, but it is very unlikely to explain a substantial fraction of eQTL^gbM^. First, fewer than 2% of +eQTL^gbM^ are confounded by known TIPs, so cryptic TIPs would have to be unrealistically abundant (>10-fold of known TIPs) to explain most +eQTL^gbM^. Second, most SVs are linked to SNPs, and SNP-based GWA therefore identifies most SV-trait associations^68^. Systematic confounding by cryptic SV would thus manifest as systematic confounding by SNPs. Because >80% of +eQTL^gbM^ are not confounded by SNPs, to explain most +eQTL^gbM^, cryptic SV would have to be fundamentally different from known SV by being linked with gbM variation, but unlinked with genetic variation (SNPs). This is neither parsimonious nor plausible. In contrast, the conclusion that eQTL^gbM^ are very rarely confounded by SV is consistent with the findings that gbM variation develops independently of SV in mutation accumulation (MA) lines^69,70^, gbM variation in MA lines is highly correlated with that in natural accessions^71^, gbM is generally neither associated with nor affected by SV in accessions^33^, and local gbM variation in the *Arabidopsis* population is primarily caused by stochastic epigenetic fluctuations^61^.

### GbM patterns are weakly influenced by *trans* genetic polymorphism

Our results show that natural DNA methylation variation influences gene expression largely independently of local genetic polymorphism. However, methylation patterns could be governed by *trans* genetic polymorphism, such as mutations in DNA methyltransferases and other regulatory genes. Such polymorphism has indeed been identified for teM^35,36,44,72,73^, and consistently we find 47.4% of retained eQTL^teM^ genes are significantly affected by *trans* QTLs (Figure 1K). Each *trans* QTL on average explains 28.5% of associated teM variation, with all *trans* QTLs accounting for 13.4% of teM variation in tested genes (Figures 1L and S8A). Thus, a substantial amount of teM variation at eQTL^teM^ genes is not epigenetic but governed by *trans* (presumably genetic) polymorphism.

Previous studies of *trans* effects on gbM have produced mixed results. Several analyses of broadly distributed *Arabidopsis* accessions did not identify strong *trans* influences on gbM^35,36^, whereas a study of Swedish *Arabidopsis* did find strong *trans* effects for around 1,300 gbM genes^44^. Indeed, if we consider only the 133 Swedish accessions, we find strong *trans* effects, with on average 37.9% of gbM variance explained at 11.5% of eQTL^gbM^ (4.4% gbM variance explained overall; Figures S8B-S8D). However, when we consider all 625 worldwide accessions, these *trans* effects nearly disappear – 9.7% of genes have significant *trans* QTLs, which on average explain 10.5% of gbM variance, with *trans* genetic variation accounting for only 1% of gbM variance over all tested eQTL^gbM^ (Figures 1K, 1L, and S8A-S8D). Notably, a panel of 133 randomly chosen worldwide accessions (same size as the Swedish panel) produced results that are almost identical to those of the Swedish panel, and significantly different from the entire worldwide panel (Figures S8B-S8D). This indicates that estimates of *trans* effects on gbM variation are inflated in analyses of small populations, a phenomenon known as the Beavis effect^74,75^. Therefore, *trans* genetic polymorphism explains only a very minor fraction of overall gbM variation.

Recently, we identified several *trans* factors that influence global gbM^61^. However, in the same study we found that local gbM variation is primarily caused by epigenetic fluctuations, with *trans* factors affecting overall global gbM levels of accessions^61^. Consistently, global gbM levels of accessions are weakly associated with gbM levels of individual genes (*R^2^*<0.1 for ∼80% genes, linear model; Figure S8E), indicating that gbM at individual genes evolves independently of *trans* factors that cause global gbM differences, such as the high overall gbM in *Arabidopsis* accessions from northern Sweden^35,61^. Taken together, our results support earlier conclusions that local gbM variation is not strongly affected by *trans* (or *cis*) genetic polymorphism, but is instead primarily epigenetic^36,61^.

### GbM and *cis* SNPs are associated with comparable effects on expression variance

Our results indicate that gbM is associated with expression variation independently of local and *trans* genetic polymorphism in natural populations. A key question is how much expression variance is accounted by gbM relative to local genetic variation. A published attempt to employ a statistical model to partition variance components was confounded by the linkage of methylation and SNPs, so that the model could find that most of the expression variance is accounted by SNPs, but also that SNP effects are marginal after accounting for associations between methylation and expression^45^. An alternative approach is to calculate the effects of gbM on expression variation after accounting for genetic variation and *vice versa*. Because the confounding effects of SV and *trans* genetic polymorphism on eQTL^gbM^ are effectively negligible (Figures 1I-1L), we focused on *cis* SNPs. We found a strong decrease in the percent expression variance explained by gbM for lost positive and negative eQTL^gbM^ after accounting for SNPs (Figure 2A). However, we did not find a significant change for gbM effects on expression of retained positive and negative eQTL^gbM^ genes after accounting for SNPs (Figure 2A), with gbM on average explaining about 10% of expression variance. We obtained analogous results for effects of teM on expression variance, though these are not statistically significant (Figure S9A).

**Figure 2.**
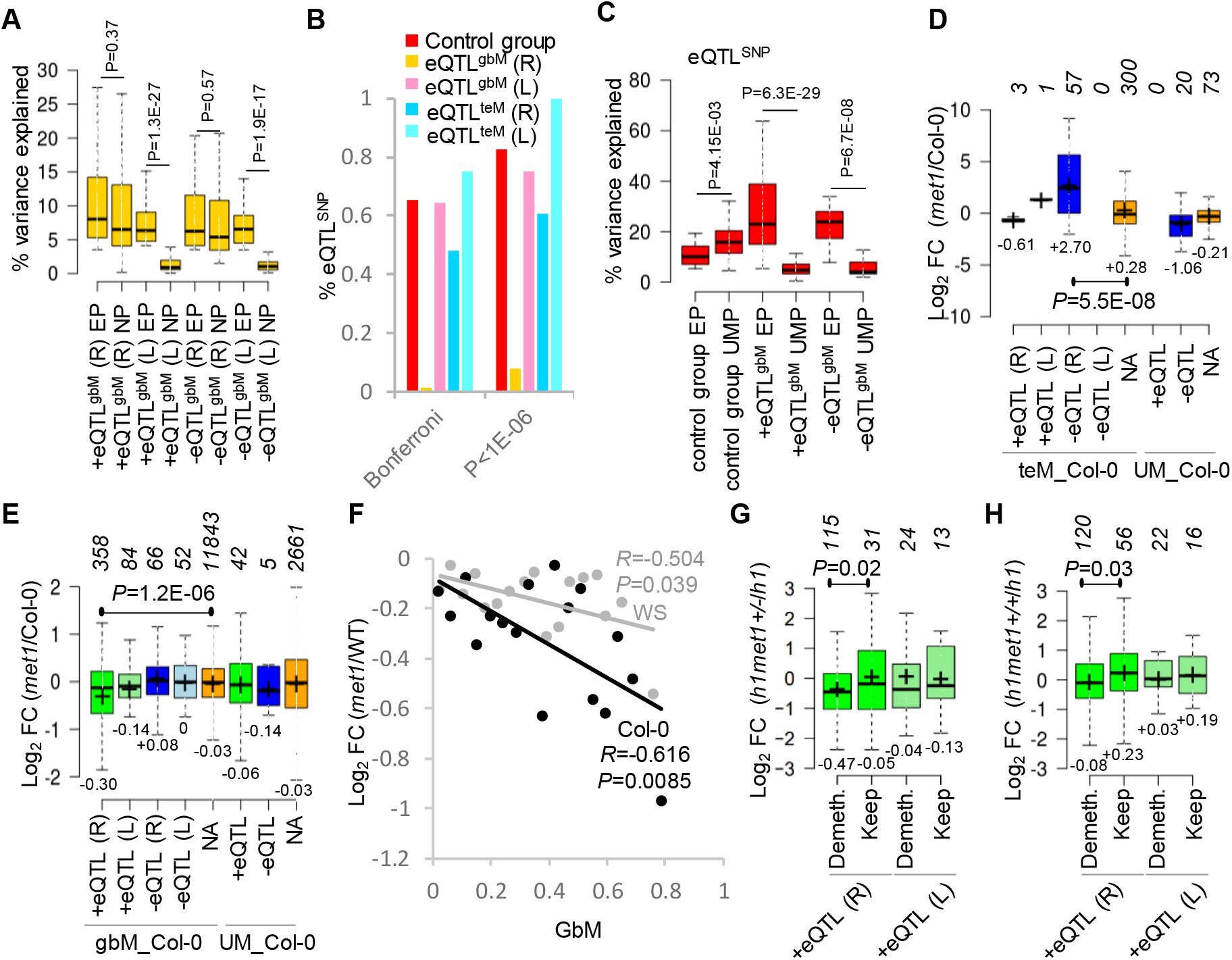
GbM promotes gene expression. (**A**) Effect sizes of gbM on expression of positive and negative retained (R) or lost (L) eQTL^gbM^ genes in the entire population (EP) and after accounting for expression GWA SNPs in nested populations (NP). (**B**) Percent of eQTL^SNP^ detected as significant at the Bonferroni threshold and an arbitrary relaxed threshold (*P*<1E-06) after accounting for methylation variation. (**C**) Effects of SNPs on expression variation of a subset of control group (NA^gbm^) genes and retained eQTL^gbM^ genes in the entire (EP) and unmethylated populations (UMP). (**D** and **E**) Expression in *met1* seedlings compared to Col-0 of Bonferroni (α=0.05) eQTL^teM^ (**D**) and eQTL^gbM^ (**E**) genes retained (R) or lost (L) after accounting for genetic variation. Genes methylated (gbM_Col-0; teM_Col- 0) and unmethylated (UM_Col-0) in Col-0 were analyzed separately. Numbers of genes within each group are indicated. *P*, Wilcoxon rank sum test. (**F**) Relationship between gbM levels of retained +eQTL^gbM^ genes in Col-0 and WS wild type (WT) plants and log2 fold change in expression in *met1* compared to wild type. Genes were grouped by gbM levels. *R* and *P* values correspond to Pearson’s correlation. (**G** and **H**) Expression in *h1met1+/-* (**G**) and *h1met1+/+* (**H**) compared to *h1* of Bonferroni (α=0.05) retained (R) or lost (L) eQTL^gbM^ genes that are either demethylated or keep methylation. *P*, Student’s t-test.

To evaluate how methylation in gene bodies confounds *cis* eQTL^SNP^, we performed GWA analysis only in UM accessions for the 185 retained eQTL^gbM^ and 35 retained eQTL^teM^ genes that are UM in at least 100 accessions (185 NA^gbM^ genes that are UM in at least 100 accessions were randomly selected for a control group). For the control group, 65.3% of *cis* eQTL^SNP^ detected in the entire population at the Bonferroni threshold remain significant after accounting for methylation effects, and 82.7% are significant at a relaxed *P*<1E-06 threshold (Figure 2B). Most of the eQTL^SNP^ detected in the entire population are also detected in UM populations for lost eQTL^gbM^, and retained as well as lost eQTL^teM^ genes (Figure 2B). However, very few (1.3% at the Bonferroni threshold and 7.8% at *P*<1E-06) eQTL^SNP^ remain in retained eQTL^gbM^ genes after accounting for methylation (Figure 2B). This indicates that most *cis* eQTL^SNP^ are detected in retained eQTL^gbM^ genes due to linkage with gbM, as the eQTL^SNP^ effectively disappear in analyses of unmethylated accessions.

Local SNPs on average explain 12.5% of expression variance in control group genes having *cis* eQTL^SNP^ at the Bonferroni threshold in the entire population (Figure 2C), which is comparable to the expression variance explained by gbM at retained eQTL^gbM^ (10.8% for +eQTL^gbM^; 8.7% for -eQTL^gbM^; Figure 2A). Analysis of control genes in UM accessions produces a significant increase in expression variance explained by *cis* SNPs (Figure 2C), which is expected due to the smaller population sizes^74,75^. An analogous (non-significant) increase in expression variance explained by *cis* SNPs is observed in UM populations for retained eQTL^teM^ (Figure S9B). In contrast, we find a 4 to 5-fold decrease in the effects of *cis* SNPs on expression variance of retained positive and negative eQTL^gbM^ genes in UM populations (Figure 2C), indicating that these eQTLs^SNP^ are overwhelmingly detected due to linkage of SNPs with gbM. Importantly, this analysis does not evaluate the influence of *trans* genetic polymorphism on gene expression, but seeks to compare the effects of local gbM and SNP polymorphism. Our results show that, in comparison to *cis* SNPs, gbM principally accounts for expression variance of retained eQTL^gbM^ genes and explains about the same extent of expression variance as *cis* SNPs do in control genes.

### Loss of gbM specifically downregulates +eQTL^gbM^ genes

Our results show that gbM explains a significant amount of natural gene expression variation. To determine whether intragenic DNA methylation directly affects gene expression, we analyzed published RNA sequencing data from *met1* mutant seedlings and wild type Col-0 controls^49^. Mutation of the *MET1* methyltransferase causes complete loss of gbM and nearly complete loss of mCG throughout the genome^76^. For genes passing the Bonferroni threshold, we established three groups of control loci that should not be directly affected by *met1*: NA^gbM/teM^ genes (no association between methylation and expression), eQTL^gbM^/^teM^ genes that lack methylation in the wild type control accession (Col-0), and Col-0 methylated eQTL^gbM^/^teM^ genes lost due to linked GWA SNP variation. None of the control groups showed significantly altered expression in *met1* plants, nor did any group of -eQTL^gbM^ or +eQTL^teM^ genes (Figures 2D and 2E). Only -eQTL^teM^ and +eQTL^gbM^ genes methylated in Col-0 are significantly affected by *met1* (Figures 2D and 2E).

Col-0 methylated -eQTL^teM^ genes are strongly overexpressed in *met1* (over 6-fold of wild-type; Figure 2D), consistent with the established repressive activity of teM^25,35,36^. In contrast, +eQTL^gbM^ genes are modestly downregulated (expressed at about 80% of wild-type; Figure 2E). Analysis of genes that passed less stringent significance thresholds (0.05 and 0.1 FDR) produced similar results (Figures S10A and S10B). Analysis of additional Col-0 *met1* leaf ^49^ and inflorescence^77^ RNA-seq datasets, and of *met1* microarray data in the WS wild-type background^78^, also produced analogous results (Figures S11A-S11F). *MET1* inactivation alters non-CG methylation and histone modifications^79^, which could influence gene expression, but are not significantly changed in any relevant gbM gene category (Figures S12A and S12B). The reduced expression of only those +eQTL^gbM^ genes that are methylated in Col-0 or WS and not confounded by genetic variation indicates that gbM promotes gene expression. Furthermore, retained +eQTL^gbM^ genes with higher methylation in parental accessions show stronger downregulation in *met1* (Col-0 and WS; Figure 2F), consistent with gbM quantitatively promoting expression.

To further evaluate whether gbM loss alters gene expression, we analyzed a plant that is heterozygous for *met1* (*met1*+/-), has relatively normal TE methylation, and limited gbM loss^49^. This plant also contains loss-of-function mutations in two histone H1 genes^49^ and therefore expression was analyzed with respect to *h1* mutant controls. Retained +eQTL^gbM^ genes demethylated in this plant have significantly decreased (∼35%) expression in comparison to retained +eQTL^gbM^ genes that maintain gbM (Figure 2G), specifically linking gbM loss with reduced expression. Moreover, we isolated six *h1met1*+/+ progeny of *h1met1*+/-. These plants exhibit mosaic demethylation of gbM genes, whereas TE methylation is comparatively normal (Figures S13A and S13B). Retained +eQTL^gbM^ genes demethylated in *h1met1*+/+ plants display significantly reduced (∼25%) expression compared to retained +eQTL^gbM^ genes that keep gbM (Figure 2H). Altogether, we find that loss of gbM at retained +eQTL^gbM^ genes consistently produces decreased expression, regardless of the genetic background (Col-0 or WS), tissues (seedlings, leaves, or inflorescence), presence of functional *MET1*, or the extent of global teM or gbM perturbation. Our results therefore constitute strong evidence that gbM directly promotes gene expression. The sensitivity of +eQTL^gbM^ gene expression to gbM also reinforces the conclusion that detection of these eQTLs is not substantially confounded by linked cryptic genetic polymorphism in natural accessions, because the effects of such polymorphism should be independent of DNA methylation.

### Intragenic DNA methylation is associated with plant fitness in a hot and dry climate

Modulation of gene expression by natural methylation polymorphism potentiates the possibility that methylation epialleles drive phenotypic variation in natural populations. To uncover how DNA methylation shapes natural phenotypic diversity, we performed epigenome-wide association (epiGWA) analyses for relative fitness using genic mCG levels in a linear model. We used published relative fitness data of *Arabidopsis* accessions grown in open-sided greenhouses in Madrid and Tübingen^80^. Linear model associations can be confounded by population structure, producing elevated rates of false-positives^81^. Therefore, we used a ‘genomic control factor’^82^ to correct association statistics (Table S6). We identified two significant (FDR 0.05) epigenetic relative fitness QTLs (rfQTLs) for which mCG is positively correlated with fitness in Madrid (hot climate) under low rainfall and high-density population growth (MLP conditions; Figures 3A and S14). One (rfQTL^gbM^) corresponds to gbM in *Proline Transporter 1* (*PROT1*; AT2G39890) and the second (rfQTL^teM^) to teM in an uncharacterized gene encoding a FBD/Leucine Rich Repeat containing protein (AT1G19410). Identification of *PROT1* was particularly intriguing, as regulation of proline accumulation is a hallmark plant response to abiotic stresses including drought^83^.

**Figure 3.**
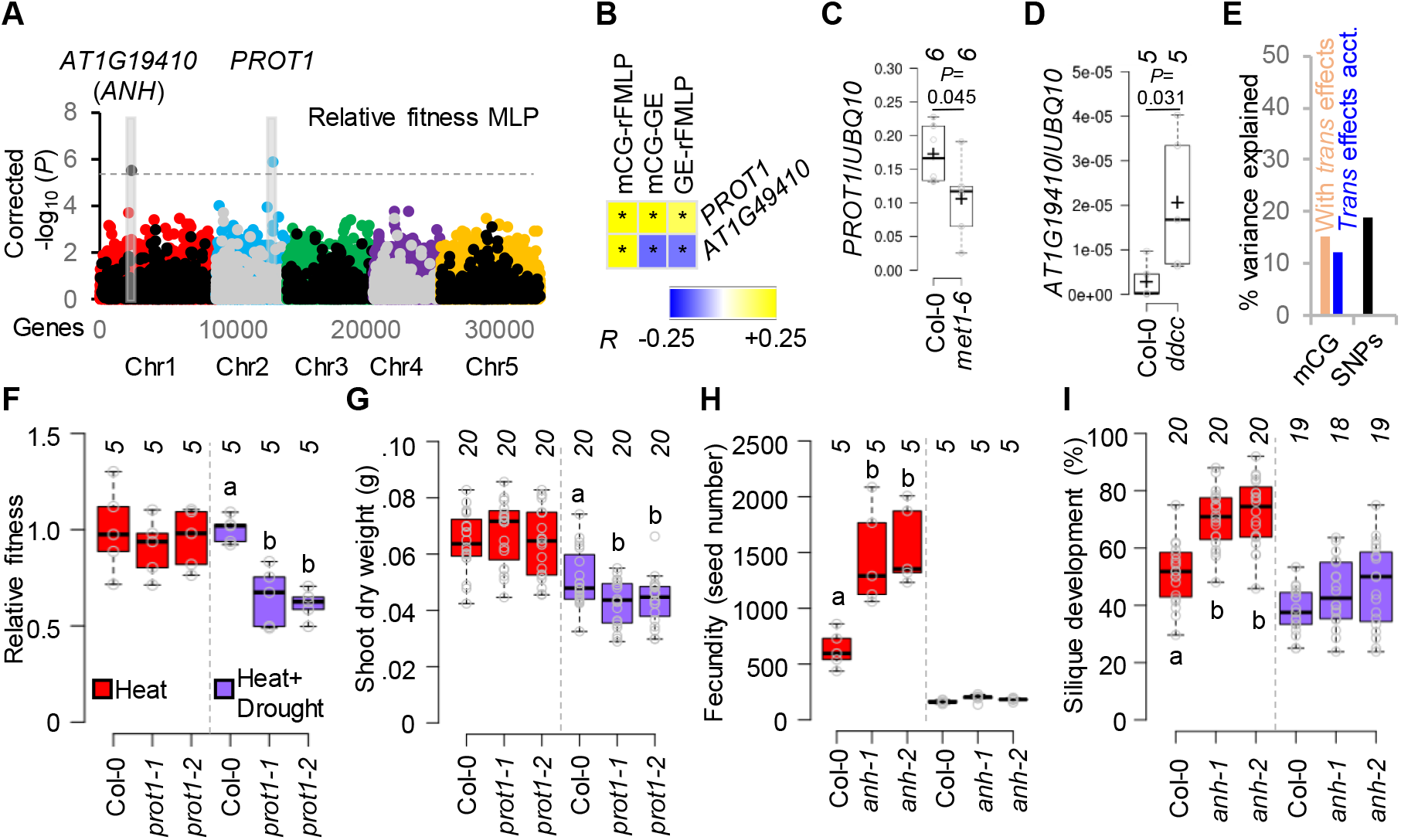
Intragenic DNA methylation affects relative fitness. (**A**) Manhattan plot of epiGWA mapping for relative fitness in MLP conditions. GbM markers are depicted with colored dots and teM markers are shown using grey and black dots. Horizontal dashed line shows 0.05 FDR. (**B**) Tripartite associations between intragenic DNA methylation (mCG), gene expression (GE), and relative fitness in MLP (rFMLP). \**P*<0.05, Pearson’s correlation test. (**C** and **D**) Transcript levels of *PROT1* (**C**) or *AT1G19410* (*ANH*; **D**) relative to *UBQ10* in Col-0 and indicated mutants, assessed by qRT-PCR in six (**C**) or five (**D**) biological replicates. *P*, two-tailed Student’s t test. (**E**) Percent variation in MLP relative fitness explained by epigenetic (mCG) and genetic (SNPs) QTLs. Effects of *trans* genetic QTLs on mCG variation were accounted (acct.). (**F**-**I**) Fitness (**F**) or shoot dry weight (**G**) of *prot1* mutants (two independent alleles) and fecundity (**H**) or fertility (**I**) of *anh* mutants (two independent alleles) relative to Col-0 under heat or joint heat and drought stress. Numbers of independent experiments are indicated for fitness (**F**) and fecundity (**H**), and plant numbers are indicated for shoot weight (**G**) and fertility (**I**). Different letters signify *P*<0.05, one-way ANOVA, Tukey’s test.

*PROT1* is a typical gbM gene methylated in 95.1% of accessions, but *AT1G19410* is unusual, with teM in 73.4% of accessions (Table S7), suggesting that teM in this gene may be functionally important. We identified *PROT1* as a +eQTL^gbM^ at the Bonferroni threshold, and *AT1G19410* as a -eQTL^teM^ at FDR 0.05 (Table S2), consistent with the general effects of gbM (positive) and teM (negative) on gene expression. Tripartite analysis confirmed that *PROT1* and *AT1G19410* mCG levels are significantly associated with expression across accessions (positively for *PROT1* and negatively for *AT1G19410*), and revealed that expression of both genes is significantly associated with relative MLP fitness (again positively for *PROT1* and negatively for *AT1G19410*; Figure 3B). Consistent associations between mCG, fitness, and expression were detected after accounting for SV, in two independent *PROT1* and *AT1G19410* SNP haplogroups, and in haplogroups after accounting for SV (Figure S15A and Table S8). *PROT1* is downregulated by 38% in *met1* (Figure 3C), confirming that gbM promotes *PROT1* expression. *AT1G19410* teM is lost in plants that lack DRM and CMT methyltransferases (Figure S15B), and in such *ddcc* mutants^84^ *AT1G19410* expression increases about 7-fold (Figure 3D), confirming the repressive effect of teM.

To understand the relative contribution of epigenetic and genetic variants to trait heritability we performed genetic GWA analyses for relative fitness. We detected two significant genetic QTLs (rfQTLs^SNP^) only in MLP (Table S9), suggesting that these harsh conditions (heat, drought, crowding) are conducive to identification of fitness QTLs. Notably, rfQTLs^gbM/teM^ do not colocalize with rfQTLs^SNP^ (Figure S16). Heritability estimates indicate that the two rfQTLs^SNP^ explain 18.9% of the relative fitness variance in MLP environment (Figure 3E). Strikingly, the two epigenetic rfQTLs also account for a substantial proportion of fitness variance (12.1% after accounting for *trans* QTL effects; Figure 3E). These results illustrate that natural epigenetic variation substantially contributes to the heritability of a complex plant trait.

### *PROT1* and AT1G19410 (*ANAHITA*) regulate plant fitness

The above results make clear predictions about the effects of gene inactivation on fitness in hot and dry conditions: *PROT1* inactivation should reduce fitness, whereas *AT1G19410* inactivation should enhance fitness. Genetic inactivation of *PROT1* indeed caused about 35% fitness reduction under experimentally induced joint heat and drought stress (Figure 3F). Mutant plants produced less biomass and had decreased survival to fruit, but had the same fecundity (seed set) as wild type (Figures 3G and S17A). Consistently, *PROT1* gbM is specifically associated with survival in MLP conditions (Figures S18A-S18C). Thus, the effects of *PROT1* inactivation match those expected from the associations between gbM, gene expression, and relative MLP fitness (Figure 3B).

Inactivation of *AT1G19410* resulted in a slight (∼13%) but non-significant increase in relative fitness under heat and drought stress (Figure S17B). However, *AT1G19410* mutants have greatly enhanced (>2-fold) fitness under heat stress alone, with >2-fold increased fecundity and significantly increased fertility (percent of flowers developing siliques; Figures 3H, 3I, and S17B). Therefore, we named *AT1G19410 ANAHITA* (*ANH*) after the ancient Persian goddess of fertility and water. Notably, the association of *ANH* teM is much stronger with fecundity than survival in MLP conditions (Figures S18D-S18F). Thus, although both genes influence relative fitness under experimental conditions, *PROT1* specifically influences survival, whereas *ANH* affects fecundity. These results reveal the power of epiGWA analyses to identify novel regulators of plant fitness.

### Flowering trait heritability is comparably influenced by epigenetic and genetic variation

Our results demonstrate that gbM and teM variation influences plant fitness, and thus presumably adaptation to the environment. A trait generally recognized as adaptive in *Arabidopsis* populations is timing of the transition to flowering to synchronize reproduction with the environment^85–87^. Therefore, we performed epiGWA and GWA analyses using published data for nine flowering-related traits: flowering time at 10 °C (FT_10°C) and 16 °C (FT_16°C), number of days for inflorescence stalk to reach 1 cm (DTF2), number of days to opening of the first flower (DTF3), number of cauline leaves (CLN), number of rosette leaves (RLN), cauline branch number, number of primary inflorescence branches, and length of the primary inflorescence stalk^88^. Due to the structure of flowering phenotypes in *Arabidopsis* populations^89^, we could not use a linear model with genomic control to analyze quantitative methylation variation (Figure S19 and Table S10). Instead, we used a mixed linear model (MLM) that effectively corrects population structure^90–93^ (Table S11) to analyze associations between phenotypes and binary epiallelic states of genes (UM or gbM; UM or teM).

Six of the traits we examined (FT_10°C, FT_16°C, DTF2, DTF3, CLN, and RLN) are correlated with some shared genetic control^94^. EpiGWA mapping detected eight epigenetic flowering QTLs (fQTLs) at FDR 0.05 (all gbM) for these phenotypes, and no epigenetic QTLs associated with the other flowering traits (Figures 4A and S20). Seven fQTLs^gbM^ contain SVs, but all loci except *AT4G33560* were retained after accounting for SV (Figures S21A-S21G). Two fQTLs^gbM^ localized to the well-known flowering regulators *Flowering locus* C (*FLC*) and *FRIGIDA* (*FRI*)^95–100^ (Figure 4A). To examine the roles of genes underlying the remaining six associations (*AT1G26795*, *AT1G51820*, *AT2G18210*, *AT3G09530*, *AT3G43860*, and *AT4G33560*) in transition to flowering, we examined FT_16°C in mutant lines. Mutants in all genes except *AT3G43860* have significantly altered FT_16°C (Figure 4B). Given the extensive analyses of *Arabidopsis* flowering time determinants performed over several decades^101^, the discovery of five new flowering time loci underscores the power of epiGWA to identify new functional genes.

**Figure 4.**
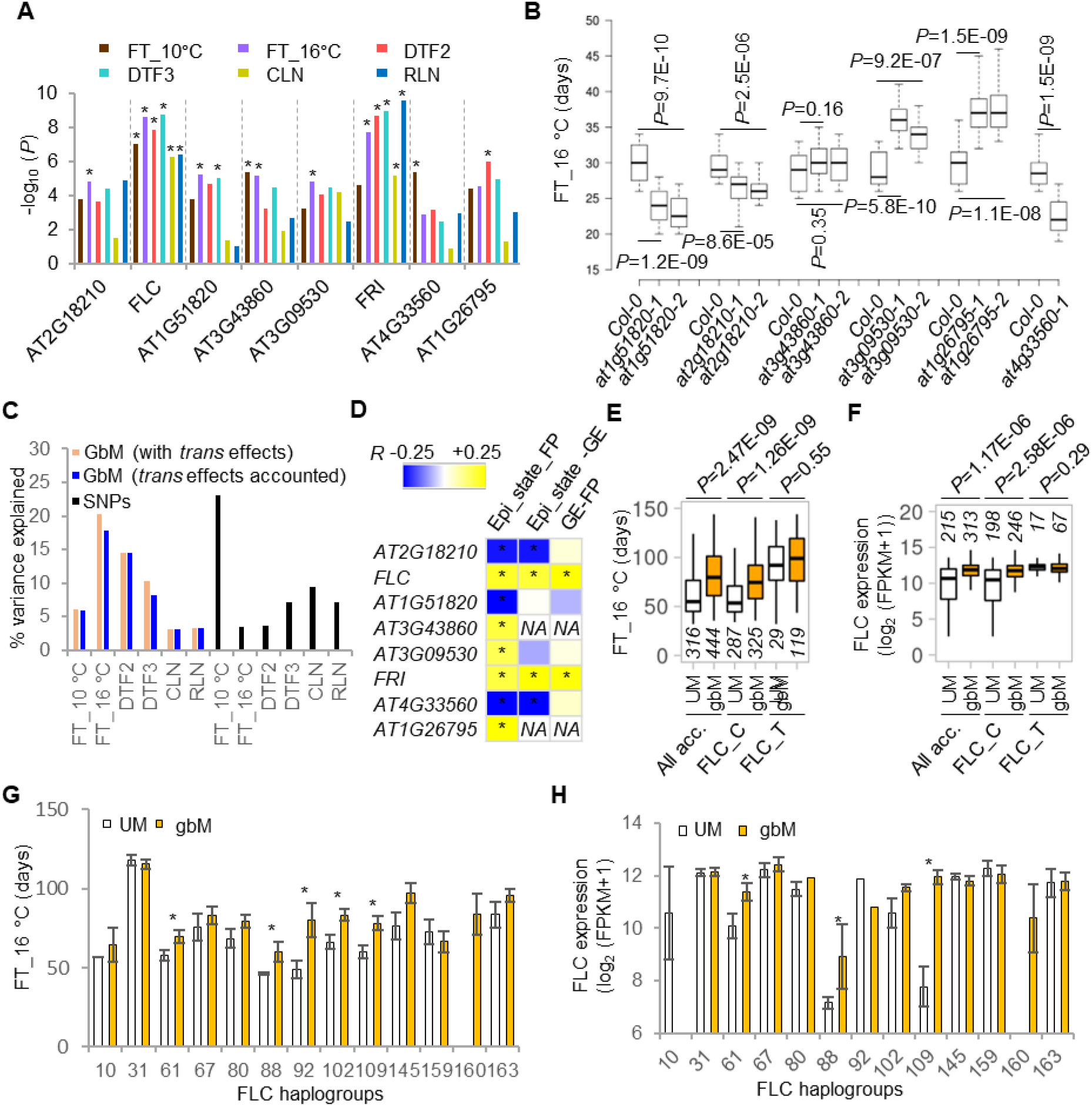
Flowering time is comparably influenced by natural genetic and epigenetic variation. (**A**) *P* values (-log10) of epigenetic flowering QTLs. *Significant at 0.05 FDR. (**B**) Genetic inactivation of fQTL^gbM^ genes causes altered flowering in Col-0. Flowering time of Col-0 and knock-out mutants of *AT1G51820*, *AT2G18210*, *AT3G43860*, *AT3G09530*, *AT1G26795*, and *AT4G33560*. Plants were grown at 16 °C. *P*, two-tailed t test. (**C**) Percent variation in flowering phenotypes explained by epigenetic (gbM) and genetic (SNPs) QTLs. Effects of *trans* genetic QTLs on gbM variation are accounted. Effect of *cis* GWA SNP and SV on fQTL^gbM^ are also accounted. (**D**) Tripartite associations between epiallelic states (Epi_state), gene expression (GE), and flowering phenotypes (FP). FT_10°C was used for AT4G33560, DTF2 for AT1G26795, and FT_16°C for the remaining fQTLs^gbM^. \**P*<0.05, MLM epiGWA for Epi_state-FP and Epi_state-GE and Pearson’s correlation test for GE-FP. (**E** and **F**) Association of FLC epiallelic states with FT_16°C (**E**) and FLC expression (**F**) in all accessions and nested populations (FLC_C and FLC_T) fixed for FLC SNP associated with flowering time. Numbers of accessions for each group are indicated. *P* values correspond to MLM epiGWA. (**G** and **H**) Association of FLC epiallelic states with FT_16°C (**G**) and FLC expression (**H**) in thirteen FLC haplogroups. Only accessions without TE polymorphism around FLC are considered to account for SV. \**P*<0.05, two-tailed Student’s t-test.

GWA analysis using SNPs identified eleven genetic flowering QTLs (fQTLs^SNP^) associated with the six correlated traits, including *FLC* and *FRI* (the only overlaps with fQTLs^gbM^), as well as three other known flowering regulators: *FT*, *SVP*, and *VIN3*^102–104^ (Figure S22 and Table S12). Except for FT_10°C, fQTLs^SNP^ explain <10% of trait heritability (Figure 4C). Strikingly, epigenetic variation explains >10% of heritability for three flowering traits before accounting for the effects of *trans* QTLs on gbM variation, and two traits after accounting for *trans* effects (Figure 4C). Accurate estimation of natural variation contribution to trait heritability is challenging^81,105^ and we analyzed only the specific genetic and epigenetic QTLs we detected. Hence, the values in Figure 4C should be interpreted with caution. For example, it is very unlikely that genetic variation explains 23% of heritability for FT_16°C but only 3.4% for FT_10°C (Figure 4C). Instead, the key conclusion is that epigenetic gbM variation and genetic variation explain comparable amounts of flowering trait heritability.

### *FLC* gbM explains *FLC* expression and flowering time in *Arabidopsis* populations

We chose two fQTLs^gbM^, *FLC* and *FRI*, for further analyses. Not only are these extensively studied regulators of flowering time and its natural variation^99,100^, but we found them associated with all (*FLC*) or nearly all (*FRI*) of the six correlated flowering phenotypes (Figure 4A). Furthermore, both genes show consistent significant associations between gbM, flowering and gene expression (Figure 4D), and both were detected as +eQTL^gbM^ at the Bonferroni threshold (Table S2). However, the *FRI* +eQTL^gbM^ was lost after accounting for genetic (SNP) variation (Table S3), *FRI* expression is not significantly altered in *met1* plants (Figure S23A), and we did not detect significant associations between *FRI* epiallelic states and FT_16°C or expression within nested genetic populations (Figures S23B and S23C). Furthermore, we found significant linkage disequilibrium^106^ (*r*=0.725, *D*^/^=0.824, *P*<0.0001) between methylation states and *FRI* SNPs, suggesting that *FRI* epigenetic and genetic QTLs are redundant. Therefore, *FRI* was excluded from our estimates of gbM effects on flowering trait heritability (Figure 4C).

In contrast to *FRI*, the *FLC* +eQTL^gbM^ was retained after accounting for genetic variation (Tables S3 and S5; expression in *met1* could not be tested because *FLC* lacks gbM in Col-0, Figure S24A), and we identified significant associations between *FLC* epiallelic states, FT_16°C and expression within a nested population as well as after accounting for SV (Figures 4E, 4F, S21G, and S21H). Because multiple *FLC* alleles affect flowering time or vernalization response^32,96,97^, we defined thirteen *FLC* haplotypes (Table S8), all of which contain gbM accessions (Figure 4G), suggesting complex gbM evolution at this locus. GbM accessions display significantly delayed flowering (FT_16°C) within five haplotypes (delay of >18 days in three haplotypes; Figure 4G), and significantly higher *FLC* expression in three of these haplotypes (Figure 4H). These findings demonstrate that gbM variation explains *FLC* expression and flowering time independently of *FLC* SNP and SV variation. Moreover, gbM at *FLC* is not associated with global gbM levels (Figure S24B) and is not influenced by *trans* QTLs in the worldwide population (Figure S24C), indicating that *FLC* gbM variation is epigenetic.

*Arabidopsis* flowering time variation is correlated with geographic regions and functionally distinct *FLC* alleles can display restricted geographic distributions^97^. We find that the density of accessions carrying *FLC* UM and gbM epialleles is generally comparable across latitudes, except 48-52° where density of UM accessions is higher, and around 63° (northern Sweden) where *FLC* gbM accessions are more common (Figures S24D-S24F). Taking advantage of the large number of accessions distributed within 48-52° latitude, we detected a strong association (*P*=7.81E-05) between epiallelic states of *FLC* and flowering time (Figure S24G). The *FLC* fQTL^gbM^ is thus not confounded by links between flowering time and latitude, variation of global gbM (Figure S24B), *FLC* genetic polymorphism (Figures 4E, 4F, 4G, 4H, S21G, and S21H), or *trans* genetic polymorphism (Figure S24C).

### GbM of the *LGM* gene explains the transition to flowering in simulated climates

*Arabidopsis* flowering time variation has been characterized under simulated local climates in Spain and Sweden across two seasonal plantings: spring and summer in 2008 and 2009^86^. EpiGWA identified significant fQTLs^gbM^ (FDR 0.05) associated with flowering time in simulated local climates within two uncharacterized genes, *AT4G15750* and *AT5G49440* (Figures 5A and S25, and Table S13). The *AT4G15750* fQTL^gbM^ shows sensitivity to seasonal planting in Sweden and to climate (Spain vs. Sweden) in summer (Figures S26A-S26D), whereas the *AT5G49440* fQTL^gbM^ is sensitive to seasons in Spain and to climate in spring (Figures 5B, 5C, S26E, S26F). Seasonal and climate sensitivities suggest these fQTLs^gbM^ influence flowering in response to environmental inputs, and hence could underlie local adaptation.

**Figure 5.**
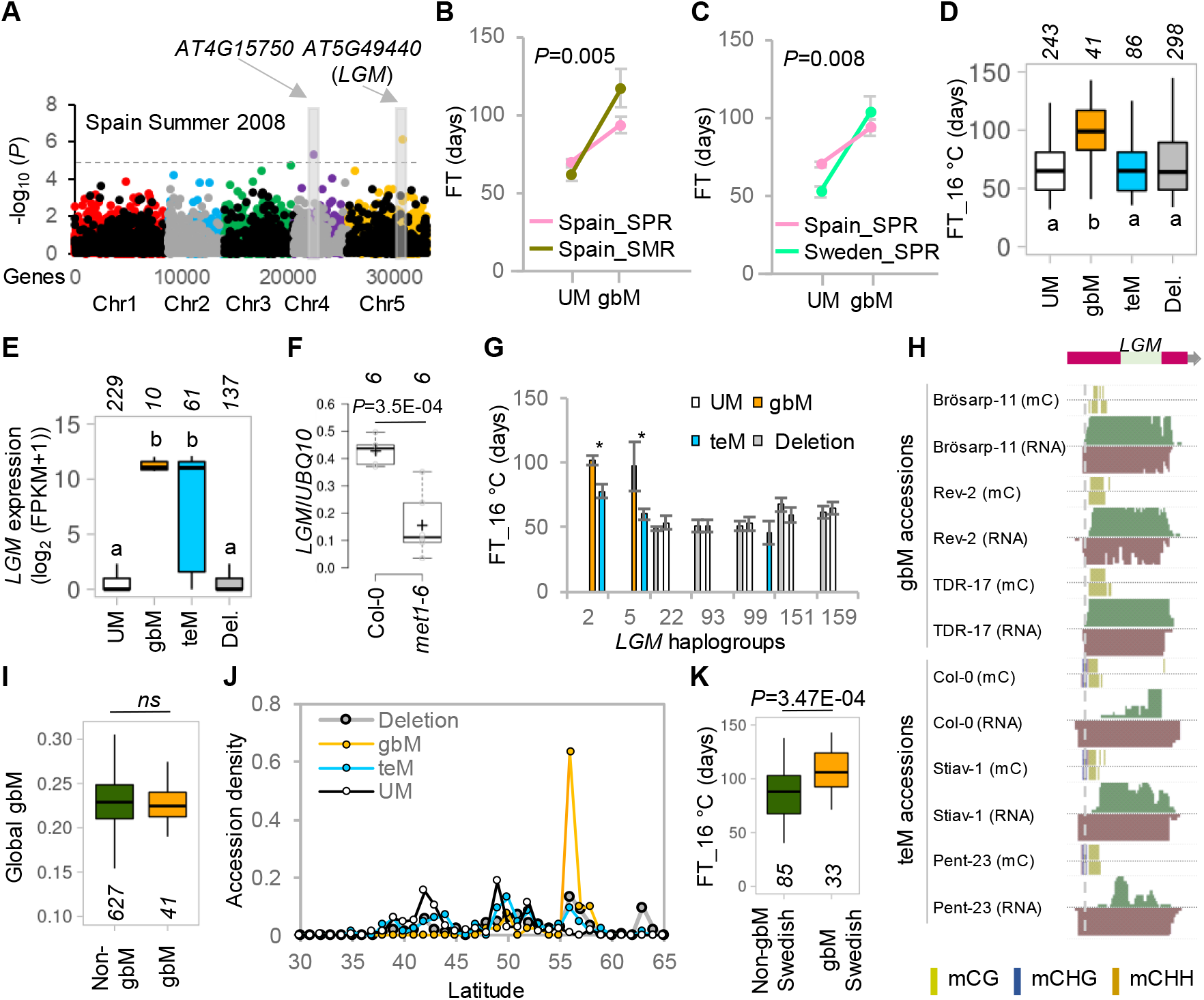
GbM in *LGM* delays flowering. (**A**) Manhattan plot of MLM epiGWA mapping for flowering time in the 2008 Spain summer planting. GbM markers are depicted with colored dots and teM markers are shown using grey and black dots. Horizontal dashed line shows 0.05 FDR. (**B** and **C**) Sensitivity of AT5G49440 (*LGM*) flowering time (FT) QTL^gbM^ to seasons (spring (SPR), summer (SMR)) in Spain (**B**) and spring climates (Spain, Sweden; **C**). *P*, two-way ANOVA. (**D** and **E**) Association of flowering (FT_16°C; **D**) and *LGM* expression (**E**) with *LGM* UM, gbM, teM, and deletion (epi)alleles in the indicated numbers of accessions. Different letters denote significant accession group differences evaluated by pairwise MLM epiGWA. (**F**) Transcript levels of *LGM* relative to *UBQ10* in Col-0 and *met1*, assessed by qRT-PCR in six biological replicates. *P*, two-tailed Student’s t test. (**G**) Effects of *LGM* gbM on FT_16° in *LGM* haplogroups. \**P*<0.05, two-tailed Student’s t-test. (**H**) AnnoJ browser snapshots of methylation along with sense (green) and antisense (brown) transcripts for representative *LGM* gbM and teM accessions. Vertical dashed line shows the long sense transcript start site. The light green box corresponds to *LGM* coding sequence. (**I**) Genome-wide average gbM levels of accessions with gbM or non-gbM *LGM*. Accession numbers are indicated. *ns* = non-significant, two-tailed Student’s t test. (**J**) Density of *LGM* UM, gbM, teM, and deletion accessions across latitude. (**K**) Association of *LGM* gbM and non-gbM (epi)alleles with FT_16°C in southern Swedish accessions. Accession numbers are indicated. *P* value corresponds to MLM epiGWA.

The *AT5G49440* fQTL^gbM^ drew our attention because it is significantly associated with flowering in six simulated climate experiments and with two flowering traits (FT_16°C and CLN) at FDR 0.05 or 0.1 (Table S14). *AT5G49440* has gbM in 6.1% of accessions, teM in 12.9%, UM in 36.4%, and is deleted in the remaining 44.6% (Figure 5D). Flowering (FT_16°C) of gbM accessions is severely delayed (by 29 days on average) compared to teM, UM or deleted accessions (Figure 5D), and hence we named *AT5G49440 Late Flowering due to Gene Body Methylation* (*LGM*). *LGM* is highly expressed in gbM and (more variably) teM accessions, whereas expression is similarly low in UM and deleted accessions (Figure 5E). Col-0 is a teM accession, and *LGM* is strongly downregulated in *met1* (which partially reduces teM; Figures 5F and S27A), indicating that intragenic methylation is required for robust *LGM* expression.

*AT5G49430*, the gene neighboring *LGM*, is a bromodomain WDR protein, a class that includes known flowering genes^107^. Therefore, the *LGM* association could arise due to effects on *AT5G49430* expression. However, *AT5G49430* is expressed similarly in accessions carrying *LGM* UM, gbM, teM, and deletion (epi)alleles, showing that *AT5G49430* does not underlie the *LGM* association (Figure S27B).

Curiously, although *LGM* is highly expressed in teM accessions (Figure 5E), we did not detect a significant association of teM with flowering time (Figure 5A), and teM, UM and deleted accessions flower similarly early (Figure 5D). Analysis within haplogroups (Table S8) revealed that gbM accessions flower significantly later in the two *LGM* haplotypes in which they are present (Figure 5G). In contrast, teM is not associated with delayed flowering in any haplotype (Figure 5G). Therefore, gbM (but not teM) at *LGM* is associated with delayed flowering independently of genetic variation.

A close inspection revealed that *LGM* is transcribed differently in gbM and teM accessions. The *LGM* locus produces abundant sense and antisense transcripts in Col-0^108^. We found that gbM accessions predominantly produce a longer sense and a shorter antisense transcript, whereas teM accessions produce shorter sense and longer antisense transcripts (Figures 5H and S27C). TeM extends further 5′ than gbM and covers the apparent transcriptional start site (TSS) of the long sense *LGM* mRNA (Figure 5H), which would be expected to silence the TSS^25^. Our results suggest that the position of gbM is crucial for producing a specific set of *LGM* transcripts that greatly delay flowering.

Global gbM levels are similar for *LGM* gbM and non-gbM accessions (Figure 5I), and gbM at *LGM* is not associated with *trans* QTLs (Figure S27D), indicating that gbM at *LGM* is evolving epigenetically. GbM accessions mainly grow in southern Sweden, around 55° latitude, where UM accessions are rare (Figures 5J, S27E, and S27F). EpiGWA analysis using only southern Swedish accessions found a significant association between gbM and flowering time (Figure 5K), showing that links between flowering time and latitude do not confound *LGM* fQTL^gbM^ detection. These results are analogous to the ones we obtained for *FLC* (Figures S24B, S24F, and S24G). Considering the much greater relative flowering delay of gbM accessions grown in simulated Sweden climate compared to Spain climate (Figure 5C), the prevalence of *LGM* gbM in southern Sweden (Figure 5J) suggests that gbM is under selection to delay flowering in this environment.

### GbM polymorphism explains natural variation in the accumulation of many minerals

GWA studies have identified numerous genes and genetic variants associated with the diversity of pant physiological traits, such as the accumulation of mineral ions^81,109,110^. Hence, we examined whether intragenic DNA methylation influences the natural variation of minerals in *Arabidopsis* leaves using published data^111^. GWA using levels of 18 minerals identified 31 associations, whereas 19 QTLs^gbM^ and 6 QTLs^teM^ were detected (Table S15). Only one of the epigenetic QTLs (HB18, associated with manganese (Mn) accumulation) colocalizes with a genetic QTL (for Rubidium (Rb) accumulation; Figure S28), indicating that – as for MLP fitness and flowering time – the effects of epigenetic and genetic polymorphism on plant traits manifest through distinct gene sets.

Next, we estimated the amount of phenotypic variance explained by QTL^gbM^ and QTL^SNP^ (the ambiguous HB18 QTL was excluded from QTL^gbM^ estimates). After accounting for SV, gbM polymorphism explained from 2.6% (arsenic; As) to 15.5% (potassium; K) of mineral accumulation variance (Figure 6A). Consistent with our results for gene expression and other phenotypes, SNP effects are comparable, explaining from 2.7% (K) to 35.3% (molybdenum; Mo) of phenotypic variance (Figure 6A). Whereas several of the genetic QTLs contain genes known to influence mineral accumulation, such as HKT1 for sodium (Na)^110^, HMA3 for cadmium (Cd)^109^, and MOT1 for Mo^112^, epiGWA identified entirely new genes (Figure S28, Table S15). After accounting for SVs, we found that 22 epigenetic QTLs still display strong associations (*P*<1E-03, MLM) between intragenic DNA methylation and ion variation (Table S15), indicating that these QTLs^gbM^ and QTLs^teM^ are not confounded by SV. Next, we analyzed 10 QTL^gbM^ genes associated with the accumulation of 4 of the most easily quantified ions – K, magnesium (Mg), Mn, and zinc (Zn) – using T-DNA insertion mutants. Plants with mutations in 9 of these genes displayed significant changes in the accumulation of relevant minerals (*p≤*0.07, 8 genes with *p≤*0.03; Figures 6B-6E). Hence, gbM polymorphism substantially explains the natural variation of physiological traits, as well as life history traits such as transition to flowering.

**Figure 6.**
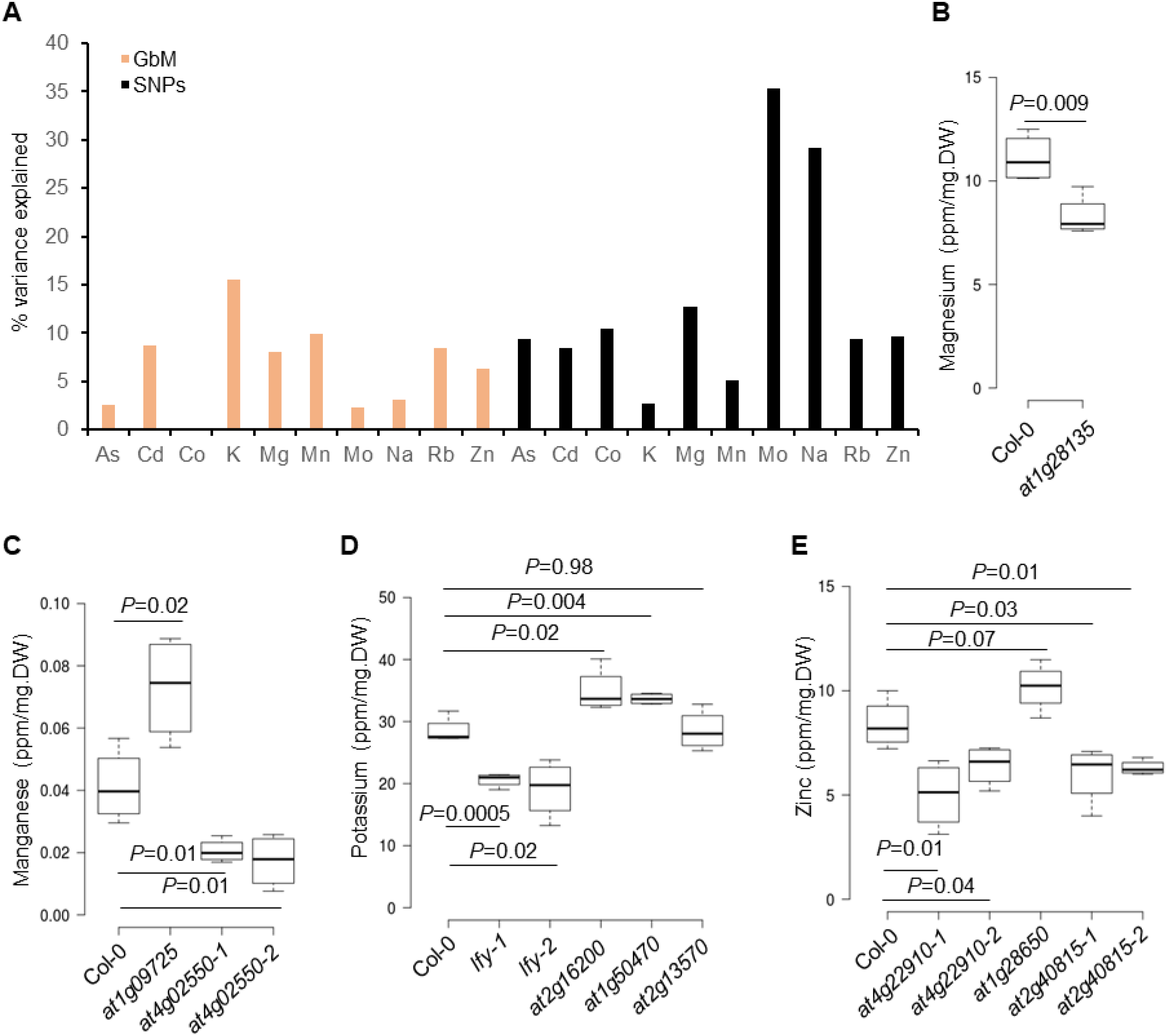
GbM affects mineral accumulation in natural *Arabidopsis* populations. **(A)** Percent variation in mineral accumulation phenotypes explained by epigenetic (gbM) and genetic (SNPs) QTLs. **(B-E)** Magnesium (**B**), manganese (**C**), potassium (**D**), and zinc (**E**) levels of Col-0 and mutants of indicated genes. Number of biological replicates is 4. *P*, two-tailed t test.

### EpiGWA directly identifies causal genes more frequently than GWA

In GWA studies, multiple *cis* SNPs are often associated with phenotypic variation due to LD between these SNPs. The associated genomic regions can be broad with multiple genes underlying QTLs^109,110,113^. SNPs that show the strongest association *P* value are frequently in non-coding regions or have unclear functional consequences^114^. Therefore, the identification of genes responsible for the relevant phenotypic variation can be challenging, particularly in the absence of candidate genes. To illustrate this point, we investigated published *Arabidopsis* GWA studies for 48 diverse traits that have 57 validated genes (Table S16) underlying SNP QTLs. We define a validated gene as one that causes the expected phenotype when mutated either in the same or other studies. By calculating the number of genes between the SNP with the lowest *P* value and the validated gene (0 means the SNP is within the validated gene), we found that the top SNP is within the validated gene in only ∼54% of cases (Figure 7A). This is likely an overestimate, because cases where researchers failed to validate an association are not always reported.

**Figure 7.**
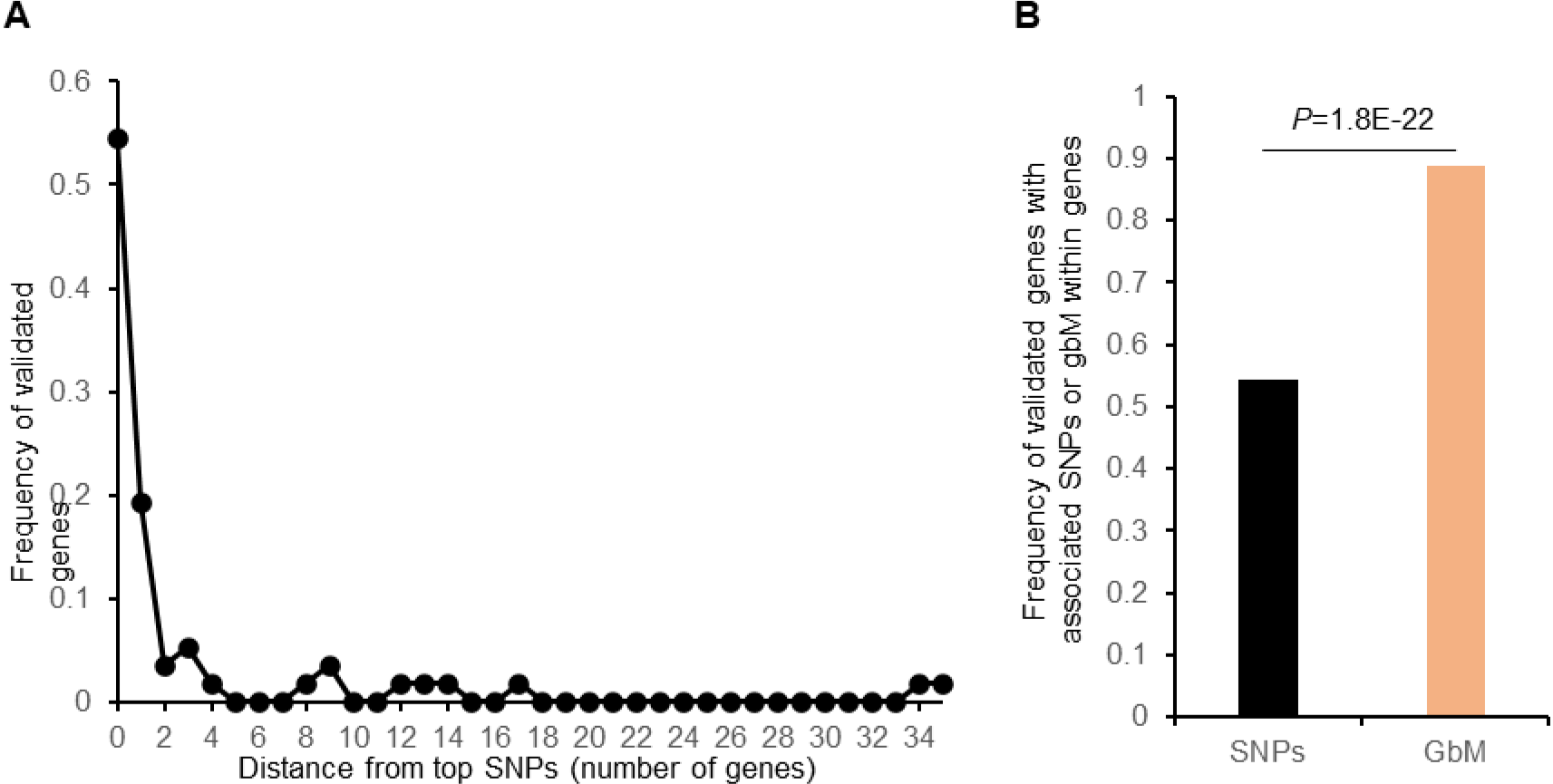
EpiGWA directly identifies causal genes more frequently than GWA. (**A**) Distribution of validated gene distances (in number of genes) from the SNP with the lowest association *P* value in GWA analysis for flowering time, ionome, root growth and development, plant regeneration, stress tolerance, seed longevity, metabolic, and DNA methylation related traits. (**B**) Frequency of SNPs or gbM variants within validated genes. For SNP-based GWA, this encompasses the intergenic region from the end of the upstream gene to the beginning of the downstream gene.

In contrast to the GWA results, we validated nearly 90% (16/18) of QTL^gbM^ using mutants of the genes that displayed associations between gbM and phenotypes, a difference that is highly statistically significant (Figure 7B). We propose that this high success rate is due to epimutation rates of gbM exceeding genetic mutation rates by about 100,000-fold^61,69,70,115^. Such high turnover should rapidly disrupt linkage between gbM patterns in nearby genes, so that only gbM in the causative gene is associated with phenotypic variance. Considering that the associations obtained with GWA and epiGWA rarely overlap (Figures S16, S22, and S28), gbM-based epiGWA may be a powerful and broadly applicable gene discovery tool, as we illustrate here by identifying 15 new genes affecting six distinct phenotypes (MLP fitness, flowering time, and accumulation of K, Mg, Mn and Zn).

### GbM variation may facilitate local adaptation

*Arabidopsis* grows in a broad range of natural environments and shows extensive local adaptation^116^. As we find that gbM affects diverse plants traits, we tested whether gbM variation may facilitate adaptation to the local environment by performing MLM epiGWA analyses using intragenic epigenetic states (UM and gbM) for 171 environmental variables^117^ (latitude, longitude, temperature, precipitation, atmospheric composition, and soil properties; Table S17). We detected 571 significant associations at FDR 0.05 between 232 genes and 115 of these variables (Table S17). SNP-based GWA analyses showed that 76.9% of epigenetic associations do not colocalize with genetic associations (Figure S29A and Table S17). GbM variation in 57 genes is associated with at least three environments (Table S17). EpiGWA *P* values for these genes are strongly correlated for associated environments (Figures 8A and 8B, and Tables S18 and S19), suggesting that multiple correlated environmental conditions impose selection on epiallelic states of individual genes.

**Figure 8.**
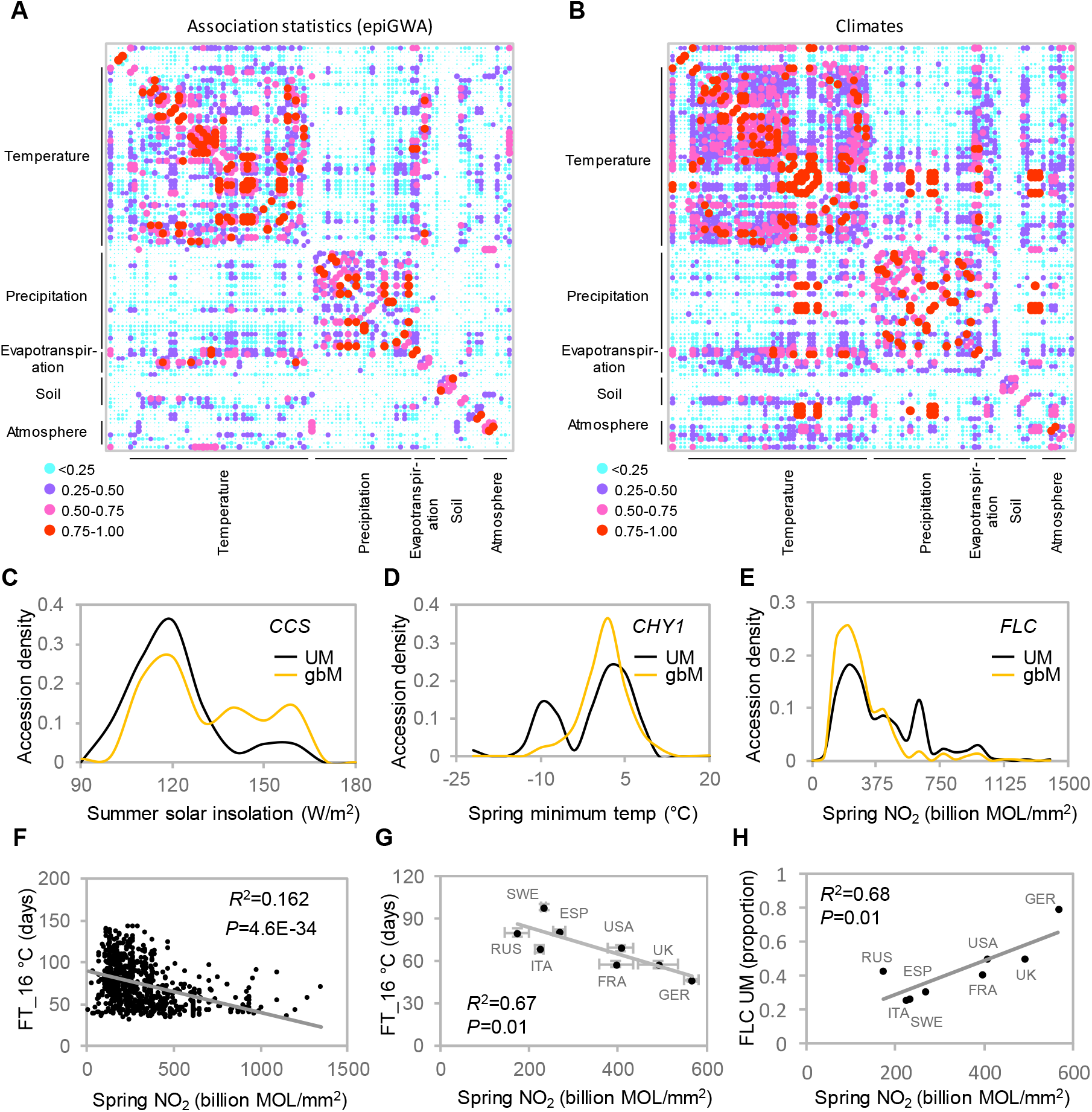
GbM variation is associated with geoclimatic variables. (**A** and **B**) Correlation (*R*^2^) matrices of epiGWA *P* values (**A**) and environmental variables (**B**) for 57 genes identified in at least three epiGWA analyses. See Tables S17 and S18 for individual environment labels in order. (**C**-**E**) Associations between gbM and environmental data for *CCS* (**C**), *CHY1* (**D**), and *FLC* (**E**). (**F**) Correlation between springtime atmospheric NO2 and flowering time (FT_ 16°C) of individual accessions. (**G**) Average (± standard error) FT_ 16°C and NO2 concentrations in Sweden (SWE), Russia (RUS), Italy (ITA), Spain (ESP), United States of America (USA), France (FRA), United Kingdom (UK), and Germany (GER). (**H**) Prevalence of *FLC* UM epiallele as a function of NO2.

Our analysis identified several notable gbM associations with a plausible functional link to environmental adaptation (Figures 8C, 8D, 8E, S29B, S29C, and S29D) that do not overlap genetic associations (Table S17). GbM in *CCS*, which mediates heat stress responses^118^, is associated with summer solar insolation, with gbM epialleles much more common in high insolation environments (Figure 8C). GbM in *CHY1*, which is involved in cold signaling and promotes freezing tolerance^119^, is associated with spring minimum temperature, with gbM epialleles rare in environments where temperature drops below -4°C (Figure 8D). GbM in *HUP9*, a regulator of flooding stress response^120^, is associated with annual precipitation (Figure S29B). *PYR1* gbM variation is associated with soil excess salts, with high salt soils almost exclusively featuring gbM epialleles (Figure S29C). *PYR1* is an abscisic acid (ABA) receptor^121^, and ABA is a central regulator of plant salt stress responses^122^. GbM variation in the calcium sensor *SOS3*^123^ associates with soil salinity and sodicity (Figure S29D), which includes measures of calcium carbonate (CaCO3) and gypsum (CaSO4·2H2O). Nearly all accessions from soils with high salinity and sodicity have UM *SOS3* epialleles (Figure S29D). These associations suggest that natural gbM variation facilitates local adaptation in native habitats.

The most striking association we discovered is between *FLC* gbM and springtime concentration of NO2, an anthropogenic atmospheric pollutant^124^, with UM *FLC* alleles prevalent in high NO2 environments (Figure 8E and Table S20). Because UM *FLC* accessions flower early (Figures 4D-4G), this association predicts that accessions from high NO2 environments should flower early. Indeed, flowering time (FT_16°C) of laboratory grown *Arabidopsis* accessions is more strongly correlated with atmospheric NO2 in native environments than with any other environmental variable, including spring frost, spring temperature, precipitation, and latitude (Figure 8F and Table S21). NO2 concentrations vary regionally, an indicator of air quality over urban and industrial centers^125^. We find that average concentrations of NO2 across countries show a remarkable linear correlation (*R*^2^=0.67) with flowering time in the laboratory (Figure 8G), suggesting that earlier flowering is advantageous in higher NO2 environments. Prevalence of the *FLC* UM epiallele in countries is also strongly correlated with NO2 (*R*^2^=0.68; Figure 8H), and enrichment of the UM allele at 48-52° latitude (Figure S24F) corresponds to Germany, which has especially high NO2 (Figures 7G and 7H). These results suggest *FLC* gbM variation is selected to adapt flowering time to atmospheric NO2 (or an unevaluated environmental factor correlated with NO2).

## Discussion

Our results demonstrate that gbM promotes gene expression (Figure 2E-H), and that its natural variation influences at least six important traits: fitness in a hot and dry climate, flowering time, and accumulation of K, Mg, Mn and Zn (Figures 3-6). Notably, epigenetically inherited intragenic methylation accounts for a substantial amount of gene expression and trait variance (Figures 2A, 2C, 3E, 4C, 6A, S9A and S9B). Although estimation of phenotypic variance effects is challenging, our ability to consistently identify and validate associations between methylation and phenotypic variance (Figures 3F-3I, 4B, 6B-6E, and 7B) indicates that the phenotypic effects of intergenic methylation are comparable to those of SNPs.

When assessing the effects of epigenetic polymorphism, as we do here, confounding effects of *cis* or *trans* genetic polymorphism must be ruled out. We demonstrate that neither known *cis* SNPs nor SVs can explain our results (Figures 1G-1L, 3E, 4C, and 6A), which leaves undetected cryptic polymorphism. However, to explain our results, *cis* genetic polymorphism must have certain characteristics. First, it must be common in the population, because we discard rare alleles in epiGWA analyses based on epiallelic states. Second, it must be systematically linked with methylation variants but not with known genetic polymorphism (either SNPs or SV), because the vast majority of the epigenetic effects we identify are not confounded by known genetic variation (Figures 1G-1L, 3E, 4C, and 6A), and because epiGWA directly identifies the relevant gene more frequently than GWA (Figure 7B), which is only possible if the causative variant isn’t linked with nearby genetic polymorphism. We are not aware of any genetic polymorphism type that fits this description: common polymorphism that is systematically unlinked with nearby SNPs or SVs. Furthermore, lack of linkage with genetic polymorphism, but linkage with gbM polymorphism that evolves ∼100,000 times faster^61,69,70,115^ isn’t theoretically plausible. Thus, an entirely new and highly implausible type of cryptic genetic polymorphism would have to be invoked to explain our results, whereas the rapid turnover of gbM naturally explains the lack of linkage with genetic variation. Furthermore, only a tiny fraction of gbM polymorphism we consider here is explained by *trans* variation (Figures 1L, 3E, 4C, S24C, and S27D), and the reliable verification of phenotypic associations with local gbM polymorphism (Figures 3F-3I, 4B, 6B-6E, S17A, and S17B) indicates that these effects cannot be explained by cryptic *trans* variation that functions independently of gbM. Overall, our results are entirely consistent with published work that shows local gbM variation is primarily epigenetic in *Arabidopsis*^36,61^.

The epigenetic effects on plant phenotypes we describe here are primarily mediated by gbM (Figures 3A, S20 and S25 and Tables S13-S15). Thus, although studies of epigenetic trait inheritance have focused exclusively on TE methylation or teM^11,22,32,62,126^, our results suggest that gbM variation is the epigenetic mechanism primarily responsible for driving phenotypic variation in natural plant populations. Furthermore, the epigenetic and genetic effects on gene expression and phenotype manifest through distinct sets of genes, or at least their relative strengths at individual loci are sufficiently different for epiGWA to detect 16 new functional genes in this study and new phenotypic associations with the environment (such as flowering time and NO2). This, and the propensity of epiGWA to directly identify the relevant gene (Figure 7B), make epiGWA a powerful gene discovery tool. We therefore expect epiGWA to reveal new genes that regulate key processes across a wide range of species, including major crops. QTLs controlling less than 10% of phenotypic variance have been successfully exploited to increase rice yield^127–130^, suggesting that gbM polymorphism could prove useful for crop improvement.

DNA methylation is mutagenic^131^, and its presence in coding sequences likely incurs a fitness cost^39^. The widespread conservation of gbM in plants and animals has therefore presented a mystery. A potential explanation is that gbM allows rapid adaptation to new or changing environments, because gbM epimutation rates are about 10^5^ times greater than genetic mutation rates^61,69,70,115^. The direct relationship between gbM and mRNA levels (Figures 1E, 2F, and S2) also indicates that gbM changes should quantitatively alter gene expression, whereas most mutations are expected to be neutral or detrimental to gene function^132^. Thus, gbM variation within a population can rapidly generate a range of gene expression for selection by the environment. This argument implies that the relative gbM contribution to phenotypic variance should decrease with divergence time, as genetic differences accumulate and exert an ever-larger effect. Indeed, mathematical modelling indicates that epigenetic fluctuations that generate local gbM variation operate on thousand-year timescales, whereas overall gbM patterns over longer timescales are genetically determined^61^. Hence, at long timescales, such as those that typically separate species, genetic differences should dominate. However, our results indicate that within the *Arabidopsis* population, epigenetic gbM variation is of comparable importance to genetic variation for driving phenotypic diversity.

The association between atmospheric NO2, flowering time and *FLC* gbM (Figures 8E-8H) presents a plausible illustration of how gbM variation may facilitate adaptation to a rapidly changing climate. The strong negative association between springtime atmospheric NO2 in native environments and flowering time under laboratory conditions (Figures 8F and 8G) suggests that accelerated flowering is adaptive under elevated NO2. Natural genetic variation at *FLC* is a major determinant of flowering time^88,96,97^, and is associated with over twenty environmental variables that are (or may be) related to flowering, including latitude, temperature and precipitation, but not NO2^117^. The majority of atmospheric NO2 (>75%) is produced by recent human activities, especially the burning of fossil fuel^124^. Therefore, *Arabidopsis* populations have had to adapt to NO2 concentrations (or a correlated unexamined environmental variable) changing over a few decades. Genetic adaptation at *FLC* apparently has not yet occurred in response to such rapid environmental alteration, or at least is too weak for detection by GWA analysis. However, epigenetic gbM variation at *FLC* is significantly associated with atmospheric NO2 (Figures 8E-8H), but not other environmental variables (Table S20), which is consistent with our observation that *FLC* gbM varies independently of DNA sequence (Figure 4G). Therefore, gbM variation at *FLC* has likely facilitated adaptation to anthropogenic NO2 increases, whereas genetic variation has been involved in adaptation to environmental conditions that vary over longer timescales. This interplay between epigenetic and genetic adaptation is consistent with evolutionary models^8–11^ and may be a generally important component of environmental adaptation.

Although this study investigates plant epigenetic inheritance, gbM variation is likely to influence population dynamics in many animal lineages. Direct comparison of plant gbM with vertebrate methylation is challenging because vertebrates have broadly methylated genomes in which methylation is the default state^133^. Furthermore, methylation is extensively reprogrammed during mammalian embryogenesis and germline development^134^, which precludes methylation inheritance at most loci^135–137^. However, many invertebrates, from basal cnidarians to chordates, display gbM patterns highly similar to those of plants^23–25,39^. There are also intriguing indications that invertebrate methylation is epigenetically heritable. For example, a cross between two wasp species produces hybrids with chromosomes that retain their ancestral methylation patterns^138^, as would be expected if methylation is epigenetically inherited. Moreover, methylation fidelity is high in honeybee sperm compared to somatic tissues ^52^, and methylation patterns can be stably transmitted from fathers to daughters^139^. These features of invertebrate methylation suggest that epigenetically heritable gbM facilitates environmental adaptation in animals. This may be especially important for species like aphids^140^, in which self-compatibility or asexual reproduction limit the generation of genetic diversity through outcrossing. By rapidly creating functional diversity, gbM may allow animal (and plant) evolution to explore a broader range of reproductive strategies.

## Acknowledgements

We thank P. Baduel and V. Colot for sharing SV data, A. Muyle for gbM conservation data, and X. Feng, C. Dean, E. Coen and Zilberman lab members for constructive comments on the manuscript. This work was supported by a European Research Council grant (725746) to DZ. This study would not have been possible without *Arabidopsis* 1001 genome, methylome, and transcriptome resources.

## Ethics declarations

The authors declare no competing interest.

**Supplementary Figures:** This file contains material and methods and supplementary figures.

**Supplementary Tables:** This file contains supplementary tables 1-24.

