## Supplementary Information for "Epigenetic inheritance mediates phenotypic diversity in natural populations"

This file contains materials and methods and figures S1-S29.

#### Materials and methods

##### Methyl-C seq data analysis

Bisulphite sequence reads were accessed for the 1001 methylomes[1] experiments from the Sequence Read Archive (SRA). Sequencing reads of 948 non-redundant *Arabidopsis* accessions were aligned to the *Arabidopsis* TAIR10 genome reference sequence[2], using BSMAP[3] with default parameters, and known SNPs and indels[4] were masked. Genes and transposons were annotated using the Araport11 annotation[5]. Methylomes were segmented into unmethylated (UM), gene body methylated (gbM) and TE-like methylated (teM) segments as previously described[6]. Methylation status of each individual CG site was called by comparing the counts of aligned reads indicating methylated and unmethylated status at the site. Fisher's Exact test was used to determine whether there was sufficient read coverage at the site to distinguish the site from a fully unmethylated site with an error rate similar to the methylation rate observed in the chloroplast of the sample in question (as an estimate of bisulphite conversion inefficiency), or from a fully methylated site with a similar error rate. For sites where these tests indicated coverage was sufficient, a binomial test was used to identify sites with significantly more methylated reads than expected at an unmethylated site. Sites with significantly more methylated reads than would be expected for an unmethylated site, but with less than 45% reads methylated, were classified as partially methylated, and generally treated as missing data. Genes' methylation status in each accession was classified as gbM, teM, both (gbM and teM), UM, or indeterminate, based on overlapping methylome segments. Genes overlapped by a gbM segment 3 or more CG sites long, with at least one CG site being called as methylated by binomial test, were classed as gbM genes, unless they are also overlapped by a TE-like methylation (non-CG methylation) segment at least 0.25 as long as the gbM segment, in which case they were classified as both. Genes overlapped by a teM segment 3 or more CG sites long were classified as teM genes, unless they are also overlapped by a gbM segment at least 0.25 as long as the teM segment, in which case they were classified as both. Genes which are not overlapped by either gbM or teM segments and which span at least 3 sites called as unmethylated by binomial test, were classified as UM. The remainder of genes were classified as indeterminate. The mean CG methylation level of gbM or teM genes was assessed for each gene by summing the number of spanned CG sites called as methylated and dividing by the number of spanned CG sites called as either methylated or unmethylated by binomial test.

##### Estimation of prevalence of teM across gbM conservation bins

The number of genes having gbM or teM epigenetic states was determined in 948 *Arabidopsis* accessions. Pearson's correlation analysis for the number of gbM and teM genes was performed using accessions with more than 60% sequencing coverage of genomes. Conservation of epiallelic states of genes was analyzed as a fraction of accessions having gbM or teM and the total available calls. Average prevalence of teM within gbM conservation bins was estimated in four gbM categories (0, >0<10%, 10-90%, and >90%), decile gbM bins, and percentile gbM bins. To compare our results with published findings, identical analyses were performed using available data[7] with restrictive definitions of gbM and teM.

##### Associations of intragenic DNA methylation with expression levels

RNA sequencing data for 625 *Arabidopsis* accessions with information of gene specific mCG levels were retrieved from GEO: GSE80744[1]. Genes showing no detectable expression in leaves of any of these accessions were discarded from association analyses. Furthermore, to avoid confounding by low allele frequencies these analyses were performed using gbM and teM genes having at least one mCG site in more than 10% *Arabidopsis* accessions. This allowed us to examine associations between mCG levels and gene expression for 18,679 gbM and 1,442 teM genes. Expression levels of genes were regressed on mCG levels in a linear model. Association *P*-values for Pearson correlation were estimated using SigmaPlot 14.0.

Bonferroni (α=0.05), or 0.05 and 0.1 false discovery rate[8] (FDR) corrections were implemented to account for multiple tests. The percentage of expression variance explained by intragenic DNA methylation was calculated as

$$\text{PVE} = \frac{(\beta)^2 (VmCG)}{VP}$$

where $V_{mCG}$ is the variance of mCG, $V_P$ corresponds to phenotypic (expression) variance, and $\beta$ effects for each association test were calculated as;

$$\beta = R \text{ x } \left( \frac{\sigma P}{\sigma mCG} \right)$$

where $R$ is Pearson correlation coefficient, $\sigma P$ corresponds to standard deviation of gene expression and $\sigma mCG$ is standard deviation of mCG in the population.

##### Gene feature annotation

Numbers of CG (CGG or CGT or CGC or CGA) sites were enumerated by scanning annotated genes[5] within the Col-0 reference sequence[2] with a three base window and step size of one base. Gene lengths were obtained from the Col-0 annotation[5]. Then, CG dinucleotide frequencies were calculated by normalizing the number of CG sites to a gene's annotated length. Mean expression level of each gene was calculated across 625 accessions.

##### Pipeline to account for SNP effects on the expression of eQTL$^{gbM/teM}$ genes

To disentangle the effects of intragenic methylation on expression from *cis*-acting DNA sequence changes, we performed GWA analyses for the expression of 765 eQTL$^{gbM}$ and 217 eQTL$^{teM}$ Bonferroni genes using 1001 genomes SNP[4] data in an accelerated mixed model[9]. Colocalization of each *cis* eQTL (eQTL$^{SNP}$) significant at Bonferroni threshold ($\alpha$=0.05) with epigenetic eQTL was determined. The eQTL$^{gbM/teM}$ genes for which no colocalized *cis* eQTL$^{SNP}$ were detected are considered to affect gene expression variation independently of genetic variation (retained eQTL$^{gbM/teM}$) (Figures 1G, 1H, and S6). In cases where eQTLs$^{gbM/teM}$ colocalized with eQTLs$^{SNP}$, the original population of accessions was separated into two nested populations, each fixed for the GWA SNP (Figure S6). Associations between intragenic DNA methylation and expression of these genes were re-examined within nested populations to account for the effects of SNP variation on expression. The genes that exhibited significant association between intragenic DNA methylation and expression in at least one nested population were also classified as retained eQTL$^{gbM/teM}$. Genes without significant associations between intragenic DNA methylation and gene expression in nested populations were considered likely confounded by linked SNPs in the population. Accordingly, these eQTL$^{gbM/teM}$ were classified as lost eQTL$^{gbM/teM}$ genes. To account for GWA SNP effects on expression variance, percent variance explained by methylation was calculated in nested populations as described above.

##### Haplotype analyses

To account for allelic heterogeneity association between methylation and expression were examined within haplotypes. SNPs within, and 4 kb upstream and downstream of genes were extracted from a preimputation version of the 1001 genome SNP panel[4]. Sequences were aligned, and the accessions invariant for SNPs over the entire region for each gene were classified into a haplogroup. Haplogroups comprising less than 15 accessions were discarded from association analyses to have a reasonable number of accessions to perform association analyses. Associations of mCG with gene expression or phenotypes were examined within haplogroups to fully account for the effects of local SNP variation on expression or phenotypic variation.

#### Accounting for structural variation effects on epigenetic QTLs

Structural variants were identified within epigenetic QTLs and 4 kb upstream and downstream using published TE polymorphism data in *Arabidopsis* accessions[10]. Associations between structural polymorphism and expression were examined using a linear model. To account for the effects of structural variants on epigenetic QTLs, we analyzed associations between methylation and phenotypes (expression, relative fitness, and flowering time) in populations without structural polymorphisms.

#### Epigenome-wide association studies for relative fitness

EpiGWA analyses for relative fitness were performed using published relative fitness data[11] of 412 *Arabidopsis* accessions with sufficient mCG information. Common garden experiments had been performed in two climatically distinct field stations in Madrid (M) and Tübingen (T)[11]. Madrid presents a climate that transitions between Mediterranean and semi-arid climates and Tübingen is characterized by a temperate climate with no dry season and warm summers. High (H) and low (L) rainfall conditions typical of Tübingen and Madrid had been simulated during these experiments. To mimic low and high-density populations in nature, individual (I) or multiple plants (P) had been grown in pots. EpiGWA analyses were performed using a linear model to assess associations between gbM or teM levels of genes and relative fitness. For these analyses we focused on genes having gbM or teM conserved in more than 10% of *Arabidopsis* accessions. Linear model association mapping analyses may detect excessive significant marker-trait associations due to underlying population structure[12]. We however detected only two associations (*PROT1* and AT1G19410) at 0.05 FDR for relative fitness in MLP (Table S22). And, gbM variation in one gene *MuDR* (AT1G64255) is associated with relative fitness in MLI at 0.1 FDR. We next used quantile-quantile (QQ) plots and genomic control (GC) inflation factor λ[13] to assess confounding of association statistics (Figure S14 and Table S6). λ was calculated using unlinked markers as

$$\lambda = \frac{Median\ X^2\ observed\ P}{Median\ X^2\ expected\ P}$$

where $X^2$ is the chi-square and $P$ is the $P$ value.

λ varied between phenotypes and ranged from 0.91 (relative fitness MHP) to 1.48 (relative fitness TLI) (Table S6). To control confounding effects of population stratification, association statistics were corrected using λ and genome-wide significance threshold was recalculated using corrected $P$ values. Both *PROT1* and AT1G19410 associations were significant at 0.05 FDR, however, *MuDR* was not significant at 0.1 FDR. Associations between intragenic DNA methylation and fitness significant at 0.05 FDR[8] are called epigenetic QTLs during this study. Tripartite associations between mCG levels, gene expression, and relative fitness in MLP for *PROT1* and AT1G19410 were analyzed using a linear model. The percentage of relative fitness variance in MLP that can be explained by epigenetic QTLs was calculated as described above for DNA methylation-expression association analyses.

#### Epigenome-wide association studies for flowering related traits

Three types of epiGWA mapping were performed for flowering related traits to identify the best model to account for confounding effects of population structure. A linear model was employed using mCG levels of genes, and two models, generalized linear model (GLM) and mixed linear model (MLM), were used for epiGWA using epiallelic states (UM or gbM; UM or teM) of genes. The methods for determination of epiallelic states of genes are described in Methyl-C seq data analysis section. The number of *Arabidopsis* accessions used for these epiGWA analyses are listed in Table S23.

Linear model epiGWA mapping was performed to examine associations between mCG levels of genes (>10% gbM or teM conservation) and flowering time data (flowering time at 10 °C (FT_10°C) and 16 °C (FT_16°C))[4]. Association statistics for these epiGWA analyses were highly confounded (λ=4.50 for FT_10°C and λ=4.52 for FT_16°C; Figure S19 and Table S10). Around 7,500 genes showed significant associations between mCG levels and flowering time at 0.05 FDR (Figure S19). Applying uniform λ correction for association $P$ values in such cases is unsatisfactory for correcting population structure at genes with strong differences in mCG levels across subpopulations and can also result in a loss of statistical power at genes with uniformly distributed mCG levels[14,15]. Given the correlation of flowering with geographic regions, similar confounding of association statistics has been reported for flowering related traits in *Arabidopsis* GWA studies[12]. Strong confounding of $P$ values renders linear model epiGWA using mCG levels inappropriate for association mapping in structured populations.

Next, we used binary epiallelic states of genes to perform GLM and MLM epiGWA mapping using FT_10°C and FT_16°C flowering time phenotypes and seven additional flowering related phenotypes[16] (number of days for inflorescence stalk to reach 1 cm (DTF2), number of days to the opening of first flower (DTF3), number of cauline leaves (CLN), number of rosette leaves (RLN), cauline branch number (CBN), primary number of inflorescence branches (RBN), and length of primary inflorescence stalk). GLM implemented in TASSEL [17] is a fixed effects linear model that we used to test associations between epiallelic states and phenotypes. Association $P$ values for several of the flowering phenotypes deviated significantly from expected distribution of $P$ values as evidenced by QQ plots and $\lambda$ estimates (Figure S20 and Table S11). Hence, GLM using epiallelic states is also inappropriate for epiGWA mapping in structured populations. Next, a MLM[17] that includes both fixed and random effects was used to correct population structure. MLM can be presented as

$$Y = \beta X + Zu + e$$

where $Y$ represents the vector of phenotypes, $\beta$ denotes the vector containing fixed effects including genetic markers and population structure (Q-matrix), $u$ captures variance due to relatedness between individuals (kinship (K) matrix), $X$ and $Z$ are the design matrices, and $e$ captures variance due to the environment. The Q-matrix of population membership estimates was derived from principal component analysis of epiallelic states. The K matrix accounts for epigenome-wide patterns of relatedness between the individuals and was estimated using the identity-by-state method[17]. QQ plots and $\lambda$ estimates based on MLM epiGWA showed no significant deviation of distribution of association $P$ values from null distributions (Figure S20 and Table S11). MLM was thus used to dissect the epigenetic architecture of flowering related phenotypes. Genes having methylation calls in <10% accessions were removed. The percentage of phenotypic variation explained by significant epigenetic QTLs at 0.05 FDR with minimum allele frequency (MAF) >5% was calculated using TASSEL[17] as the ratio of sum of squares (SS) of epiallelic markers (after fitting all other model terms) to total SS.

Association between epiallelic states and expression levels of eight flowering epiQTL genes was analyzed using MLM epiGWA mapping. To examine associations between gene expression and phenotypes, flowering phenotypes were regressed on quantitative variation of gene expression in a linear model. Associations between epiallelic states and flowering or gene expression phenotypes in nested, regional, or allelic populations of *FLC*, *FRI*, and *LGM* were tested using MLM epiGWA analyses.

#### Epigenome-wide association studies for leaf ionome

Data for accumulation levels of 18 mineral elements[18] in leaves of 934 *Arabidopsis* accessions were used for epiGWA analyses to identify gbM and teM variants associated with the diversity of these traits. EpiGWA analyses were performed using MLM implemented in Tassel[17] as described above. We filtered out rare (MAF<5%) gbM and teM variants. FDR 0.05 correction[8] was implemented to account for multiple tests and identify significant associations.

#### Epigenome-wide association studies for geoclimatic variables

Data for 171 geoclimatic variables[19] were used for epiGWA analyses to identify gbM variants associated with environmental variation in the native range of *Arabidopsis* accessions. EpiGWA analyses were performed using MLM implemented in Tassel[17] as described above. We filtered out rare (MAF<5%) gbM variants. FDR 0.05 correction[8] was implemented to account for multiple tests and identify significant associations.

#### Density of accessions across latitude or environmental variables

Latitude coordinates of accessions were downloaded from the Arapheno database[20]. Density of *FLC* UM and gbM epialleles was calculated in a window of 1° latitude. Density of epiallelic forms is the fraction of accessions with an epiallele in 1° latitude within all accessions with that epiallele across the geographic range. Density of *FLC*, *CHY1*, *CCS HUP9*, *SOS3*, and *PYR1* UM and gbM accessions was determined across environments using a similar approach. Density of *LGM* UM, gbM, teM, and deletion epi(alleles) across latitude was determined in a 1° window.

#### Genome-wide association studies for relative fitness, flowering, and ionome phenotypes

GWA analyses were performed for relative fitness in eight climates[11],nine flowering related phenotypes[4,16], and levels of 18 minerals[18] using the same accessions as for epiGWA analyses. GWA mapping was carried out using 1001 genomes SNP data[4] with an accelerated mixed model (AMM)[9] implemented in PyGWAS: Python library for running GWAS (version 1.7.4). AMM has been shown to work well in previous studies for flowering and other phenotypes[9,21,22]. Single nucleotide polymorphisms (SNPs) with MAF > 5% in the population were considered. 0.05 FDR correction[8] was implemented to account for multiple tests and identify genetic QTLs. The percentage of phenotypic variance explained by significant genetic

QTLs was estimated as described above (SS of SNP markers (after fitting all other model terms)/total SS).

#### Genome-wide association to account for effects of *trans* QTLs on methylation variation

GWA analyses were performed for mCG levels of retained Bonferroni eQTL$^{gbM/teM}$ and fitness associated epigenetic QTLs, and for epiallelic states of flowering epigenetic QTL genes. GWA mapping was carried out as described above to identify *trans* genetic QTLs which are significant at Bonferroni threshold. These analyses were performed in three *Arabidopsis* populations; worldwide populations that we used for association mapping for gene expression and phenotypes, 133 accessions of the Swedish panel[23], and a random non-Swedish worldwide population of equal size to the Swedish panel. The percentage of mCG or epigenetic state variance explained by *trans* genetic QTLs was estimated as the ratio of SS of SNP markers (after fitting all other model terms) to total SS. Percent phenotypic (fitness in MLP and flowering related traits) variance explained by epigenetic variation is presented before and after accounting for the effects of *trans* QTLs on methylation variation.

#### RNA and bisulfite sequencing analysis

Total RNA was extracted from 4-week-old *h1-/-* and *h1-/-;met+/-* leaves using Trizol (Invitrogen, cat. No. 15596026). To remove genomic DNA from samples, 1 mg of RNA was treated with the DNA-free DNA removal kit (Thermo, AM1907). 100 ng of gDNA-depleted total RNA was used to construct RNA sequencing libraries with Ovation RNA-seq systems 1-16 for model organism (Arabidopsis) (Nugen, cat. No. 0351). To investigate the association of intragenic DNA methylation with expression level in *h1-/-;met1+/-* plants, we first defined demethylated gbM genes as ones with more than 10% CG methylation, lose more than 5% CG methylation in *h1-/-;met1+/-* vs. *h1-/-* plants, and have less than 5% CG methylation in *h1-/-;met1+/-*. The gene expression fold change in *h1-/-;met1+/-* plants (vs. *h1-/-* plants) was calculated using DeSeq2[24]. To analyze the association between gene expression and gbM change, we compared average expression fold change of demethylated gbM genes and gbM genes that retain intragenic DNA methylation *in h1-/-;met1+/-* plants.

For *h1-/-;met1+/+* plants isolated from segregating *h1-/-;met1+/-*, 100 to 700 ng of DNA-depleted leaf RNA was used to construct RNA sequencing libraries (Illumina, cat. No. 20020610 and 20019792) following the manufacturer's manual. Since segregating plants showed aberrant non-CG hypermethylation over gbM genes, we filtered out genes that gain non-CG methylation (average mCHG or mCHH>0.01). GbM genes that either lose or keep methylation were identified as described for *h1-/-;met1+/-*.

For bisulfite sequencing analysis of *h1-/-*;*met1*+/+ plants, we extracted genomic DNA (gDNA) from 4-5-week-old plant leaves. 500 ng gDNA was sheared to 100-1000 bp using Bioruptor Pico (Diagenode). gDNA libraries were constructed using NEBNext Ultra II DNA library prep kit for Illumina (New England Biolabs, cat. No. E7645). We performed bisulfite conversion twice (QIAGEN, cat. No. 59104) with ligated libraries and amplified libraries by PCR. Sequenced reads were mapped with the bs-sequel pipeline (https://zilbermanlab.net/tools/).

RNA-seq and DNA methylation data are deposited in GEO with accession GSE183785.

#### Quantitative real-time PCR

Transcript levels of *PROT1*, *FRI*, and AT5G49440 (*LGM*) were quantified in Col-0 and *met1-6*[25] with plants grown in a chamber with cycles of 16 h light (120 µE $m^{-2}$ $s^{-1}$) at 27 °C day and 16 °C night temperatures without humidity control and shoots of 3-week-old plants were harvested. Each sample was a pool of 5 plant shoots, and samples were harvested from six independent experiment. For quantification of AT1G19410 (*ANH*) mRNA levels, Col-0 and *ddcc*[26] plants were grown for 10 days as described above, then a 12 h cold treatment (4 °C) was applied to induce and detect the expression of *ANH* [27]. *ANH* transcript abundance was analyzed from five independent experiments with 25 plant shoots pooled per experiment. Total RNA was extracted using the SV Total RNA Isolation System (Promega, cat#Z3101). One µg of total RNA was used for first strand cDNA synthesis using SuperScript™ IV Reverse Transcriptase (Invitrogen, 18090050) and Oligo(dT)15 Primer (Invitrogen, 18418012) in a final volume of 25 µL, according to the manufacturer's instructions. For qRT-PCR, 25 ng of first strand cDNA was used as template. qRT-PCR was performed in triplicate using the CFX Connect Real Time PCR Detection System (Bio-Rad). cDNA amplification was monitored using SensiFAST SYBR No-ROX One-Step Kit (Biolone, Bio-72005) at an annealing temperature of 60 °C. *UBQ10* (AT4G05320) was used as an internal control. The primer sequences used for the analysis of *PROT1*, *FRI*, *LGM*, *ANH* and *UBQ10* are listed in Table S24. Relative transcript levels of genes of interest (GOI) compared to *UBQ10* were determined using the equation $RTL = [(E)^{-Ct}]^{GOI} / [(E)^{-Ct}]^{UBQ10}$.

#### Quantification of minerals in plant samples

Wild-type and mutant plants were grown in four biological replicates to analyze the accumulation of minerals in the shoots. Oven dried samples (~15 mg) were placed in a vessel (Environmental Express® cat#SC415) with 1 mL of nitric acid 65% (EMD Millipore cat #1.00456.2500) and hydrogen peroxide 30 % (Sigma-Aldrich cat #H3410-1L) and left at room temperature overnight. The samples were then digested using an Environmental Express® Hotblock digestion system (cat #SC196) set at 80˚C for 8 hours. Microwave-induced plasma optical emission spectrometer 4210 (MP-AES Agilent Technologies) coupled with an autosampler SPS4 (Agilent Technologies) was used to quantify K, Mg, Mn, and Zn at 769.897 nm, 280.271 nm, 4003.076 nm, and 202.548 nm, respectively. Standard curves for each element were used to determine mineral concentrations in samples.

#### Plant materials, growth conditions, and phenotyping

For relative fitness phenotyping under drought and heat stress, seeds of *Arabidopsis* Col-0 accessions and homozygous T-DNA insertion mutant lines for *PROT1* (*prot1-1*; SALK_030711C and *prot1-2*; SALK_018050C) and *ANH* (*anh-1*; SALK_098287C and *anh-2*; SALK_036488C) were obtained from Nottingham Arabidopsis Stock Centre (NASC). Seeds were stratified at 4 °C for seven days and germinated in 9 cm pots containing vermiculite. Each pot contained 4 plants. Plants were grown in a chamber with cycles of 16 h light (120 μE $m^{-2}$ $s^{-1}$) and 8 h dark, with 16 °C night and 27 °C day temperatures to induce heat stress. For well-watered conditions soil water content (SWC) was maintained at 60%, and 25% SWC was used for drought stress. Each pot was weighed daily to adjust SWC. Survival to fruit for Col-0 wild type plants and *prot1* and *anh* mutant plants was scored before harvesting under heat or joint heat and drought stress. Number of seeds produced by surviving plants was recorded as a measure of fecundity. Fitness of each genotype under heat or combined heat and drought stress was calculated as a product of percent survival and average fecundity during each experiment. Relative fitness of *prot1* and *anh* was estimated with respect to the average fitness of Col-0 within each condition. To understand the phenotypes that could contribute to differences in relative fitness of *prot1* and *anh* mutant plants, the three genotypes were phenotyped for shoot biomass and fertility. Shoot biomass for Col-0, *prot1* and *anh* plants was measured as shoot dry weight at maturity. Fertility was scored as a percentage of flowers producing siliques.

For flowering time phenotyping, seeds of *Arabidopsis* Col-0 accessions and homozygous T-DNA insertion mutant lines *AT1G51820* (*at1g51820*-1; SALK_208927 and *at1g51820-2*;

SALK_055952), *AT1G18210* (*at1g18210-1*; GABI_826B09 and *at1g18210-2*; SALK_075633), *AT3G43860* (*at3g43860-1*; SALK_201540 and *at3g43860-1*; GABI_129G07), *AT3G09530* (*at3g09530-1*; SALK_034560 and *at3g09530-2*; SALK_023893), *AT1G26795* (*at1g26795-1*; SALK_124311 and *at1g26795-1*; SALK_124319), and *AT4G33560* (*at4g33560-1*; SALK_133653) were obtained from Nottingham Arabidopsis Stock Centre (NASC). T-DNA insertion mutant lines (AT1G09725 (at1g09725; CS821762), AT4G18370 (at4g18370-1; SALK_099162C and at4g18370-2; SALK_036606C), AT4G02550 (at4g02550-1; SALK_136283C and at4g02550-2; SALK_028806C), AT1G70920 (at1g70920; CS863888), AT5G61850 (lfy-1 and lfy-9), AT2G16200 (at2g16200; SALK_082813), AT1G50470 (at1g50470; SALK_200371C), AT2G13570 (at2g13570; SALK_085886C), AT4G22910 (at4g22910-1; SALK_083656C and at4g22910-2; SALK_101689C), AT1G28650 (at1g28650; SALK_010911C); AT2G40815 ( at2g40815-1; SAIL_138_E02 and at2g40815-2; SALK_023214C), and AT1G28135 (at1g28135; SALK_017094)) for ionome analysis were obtained from Arabidopsis Biological Resource Center (ABRC). Seeds were stratified at 4 °C for seven days and germinated in 9 cm pots containing vermiculite with each pot containing three plants. Plants were grown in a chamber with cycles of 16 h light (120 µE m−2 s−1) and 8 h dark, with 16 °C constant temperature. Flowering time of each genotype was scored as number of days to the appearance of first flower.

#### Statistical analyses

Statistical significance was assessed using appropriate tests, including Student's t test, Wilcoxon rank sum test, one-way ANOVA, or two-way ANOVA. Statistical analyses were performed using SigmaPlot 14.0. Centre lines and crosses within box plots, respectively, represent sample medians and means. Box limits indicate the 25th and 75th percentiles; whiskers extend 1.5 times the interquartile range from the 25th and 75th percentiles.

### Supplementary Figures

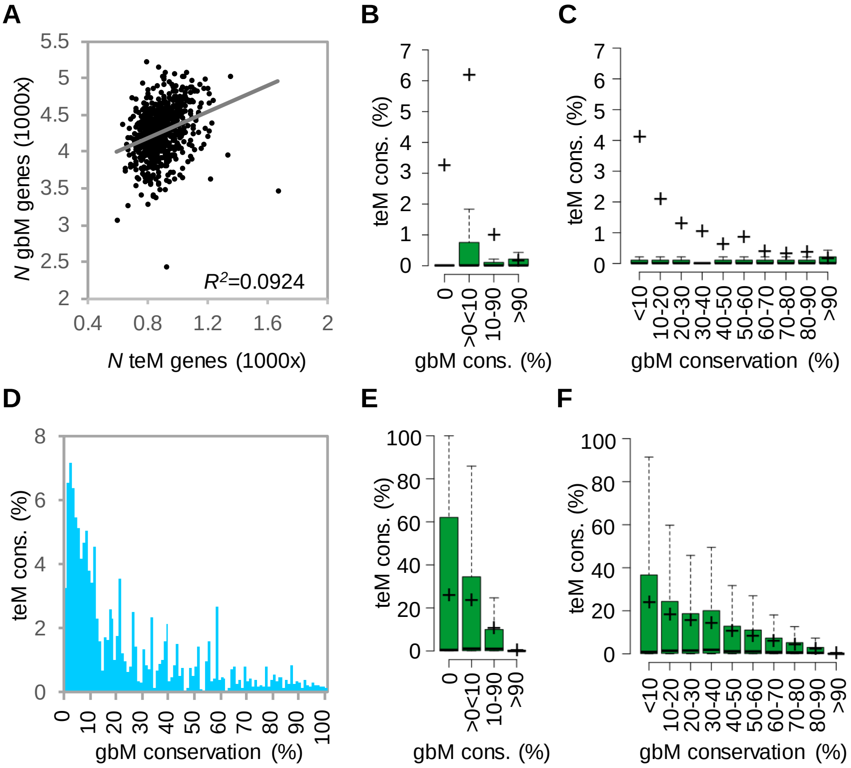


**Figigure S1. GbM and teM prevalence is independent in *Arabidopsis***. (**A**) Correlation between the number (*N*) of gbM and teM genes in 725 *Arabidopsis* accessions using published definitions of gbM and teM (46). (**B**-**D**) Conservation of teM epialleles in published [7] gbM conservation categories (**B**, four gbM classes; **C**, decile gbM bins; **D**, percentile gbM bins). In boxplots (**B** and **C**), crosses show the mean and horizontal lines the median. Analysis across three gbM conservation bins (<10%, 10-90%, and >90%) led to the published conclusion that teM frequency increases with gbM frequency[7]. However, we noted that the published <10% category contains only UM genes. Categorizing the published data[7] in various ways (**B**-**D**) shows that teM prevalence decreases with increasing gbM. These results are broadly consistent with those obtained with our gbM and teM definitions, presented in **E** (four gbM classes), **F** (decile gbM bins), and Figure 1D (percentile gbM bins).

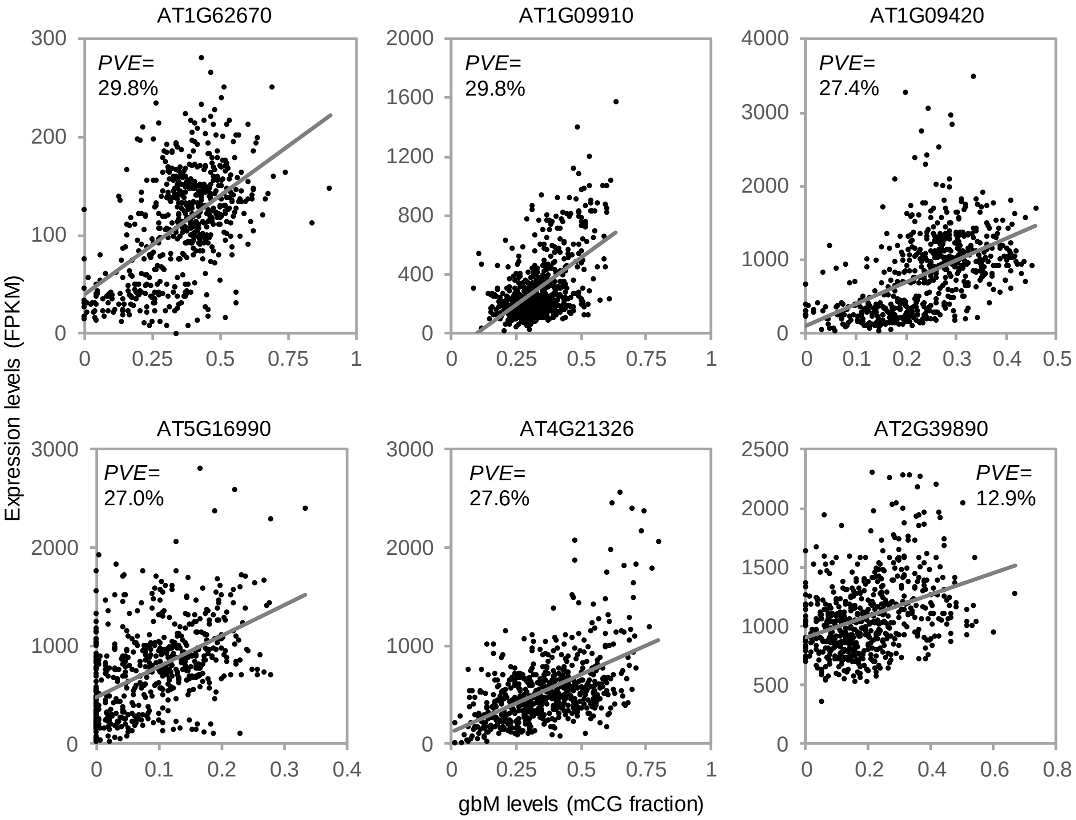


**Figure S2. Association between gbM and expression of *Arabidopsis* genes**. GbM and expression of six example genes across accessions. Percent expression variance explained (PVE) by gbM is indicated.

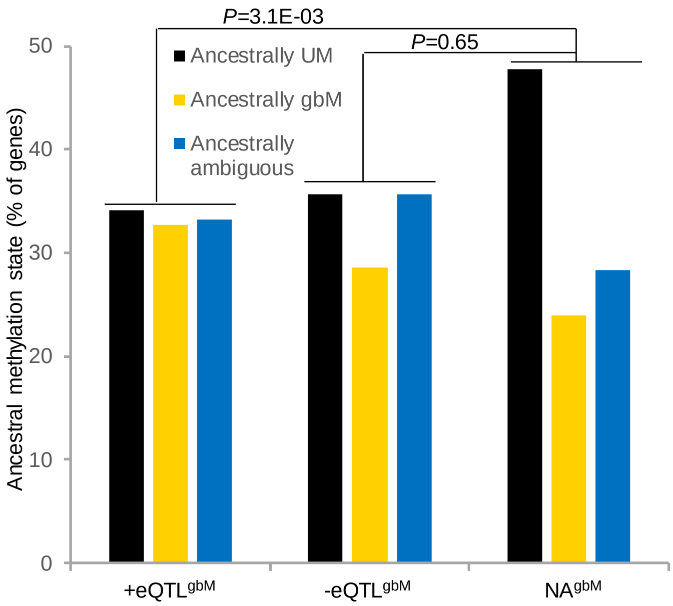


**Figure S3. +eQTL$^{gbM}$ genes are relatively enriched for ancestral gbM**. Ancestral methylation states of +eQTL$^{gbM}$, -eQTL$^{gbM}$, NA$^{gbM}$ genes. Ancestral methylation states were determined by analyzing methylation states of *Arabidopsis thaliana* orthologues in *Arabidopsis lyrata* and *Capsella rubella* and retrieved from[28]. *P* values correspond to chi-squared test.

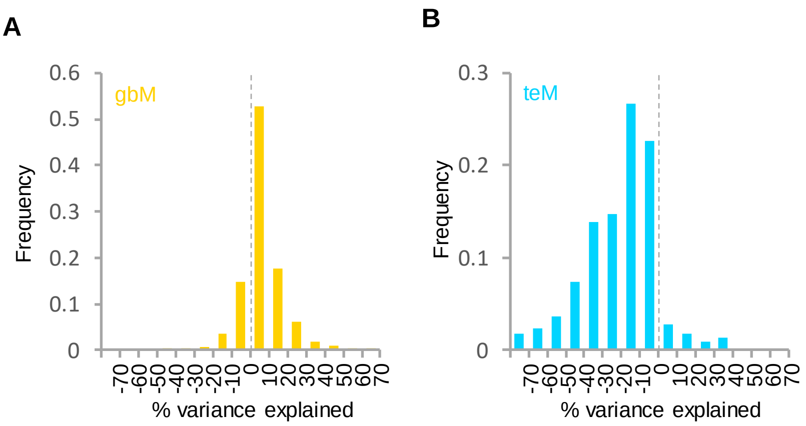


**Figure S4. Effects of gbM and teM on expression variation**. (**A** and **B**) Frequency distribution of percent expression variance of $eQTL^{gbM}$ (**A**) and $eQTL^{teM}$ (**B**) genes explained by DNA methylation.

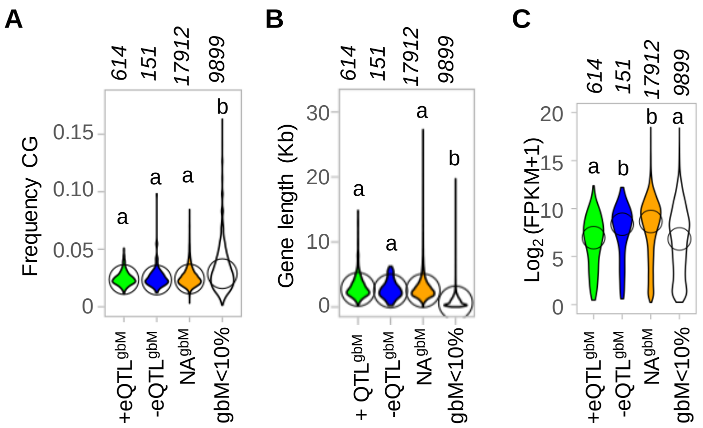


**Figure S5. +eQTL$^{gbM}$ genes are lowly expressed compared to other gbM genes**. (**A**-**C**) CG dinucleotide frequency (**A**), gene length (**B**), and gene expression (**C**) of eQTL$^{gbM}$ genes, NA$^{gbM}$ genes, and genes with gbM in <10% of accessions. Different letters signify *P*<0.001, one-way ANOVA, Dunn's test. Numbers of genes within each group are indicated.

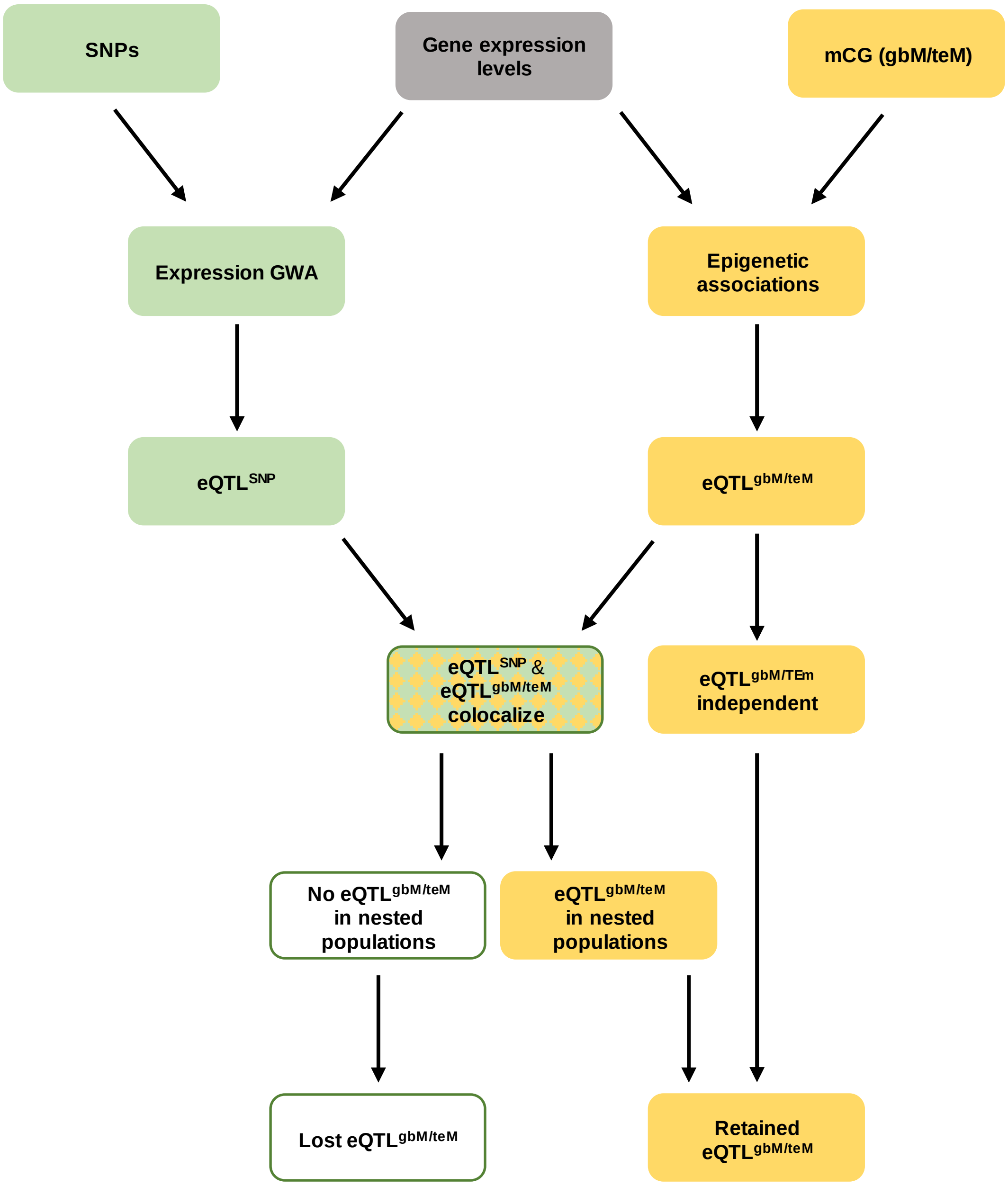


**Figure S6. Schematic of the pipeline used to account for the effects of genetic variation on eQTL$^{gbM/teM}$ gene expression**. GWA analyses were performed using SNP data to identify genetic *cis* eQTLs (eQTL$^{SNP}$) underlying expression variance of eQTL$^{gbM/teM}$ genes. If a eQTL$^{gbM/teM}$ colocalized with a eQTL$^{SNP}$, two nested populations were generated based on the GWA SNP. Association of intragenic DNA methylation with gene expression was re-examined within nested populations. eQTL$^{gbM/teM}$ were retained if they did not colocalize with eQTLs$^{SNP}$ or if a eQTL$^{gbM/teM}$ was detected in at least one nested population.

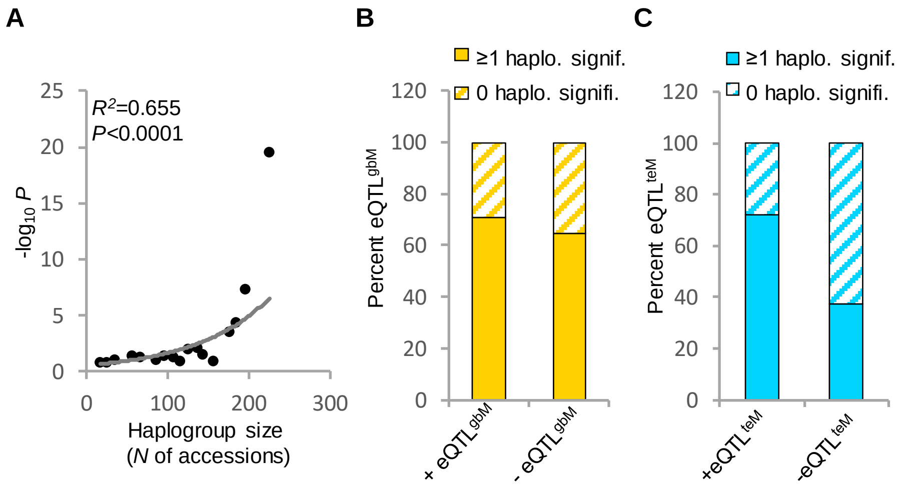


**Figure S7. Most eQTL$^{gbM/teM}$ gene expression effects are independent of local SNP variation**. (**A**) Loss of statistical power for association analyses in haplogroups due to decreasing population size. The -$\log_{10}$ *P* values for associations between gbM/teM and gene expression exhibit an exponential increase with respect to the number of accessions in haplogroups. Indicated $R^2$ and *P* values correspond to exponential regression. (**B** and **C**) Frequency of eQTL$^{gbM}$ (**B**) and eQTL$^{teM}$ (**C**) genes with a significant association between mCG and gene expression in at least one haplogroup.

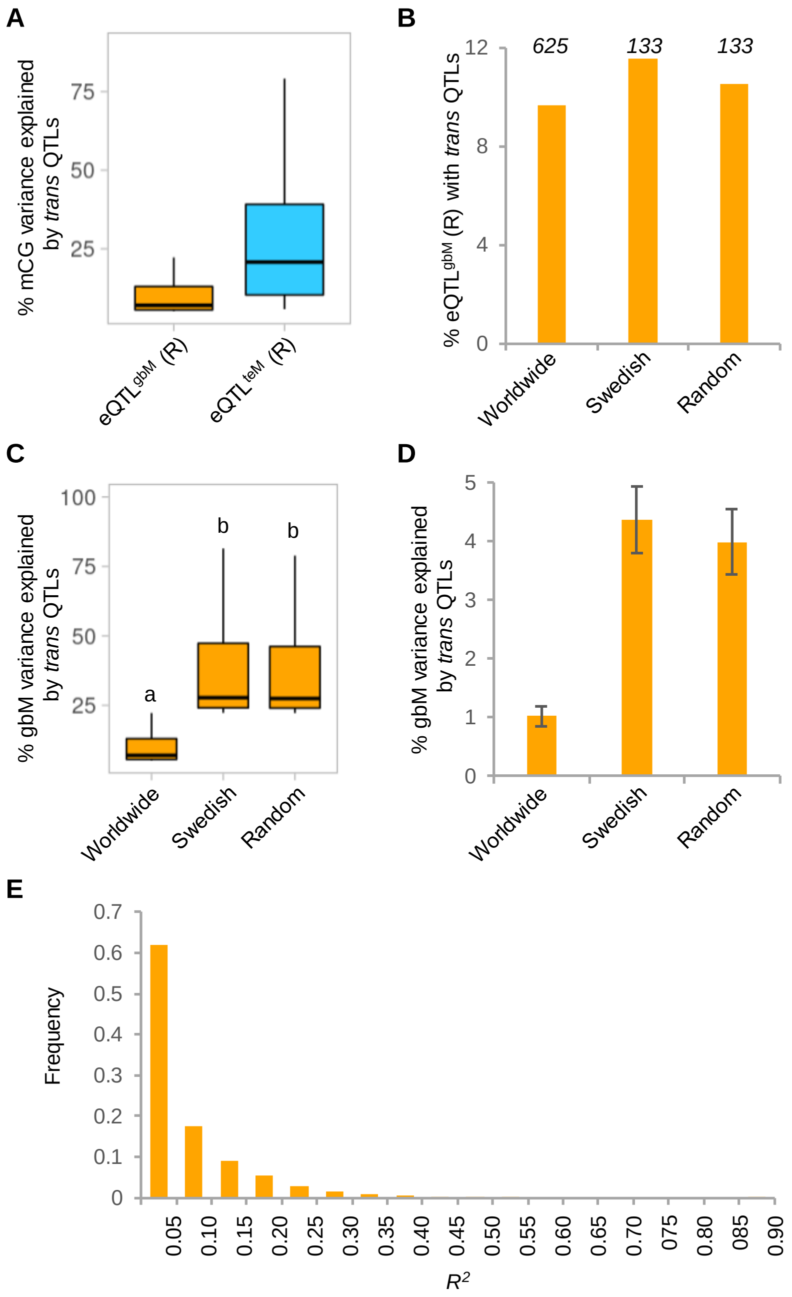


**Figure S8. Trans genetic variation explains a minor fraction of gbM variation in a worldwide population**. (**A**) Effect sizes of *trans* polymorphism on mCG variation of retained $eQTL^{gbM/teM}$ genes having *trans* QTLs. (**B**) Percentage of retained $eQTL^{gbM}$ having *trans* QTLs in the entire worldwide population, the Swedish population, and a random population of equal size to the Swedish population. Numbers of accessions in each panel are indicated. (**C**) Effect sizes of *trans* genetic polymorphisms on gbM variation of retained $eQTL^{gbM}$ genes having *trans* QTLs in worldwide, Swedish, and random populations. Different letters signify $P<0.05$, one-way ANOVA, Tukey's test. (**D**) Average effects (± standard error) of *trans* genetic variation on gbM variation of all retained $eQTL^{gbM}$ genes in worldwide, Swedish, and random populations. (**E**) Frequency distribution of correlation ($R^2$) between gbM levels of individual genes and global gbM levels of accessions. A linear model is used to estimate $R^2$.

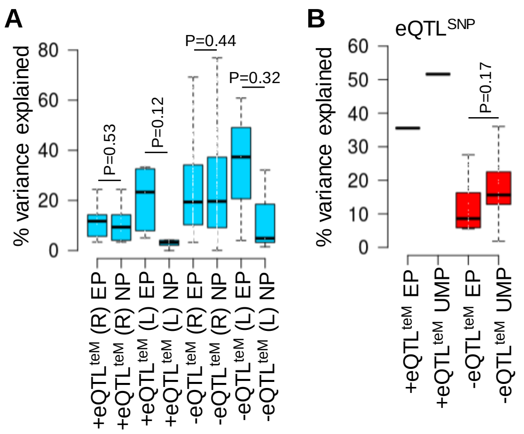


**Figure S9. TeM effects expression variance independently of SNP variation.** (**A**) Effect sizes of teM on expression of positive and negative retained (R) or lost (L) eQTL$^{teM}$ genes in the entire population (EP) and after accounting for expression GWA SNPs in nested populations (NP). (**B**) Effects of SNPs on expression variation of retained eQTL$^{teM}$ genes in the entire (EP) and unmethylated populations (UMP).

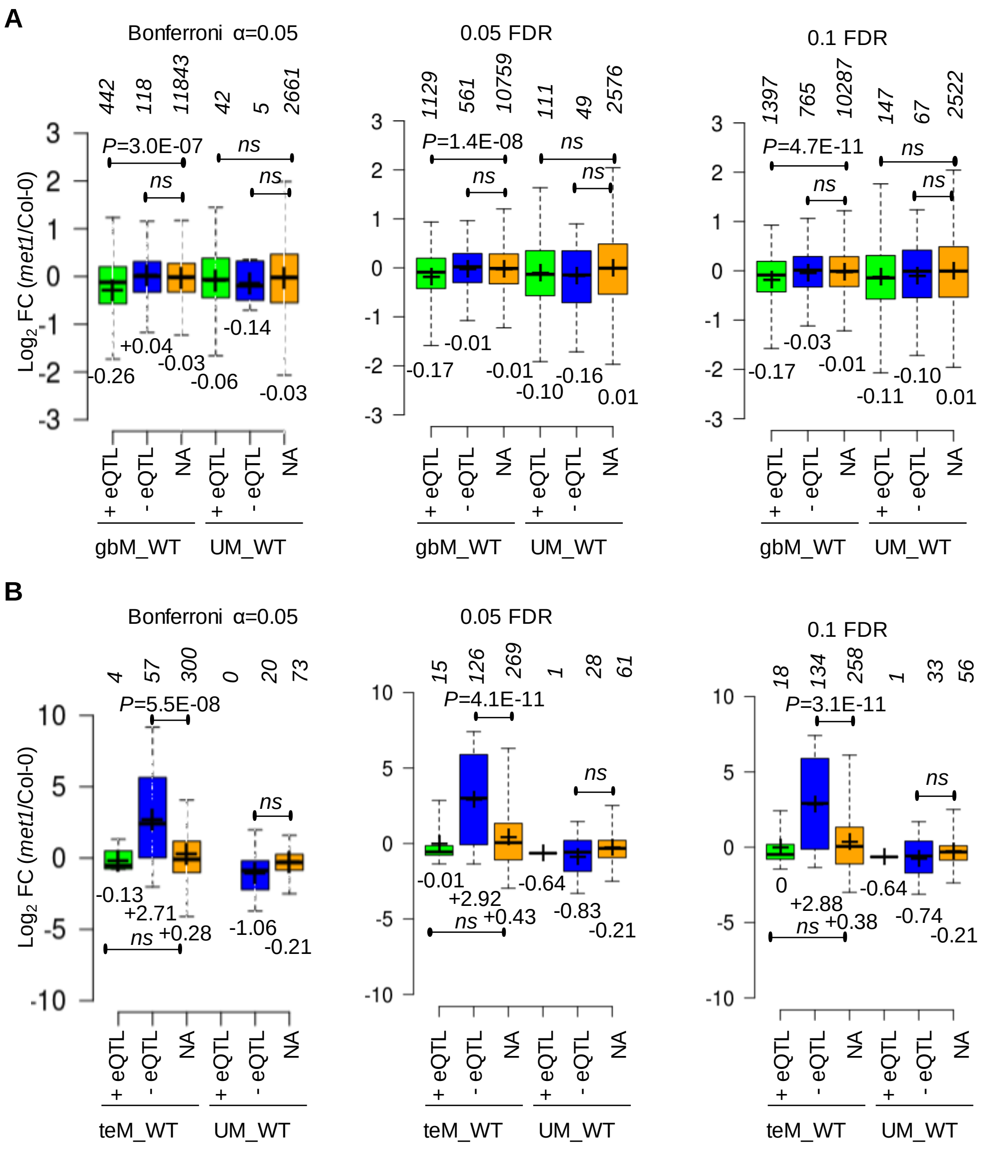


**Figure S10. GbM activates and teM represses gene expression**. (**A** and **B**) Expression in *met1* seedlings compared to Col-0 of Bonferroni (α=0.05), 0.05 FDR, and 0.1 FDR eQTL$^{gbM}$ (**A**) and eQTL$^{teM}$ (**B**) genes. Genes with intragenic DNA methylation (gbM; teM) or UM in Col-0 were analyzed separately. Numbers of genes within each group are indicated. *P*, Wilcoxon rank sum test.

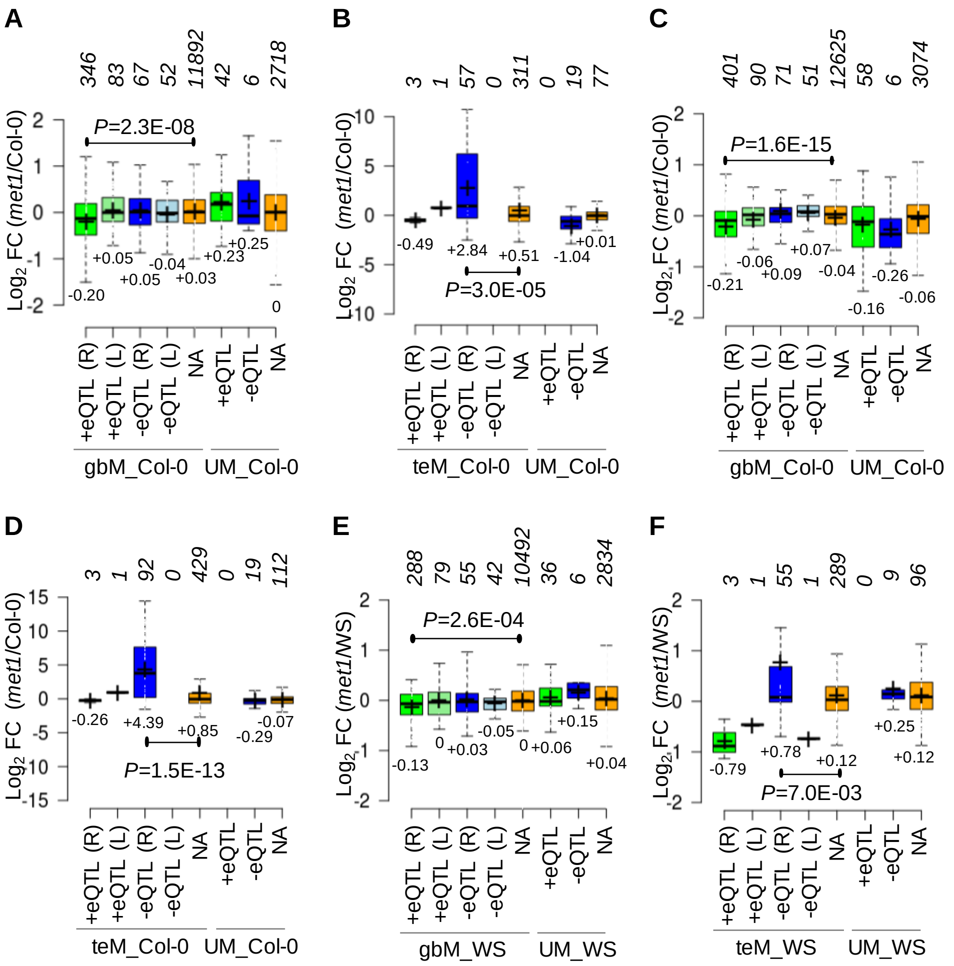


**Figure S11. GbM promotes gene expression.** (**A** and **B**) Expression in *met1* leaves compared to Col-0 of Bonferroni (α=0.05) $eQTL^{gbM}$ (**A**) and $eQTL^{teM}$ (**B**) genes retained (R) or lost (L) after accounting for genetic variation. Genes methylated (gbM_Col-0; teM_Col-0) and unmethylated (UM_Col-0) in Col-0 were analyzed separately. Numbers of genes within each group are indicated. *P*, Wilcoxon rank sum test. (**C** and **D**) Expression in *met1* inflorescence compared to Col-0 of Bonferroni (α=0.05) $eQTL^{gbM}$ (**A**) and $eQTL^{teM}$ (**B**) genes retained (R) or lost (L) after accounting for genetic variation. Genes methylated (gbM_Col-0; teM_Col-0) and unmethylated (UM_Col-0) in Col-0 were analyzed separately. (**E** and **F**) Expression in *met1* compared to WS of Bonferroni (α=0.05) $eQTL^{gbM}$ (**E**) and $eQTL^{teM}$ (**F**) genes retained (R) or lost (L) after accounting for genetic variation. Genes methylated (gbM_WS; teM_WS) and unmethylated (UM_WS) in WS were analyzed separately. Numbers of genes within each group are indicated. *P*, Wilcoxon rank sum test.

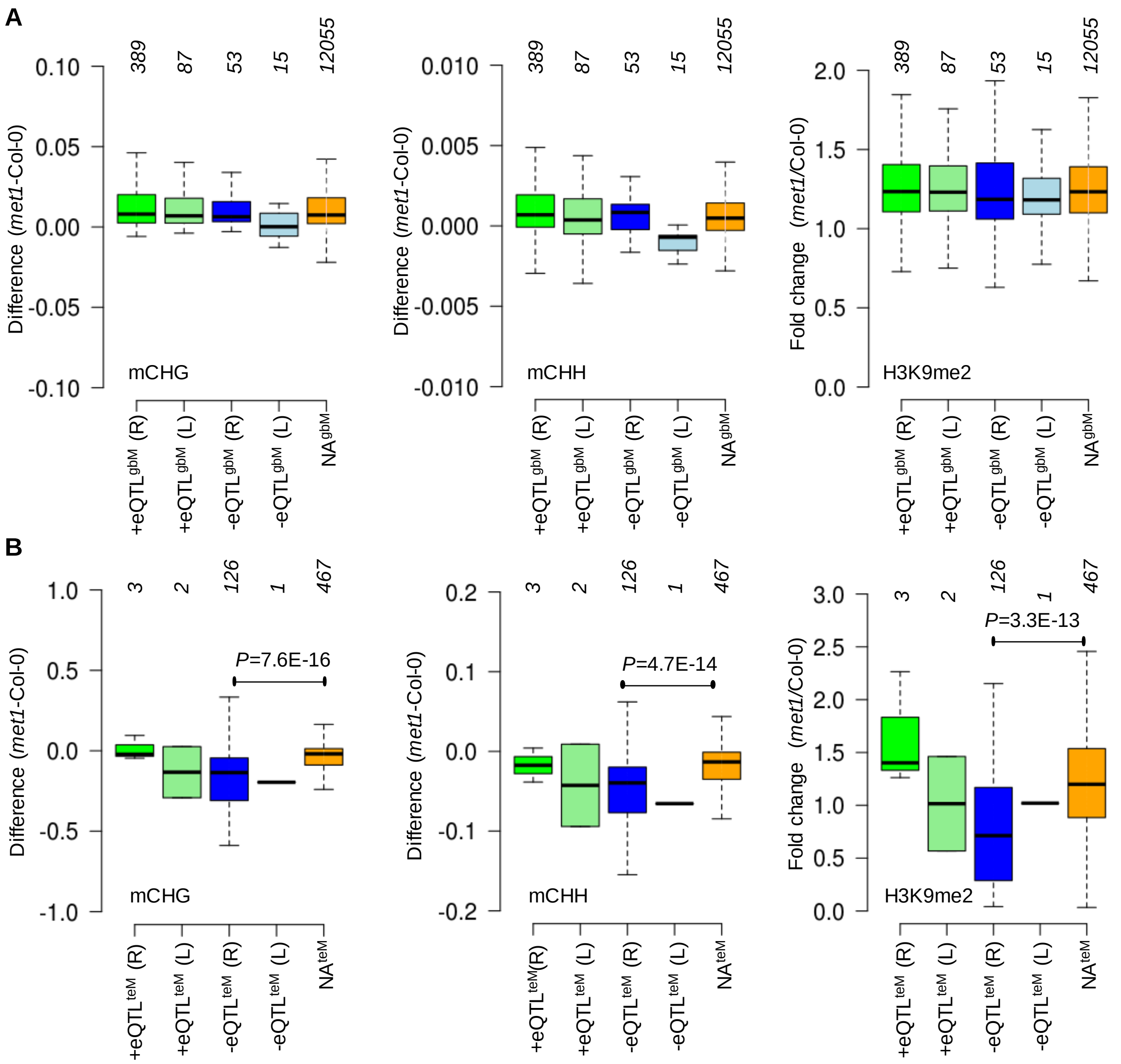


**Figure S12. Changes in non-CG methylation and H3K9me2 in *met1*.** (**A** and **B**) CHG and CHH methylation and H3K9me2 (dimethylation of lysine 9 of histone H3) in *met1* compared to Col-0 of Bonferroni ($\alpha$=0.05) $eQTL^{gbM}$ (**A**) and $eQTL^{teM}$ (**B**) genes. Published data [6] for non-CG methylation and H3K9me2 were analyzed for genes with intragenic DNA methylation (gbM; teM) in Col-0. Numbers of genes within each group are indicated. *P*, two-tailed Student's t-test, the remaining comparisons with NA genes are not significant.

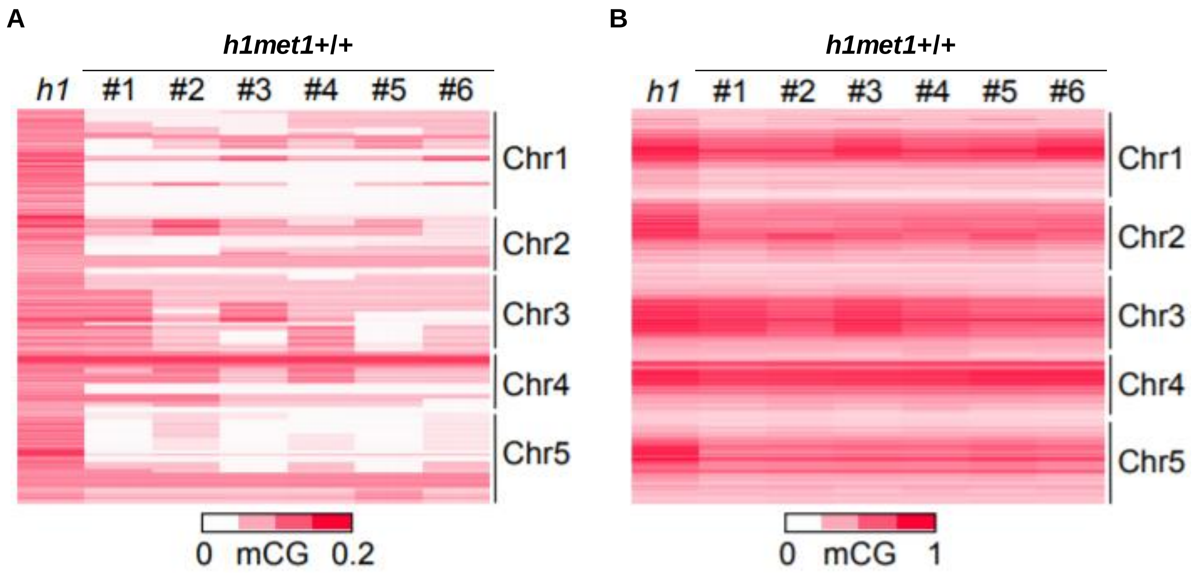


**Figure S13. DNA methylation in *h1* and *h1met1*+/+ plants.** (**A** and **B**) Heat maps showing CG methylation in gene bodies (**A**) and TEs (**B**) in *h1* or *h1met1*+/+ leaves. Six independent *h1met1*+/+ plants were isolated from segregating *h1met1*+/- plants.

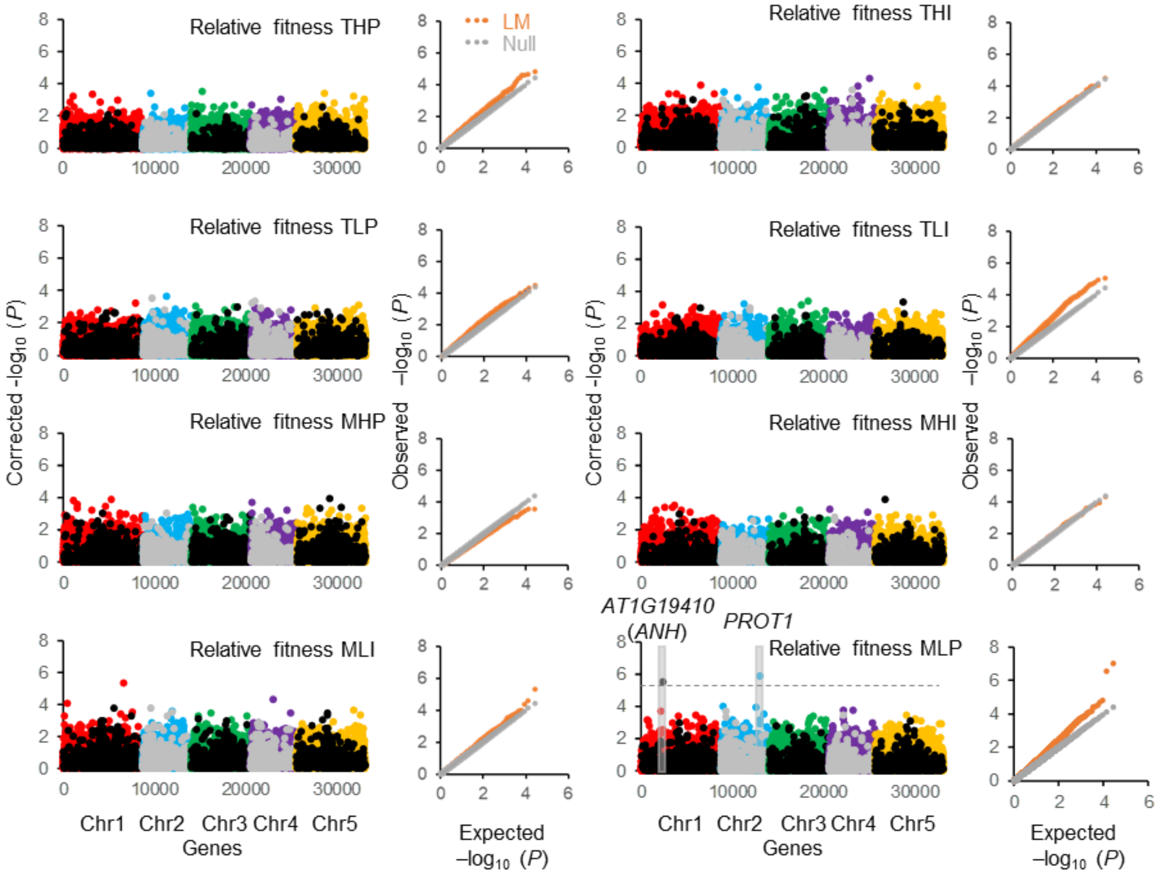


**Figure S14. Associations between intragenic methylation and relative fitness**. Manhattan and QQ plots for relative fitness epiGWA in THP, THI, TLP, THI, MHP, MHI, MLP, and MLI conditions (T, Tübingen; M, Madrid; H, high rainfall; L, low rainfall; P, high plant density; and I, individual plants). QQ plots compare the distribution of observed (orange dots) and expected (diagonal grey dots) -$\log_{10}$ *P* values. To account for confounding effects of population stratification, genomic control factor λ was used to correct association statistics. Corrected -$\log_{10}$ *P* values for gbM (colored dots) and teM (grey and black dots) markers are plotted in Manhattan plots. Horizontal dashed line shows 0.05 FDR. Manhattan plot of epiGWA mapping for relative fitness in MLP is also presented in Figure 3A.

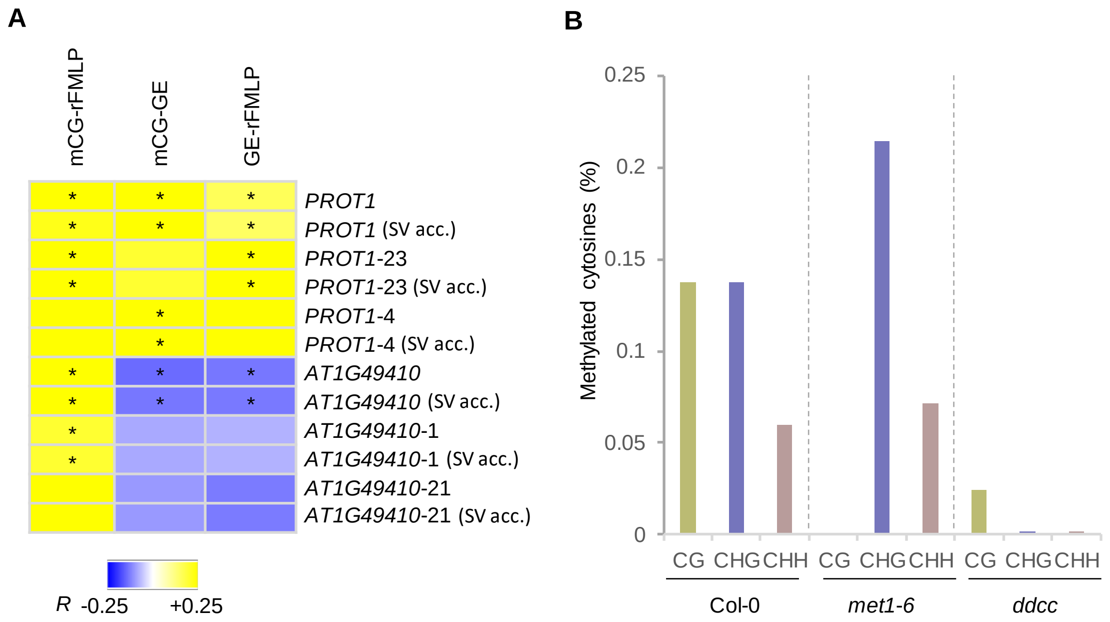


**Figure S15. Methylation and expression of epigenetic relative fitness QTLs.** (**A**) Tripartite association between intragenic DNA methylation (mCG), gene expression (GE), and relative MLP fitness (rFMLP) in the entire population and after accounting (acc.) for structural variation (SV). Associations between the three variables are shown in two independent haplogroups of *PROT1* (*PROT1-23* and *PROT1-4*) and AT1G19410 (AT1G19410-1 and AT1G19410-21) before and after accounting for SV. **P*<0.05, Pearson's correlation test. (**B**) DRM and CMT methyltransferases control teM of AT1G19410. Fractional methylation in CG, CHG, and CHH sequence contexts is shown in indicated genotypes.

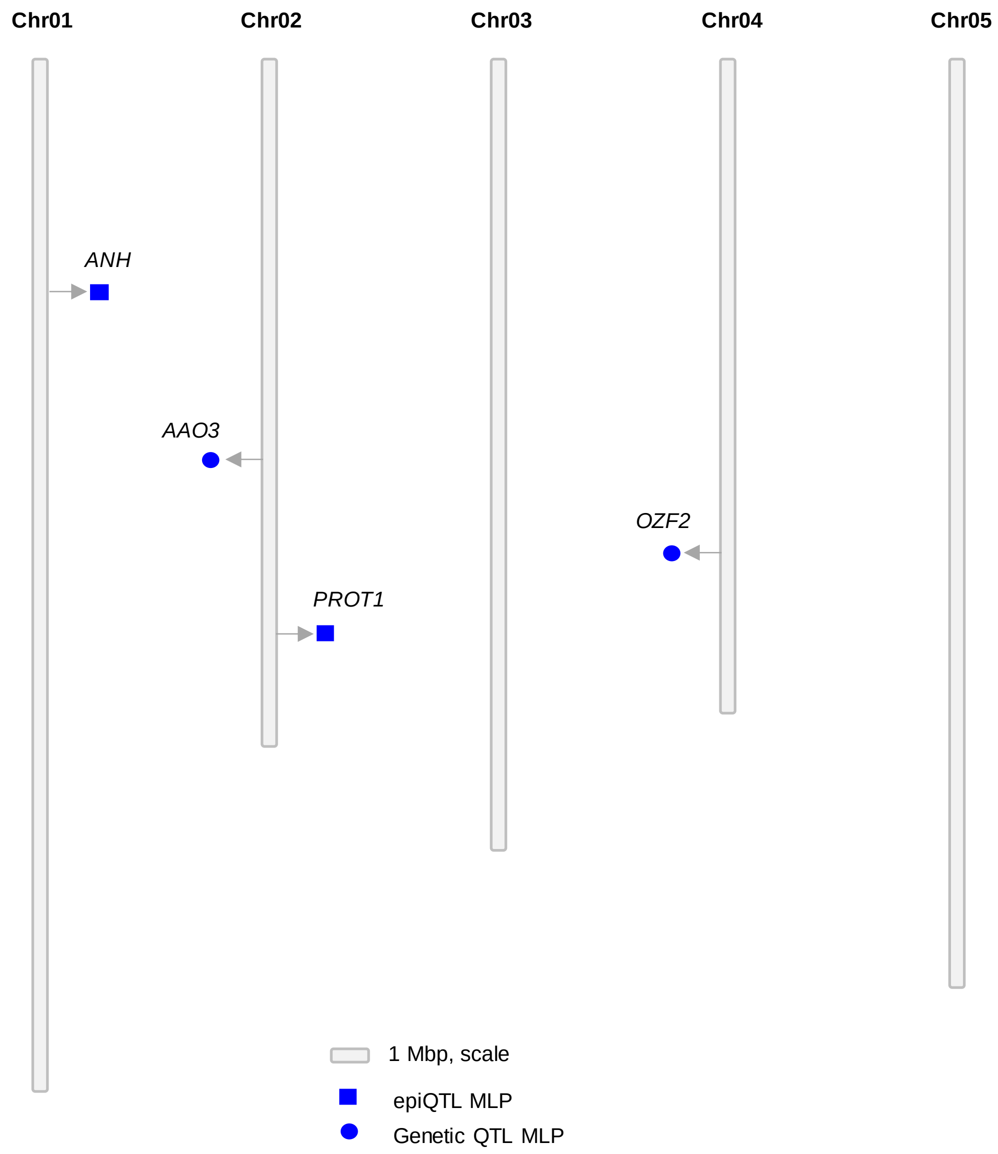


**Figure S16. Distinct epigenetic and genetic QTLs are associated with relative fitness under MLP conditions**. *Arabidopsis* physical map showing the positions of epigenetic and genetic QTLs associated with relative fitness under MLP conditions. Five chromosomes of *Arabidopsis* are presented in Mb scale. Positions of epigenetic QTLs are shown with squares to the right of chromosomes, and circles on the left depict positions of genetic QTLs.

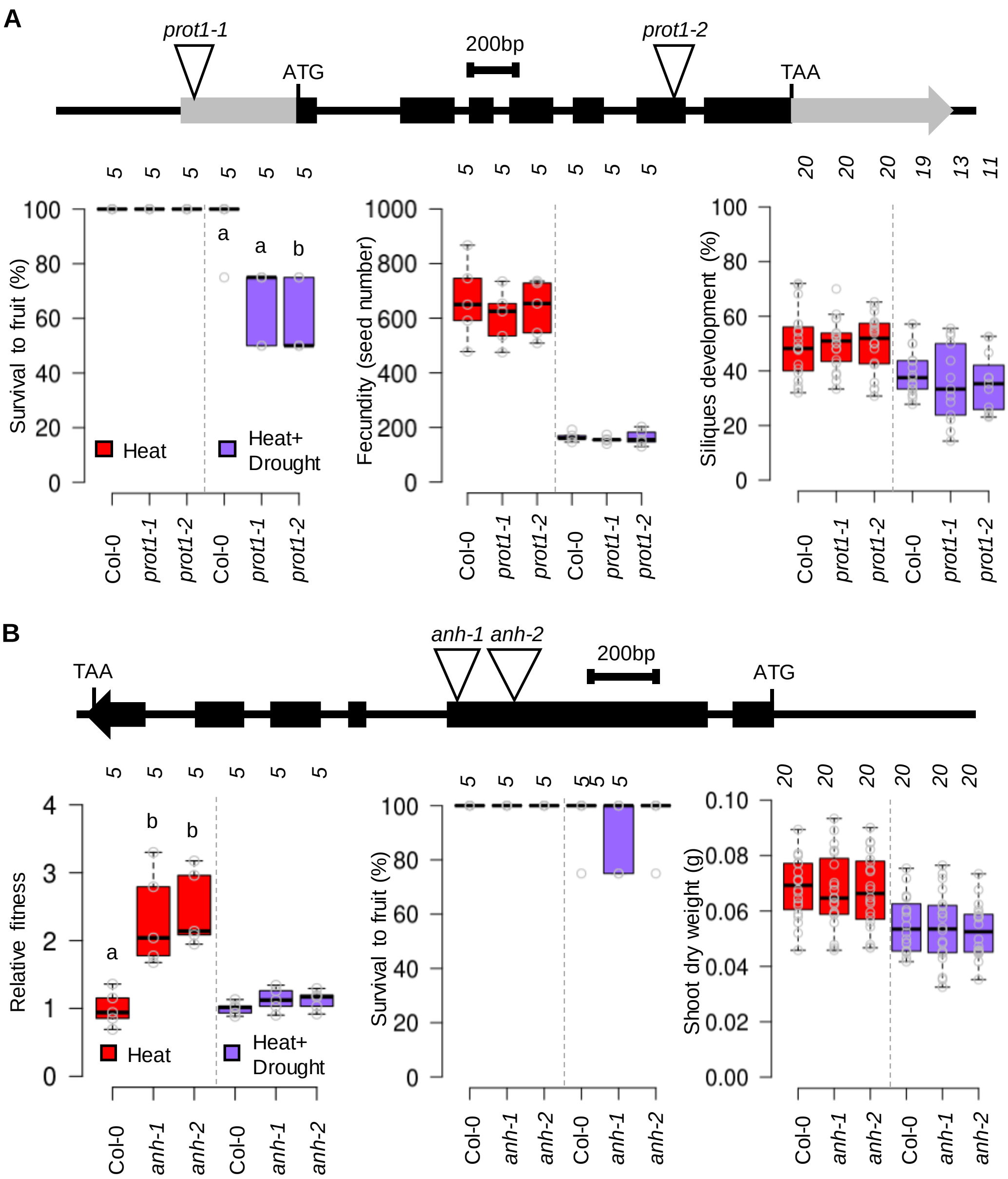


**Figure S17. *PROT1* and AT1G19410 (*ANH*) affect fitness under heat and drought stress**. (**A** and **B**) The upper panels display schematic representations of *PROT1* (**A**) and *ANH* (**B**) genomic regions with positions of the T-DNA insertions. Boxplots (**A**) show survival to fruit, fecundity, and fertility (% of flowers developing siliques) phenotypes of *prot1* mutant and Col-0 wild type plants under heat or combined heat and drought stress. Boxplots (**B**) show relative fitness, survival to fruit, and shoot dry weight of *anh* relative to Col-0. Numbers of independent experiments are indicated for survival to fruit and fecundity (**A**), and for relative fitness and survival to fruit (**B**). Numbers of plants are indicated for fertility (**A**) and shoot weight (**B**). Different letters signify *P*<0.05, one-way ANOVA, Tukey's test.

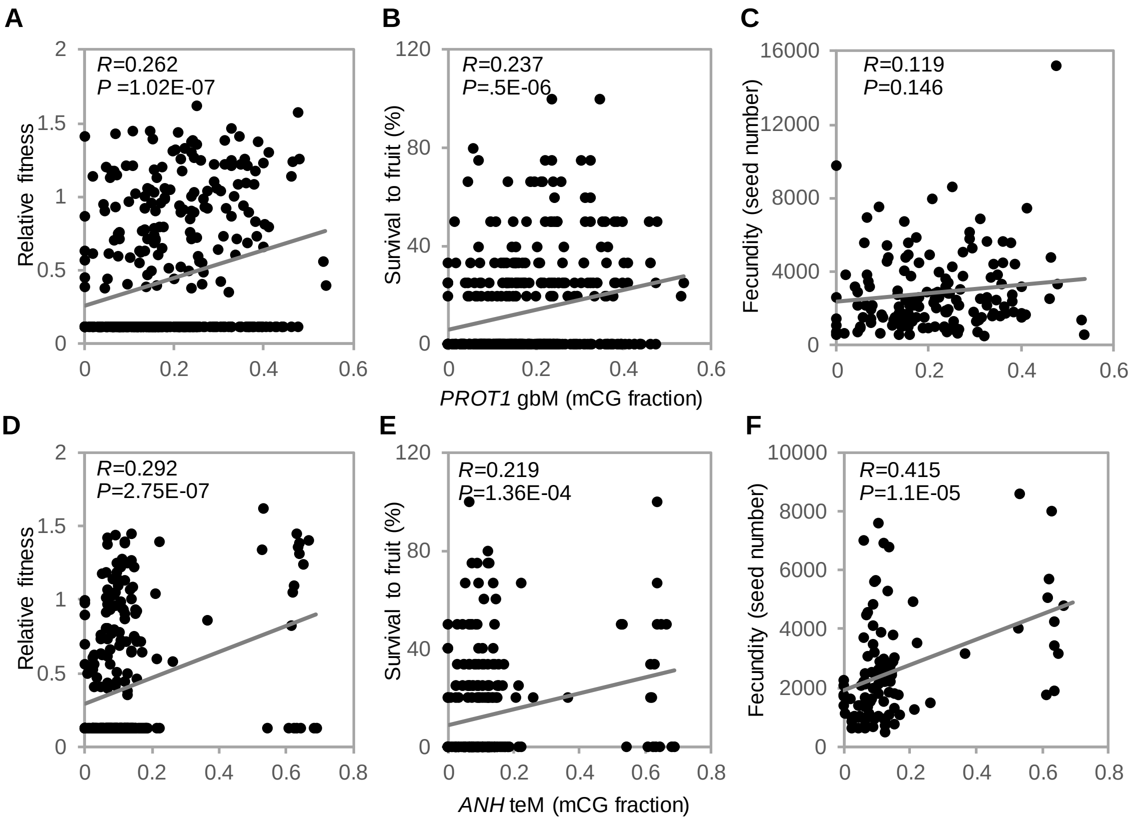


**Figure S18. *PROT1* gbM and *ANH* teM are respectively associated with survival and fecundity under MLP conditions**. (**A**-**C**) Association of *PROT1* gbM with relative fitness (**A**), survival to fruit (**B**), and fecundity (**C**). (**D**-**F**) Association of *ANH* teM with relative fitness (**D**), survival to fruit (**E**), and fecundity (**F**). Correlation coefficients (*R*) and *P* values of associations are indicated.

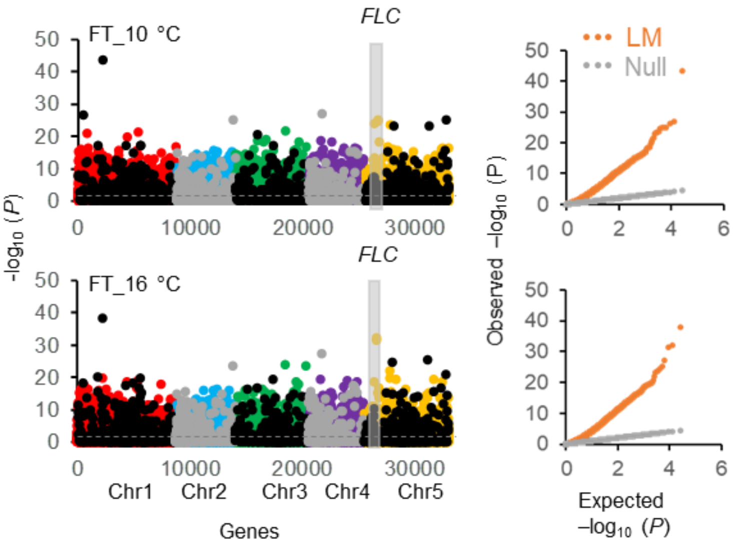


**Figure S19. Population structure confounds association statistics of linear model epiGWA mapping for flowering phenotypes**. Manhattan and QQ plots of epiGWA mapping for flowering phenotypes (FT_10°C and FT_16°C). Associations between intragenic DNA methylation (mCG) levels and flowering time were examined using a linear model. QQ plots compare the distribution of observed (orange dots) and expected (diagonal grey dots) $-\log_{10}$ *P* values. Colored dots in Manhattan plots represent gbM markers and grey and black dots correspond to teM markers. Horizontal dashed line shows 0.05 FDR. mCG levels of around 7,500 genes are significantly associated with flowering phenotypes. We expect some of these associations to be real, for example the well-known flowering regulator, *FLC*. However, it is difficult to distinguish true associations from spurious ones. Strong inflation of observed *P* values compared to expectation also results in high values for genomic control factor (λ) (Table S10). Genomic control method in cases of high λ can be anticonservative, thus is not recommended to account for population stratification[29]. Linear model GWA mapping with genomic control is generally considered inappropriate for structured phenotypes[12] and is not used in this study.

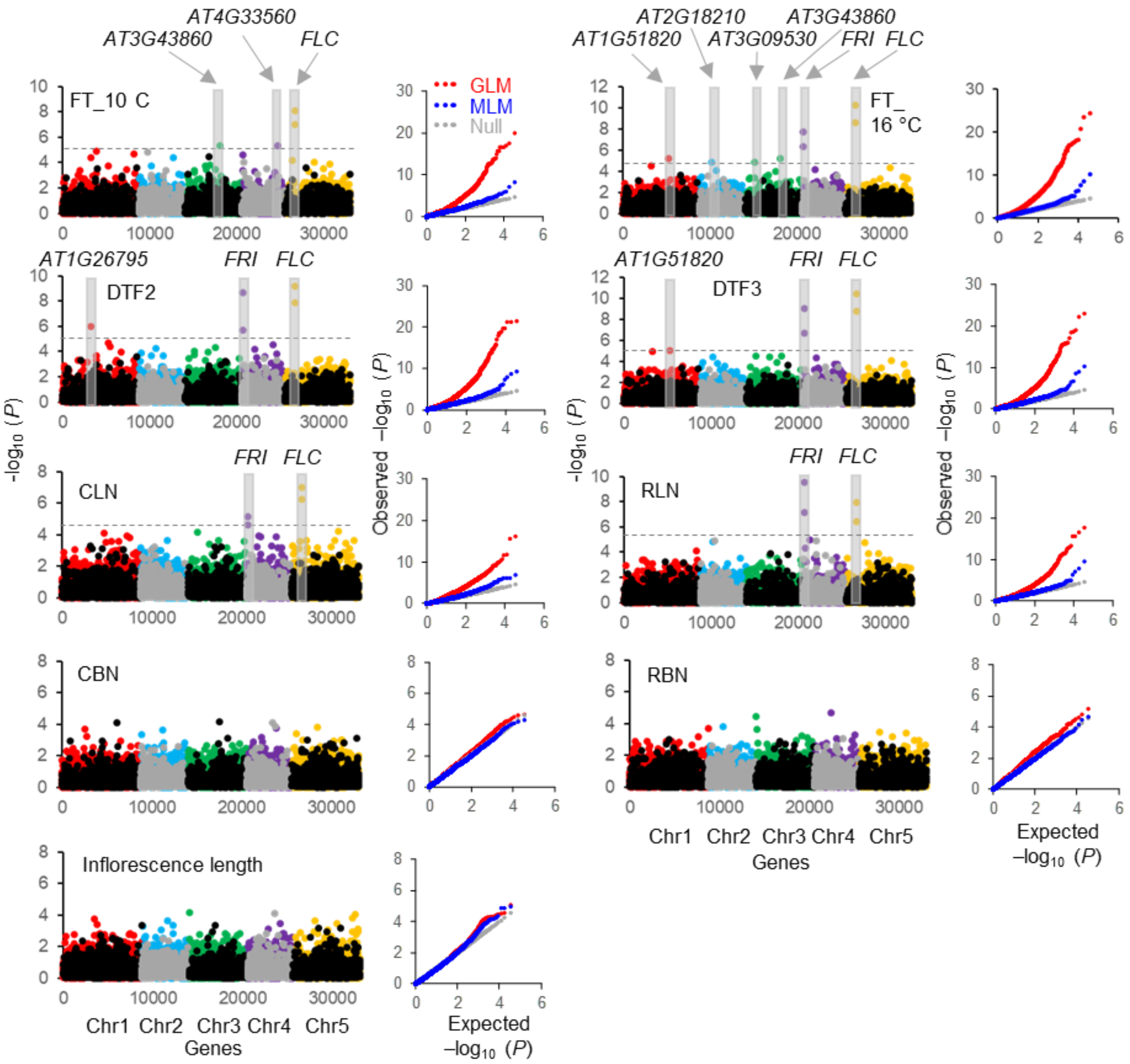


**Figure S20. Mixed linear model corrects population structure in epiGWA analyses for flowering phenotypes**. EpiGWA mapping for nine flowering related phenotypes (flowering time at 10°C (FT_10°C), flowering time at 16°C (FT_16 °C), number of days for inflorescence stalk to reach 1 cm (DTF2), number of days to opening of the first flower (DTF3), number of cauline leaves (CLN), number of rosette leaves (RLN), cauline branch number (CBN), primary number of inflorescence branches (RBN), and length of primary inflorescence stalk. Associations between epiallelic states (UM and gbM; UM and teM) of genes and each flowering phenotype were examined using a generalized linear model (GLM) or a mixed linear model (MLM). QQ plots compare the distribution of observed GLM epiGWA (red dots), observed MLM epiGWA (blue dots) and expected (diagonal grey dots) -$\log_{10}$ *P* values. In Manhattan plots, colored dots depict gbM markers and teM markers are shown using grey and black dots. Horizontal dashed lines show MLM 0.05 FDR. Epigenetic flowering QTLs (passing 0.05 FDR) are highlighted with grey bars.

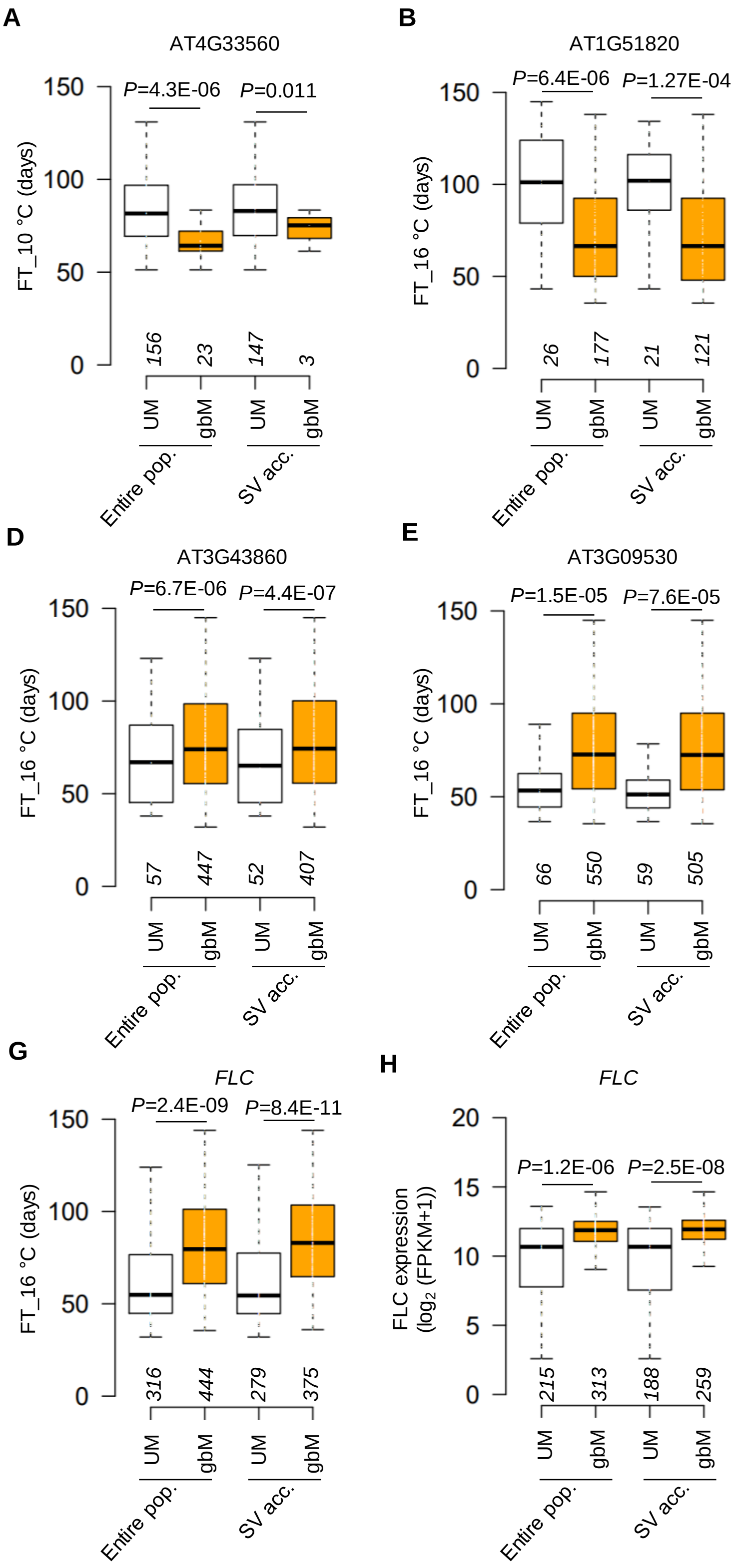


**Figure S21. Most fQTL$^{gbM}$ influence flowering time diversity independently of structural variation.** (**A-G**) Flowering time of UM and gbM accessions for *AT4G33560* (**A**), *AT1G51820* (**B**), *AT2G18210* (**C**), *AT3G43860* (**D**), *AT3G09530* (**E**), *FRI* (**F**), and *FLC* (**G**) in the entire population and after accounting for structural variation (SV). *P* values correspond to MLM epiGWA. Association between *AT4G33560* gbM and FT_10°C is marginally significant. Therefore, *AT4G33560* fQTL$^{gbM}$ is largely confounded by SV. (**H**) *FLC* expression in UM and gbM accessions in the entire population and after accounting for SV. *P* values correspond to MLM epiGWA.

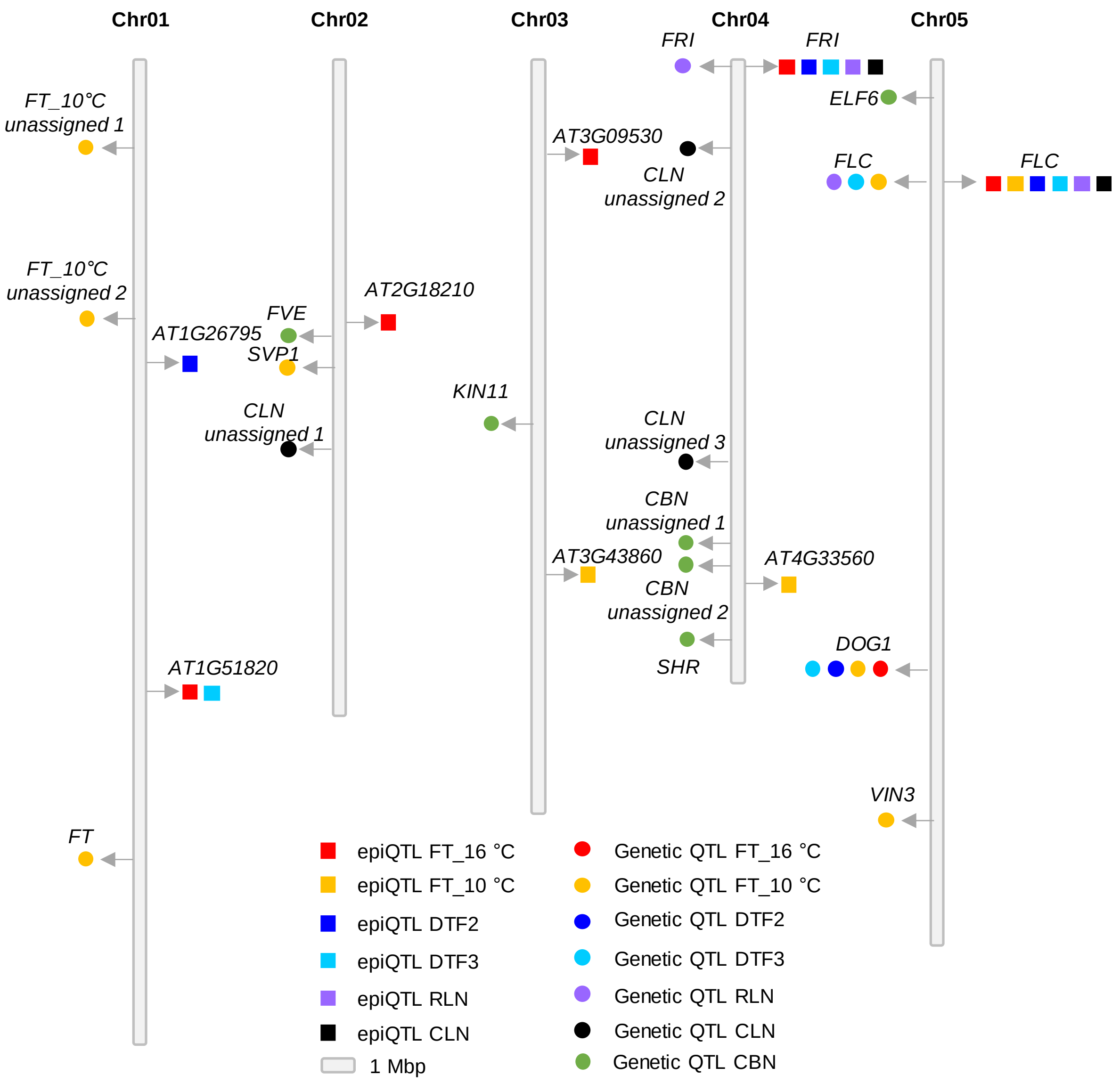


**Figure S22. Epigenetic and genetic QTLs associated with flowering phenotypes are largely independent**. *Arabidopsis* physical map showing the positions of epigenetic and genetic QTLs underlying flowering phenotypes. Five chromosomes are shown in Mb scale. Positions of epigenetic QTLs are shown with squares to the right of chromosomes, and circles on the left depict positions of genetic QTLs.

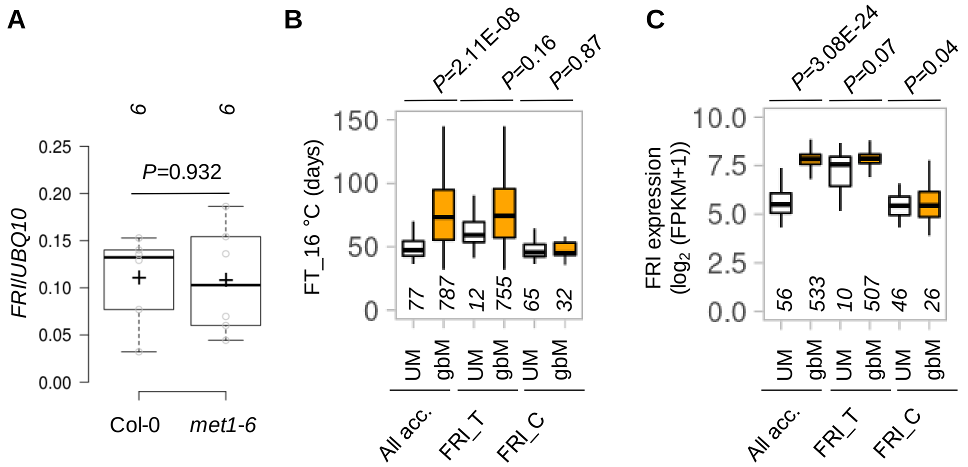


**Figure S23. *FRI* expression $QTL^{gbM}$ and flowering $QTL^{gbM}$ are confounded by genetic variation**. (**A**) Transcript levels of *FRI* relative to *UBQ10* in Col-0 and *met1*, assessed by qRT-PCR in six biological replicates. *P*, two-tailed Student's t test. (**B** and **C**) Association of epiallelic states of *FRI* with FT_16°C (**B**) and *FRI* expression (**C**) in all accessions and in nested populations (FRI_T and FRI_C) fixed for GWA *FRI* SNPs. Numbers of accessions for each group are indicated. *P* values correspond to MLM epiGWA.

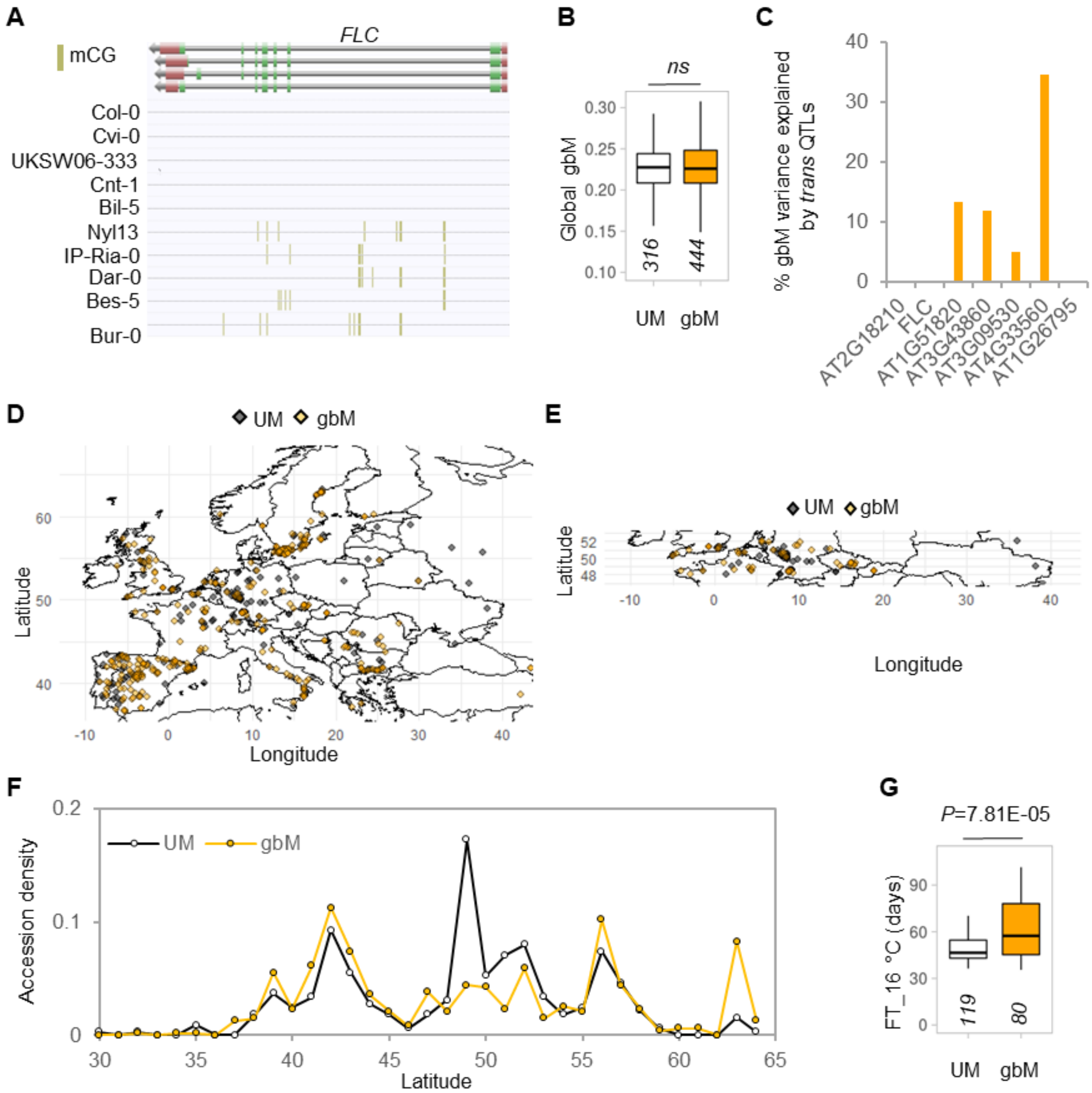


**Figure S24. Distribution of *FLC* epialleles**. (**A**) AnnoJ browser snapshot for *FLC* UM and gbM accessions. (**B**) Effects of *trans* genetic variation on gbM variation of *FLC* and other epigenetic fQTLs. (**C**) Geographic distribution of accessions having UM and gbM *FLC* epialleles. (**D**) Distribution of *FLC* UM and gbM epiallele-harboring accessions within 48-52° latitude. (**E**) Distribution of *FLC* UM and gbM epiallele-harboring accessions within 48-52° latitude. (**F**) Density of FLC UM and gbM accessions across latitude. (**G**) Association of FLC epiallelic states with FT_16°C within 48-52° latitude. Accession numbers are indicated. P value corresponds to MLM epiGWA.

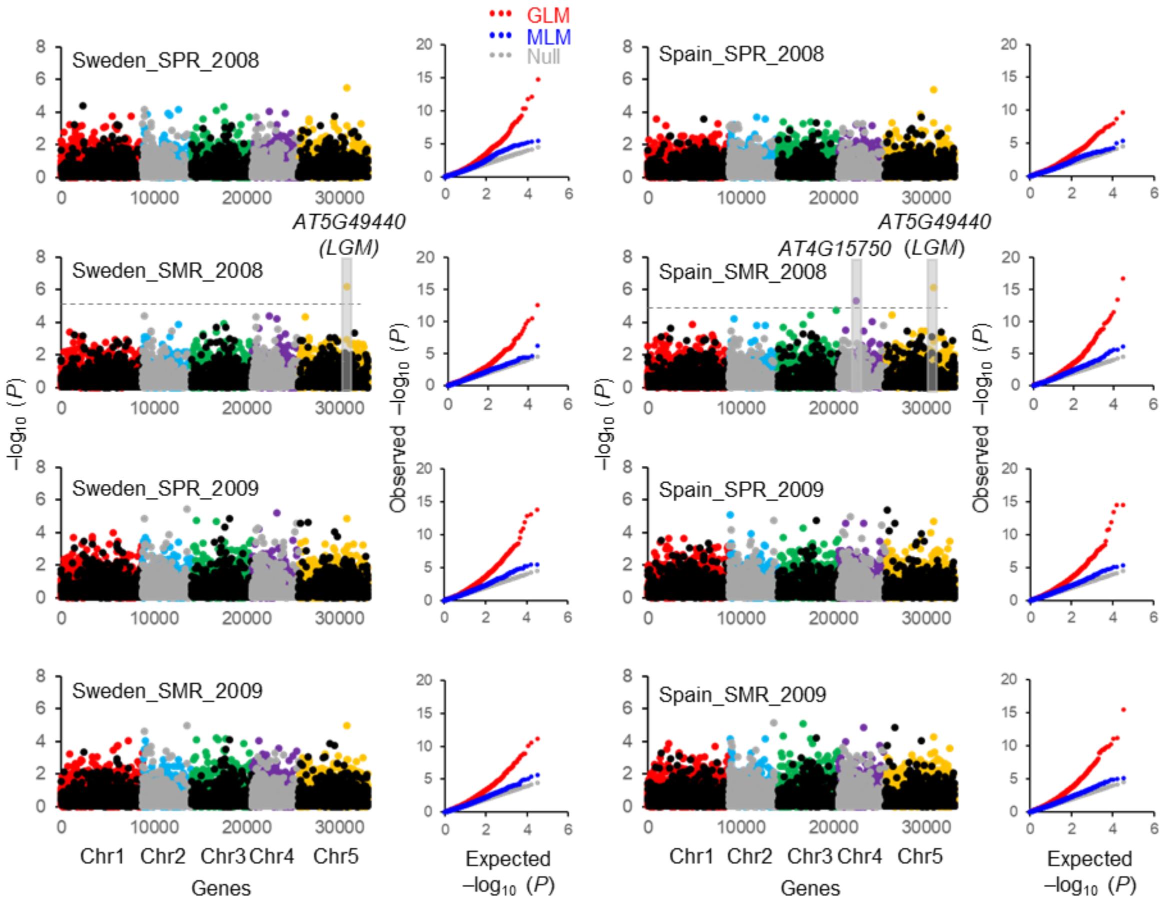


**Figure S25. GbM variation influences flowering time in simulated climates**. EpiGWA mapping for flowering time under simulated local climates in Spain and Sweden across two seasonal plantings: spring (SPR) and summer (SMR) in 2008 and 2009. EpiGWA mapping was performed using a generalized linear model (GLM) and a mixed linear model (MLM). QQ plots compare the distribution of observed GLM epiGWA (red dots), observed MLM epiGWA (blue dots) and expected (diagonal grey dots) -$\log_{10}$ *P* values. In Manhattan plot, gbM markers are depicted with colored dots and teM markers are shown using grey and black dots. Horizontal dashed lines show MLM 0.05 FDR. Epigenetic flowering QTLs (passing 0.05 FDR) are highlighted with grey bars.

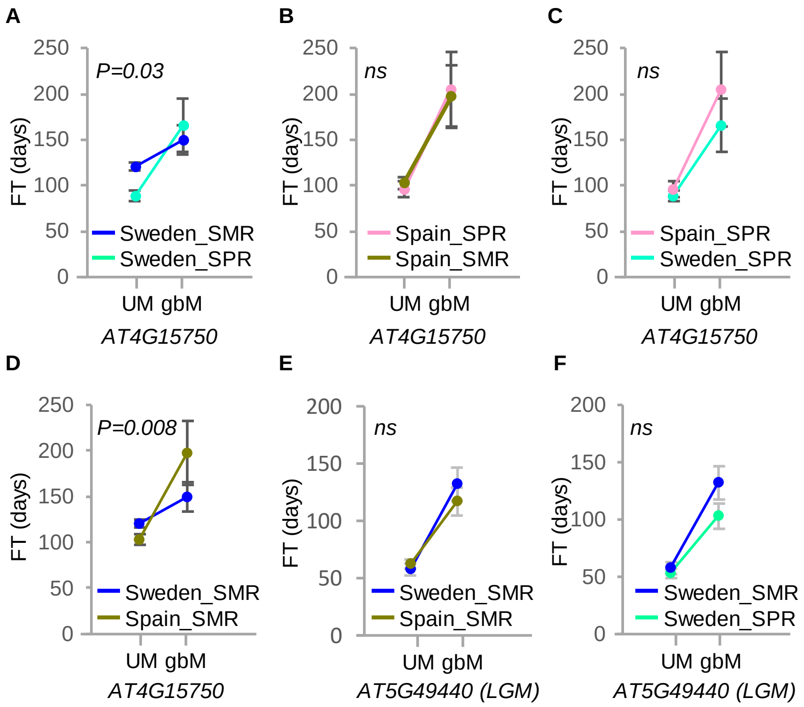


**Figure S26. GbM variation in AT4G15750 and AT5G49440 (*LGM*) affects sensitivity of flowering time to seasons and climates.** (**A** and **B**) Plots of average ± standard error (SE) flowering time (FT) of AT4G15750 UM and gbM accessions during summer (SMR) and spring (SPR) in Sweden (**A**) or Spain (**B**). (**C** and **D**) Plots of average (±SE) FT of AT4G15750 UM and gbM accessions grown in Spain and Sweden in spring (**C**) or summer (**D**). (**E** and **F**) Plots of average FT (±SE) of AT5G49440 *(LGM)* UM and gbM accessions across climates in summer (**E**) and seasons in Sweden (**F**). Sensitivity of epigenetic fQTLs to seasons or climates was assessed via two-way ANOVA.

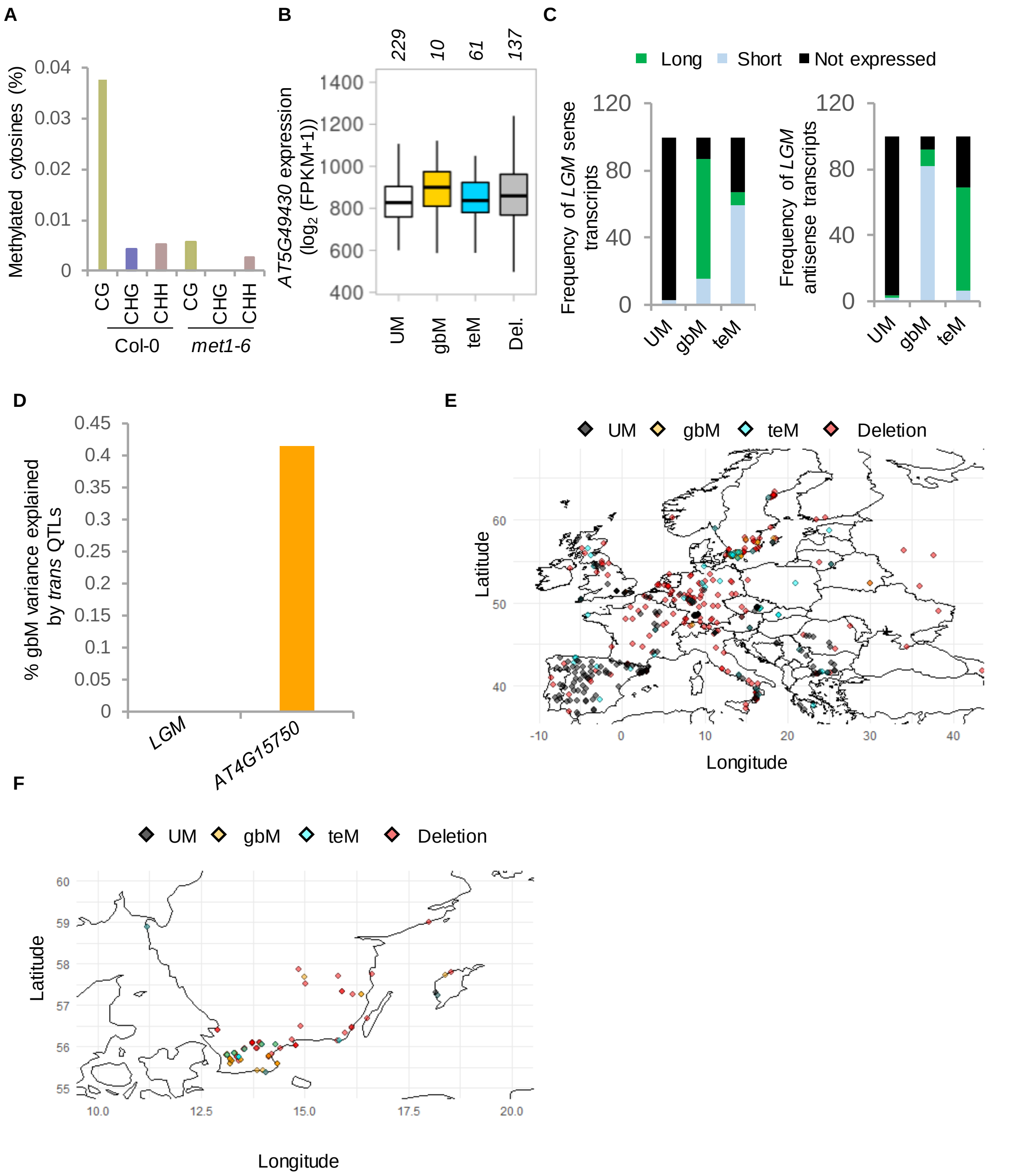


**Figure S27. Methylation, expression, and geographical distribution of *LGM* alleles**. (**A**) Fractional methylation at *LGM* in CG, CHG, and CHH sequence contexts is shown in Col-0 and *met1*. (**B**) AT5G49430 expression with *LGM* UM, gbM, teM, and deletion (epi)alleles in the indicated numbers of accessions. None of the pairwise comparisons are significant according to MLM epiGWA. (**C**) Frequency of alternative *LGM* transcripts in UM, gbM, and teM accessions. (**D**) Effects of *trans* genetic variation on gbM variation of *LGM* and *AT4G15750* epigenetic fQTLs. (**E**) Geographic distribution of *LGM* UM, gbM, teM, and deletion accessions. (**F**) Distribution of *LGM* UM, gbM, teM, and deletion accessions in southern Sweden.

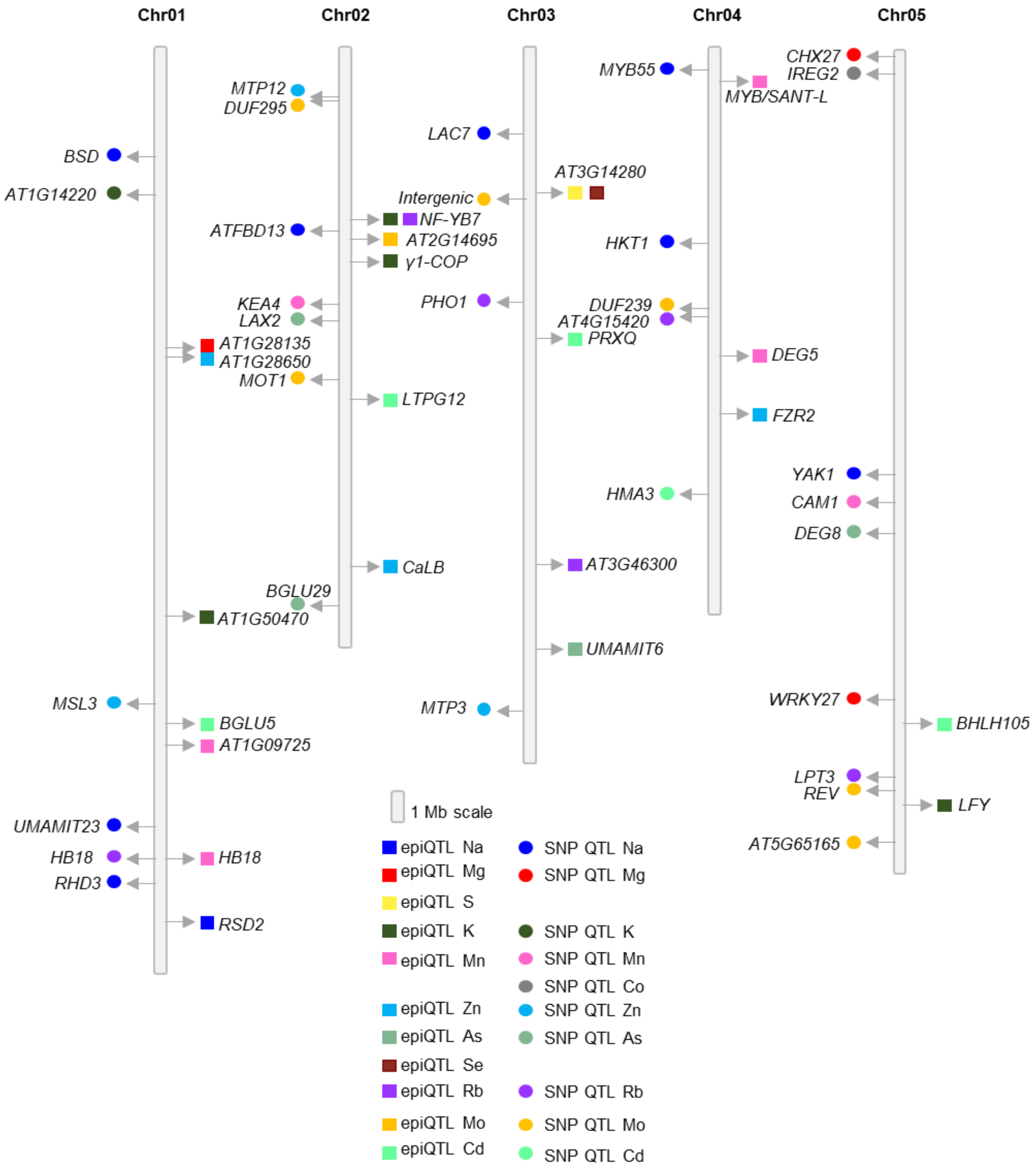


**Figure S28. Epigenetic and genetic QTLs associated with *Arabidopsis* leaf ionome are largely independent**. *Arabidopsis* physical map showing the positions of epigenetic and genetic QTLs underlying the levels of sodium (Na), magnesium (Mg), sulfur (S), potassium (K), manganese (Mn), cobalt (Co), zinc (Zn), arsenic (As), selenium (Se), rubidium (Rb), molybdenum (Mo), and cadmium (Cd) phenotypes. Five chromosomes are shown in Mb scale. Positions of epigenetic QTLs are shown with squares to the right of chromosomes, and circles on the left depict positions of genetic QTLs.

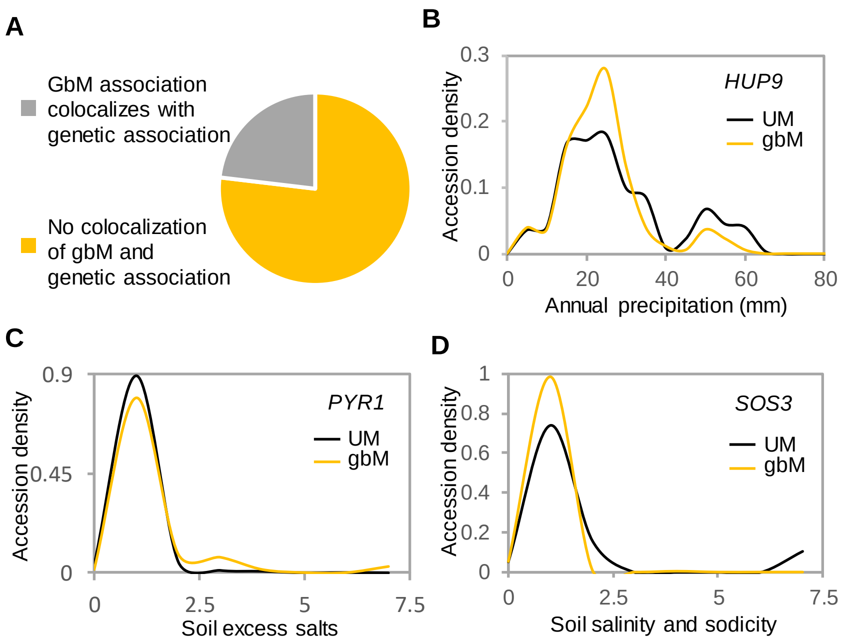


**Figure S29. Associations between gbM variation and the environment.** (**A**) Frequency of associations between gbM and environmental data colocalizing with genetic associations. (**B**-**D**) Associations between gbM and environmental data for *HUP9* (**B**), *PYR1* (**C**), and *SOS3* (**D**).
